# A novel role for Myosin-Va in mitochondrial fission

**DOI:** 10.1101/655803

**Authors:** Jackeline S Araujo, Rui M P Silva-Junior, Tong Zhang, Cara R Schiavon, Qian Chu, Melissa Wu, Carmen L S Pontes, Anderson O Souza, Luciane C Alberici, Aline M Santos, Sadie Bartholomew Ingle, Alan Saghatelian, James Spudich, Uri Manor, Enilza M Espreafico

**Affiliations:** Department of Cell and Molecular Biology, Faculty of Medicine of Ribeirão Preto, University of São Paulo, Ribeirão Preto, São Paulo, Brazil; Center for Cell-Based Therapy CEPID/FAPESP, Ribeirão Preto, São Paulo, Brazil; Department of Physics and Chemistry, Faculty of Pharmaceutical Sciences of Ribeirão Preto, University of São Paulo, Ribeirão Preto, São Paulo, Brazil; Department of Structural and Functional Biology, Institute of Biology, University of Campinas - UNICAMP, Campinas, Sao Paulo, Brazil; Waitt Advanced Biophotonics Center, The Salk Institute for Biological Studies, La Jolla, CA 92037, USA; Clayton Foundation Laboratories for Peptide Biology, The Salk Institute for Biological Studies, La Jolla, CA 92037, USA; Department of Biochemistry, Stanford University School of Medicine, Stanford, United States

**Author notes:** Co-corresponding authors: Uri Manor –, Enilza Espreafico.

**Keywords:** Myosin-Va, mitochondrial dynamics, actin cytoskeleton, Warburg effect, melanoma

## Abstract

In cancer cells metabolic changes and mitochondrial morphology are coupled. It is known that the cytoskeleton and molecular motors are directly involved in regulating mitochondrial morphology. Here we show that myosin-Va, an actin-based molecular motor, is required for the malignant properties of melanoma cells and localizes to mitochondria in these cells. Knockdown of myosin-Va increases cellular respiration rates and ROS production and decreases glucose uptake and lactate secretion. In addition, knockdown of myosin-Va results in reduced mitochondrial fission and correspondingly elongated mitochondria. We show that myosin-Va interacts with the mitochondrial outer membrane protein Spire1C, an actin-regulatory protein implicated in mitochondrial fission, and that Spire1C recruits myosin-Va to mitochondria. Finally, we show that during mitochondrial fission myosin-Va localization to mitochondria increases, and that myosin-Va localizes to mitochondrial fission sites immediately adjacent to Drp1 punctae. We conclude that myosin-Va facilitates mitochondrial fission. These data implicate myosin-Va as a target for the Warburg effect in melanoma cells.

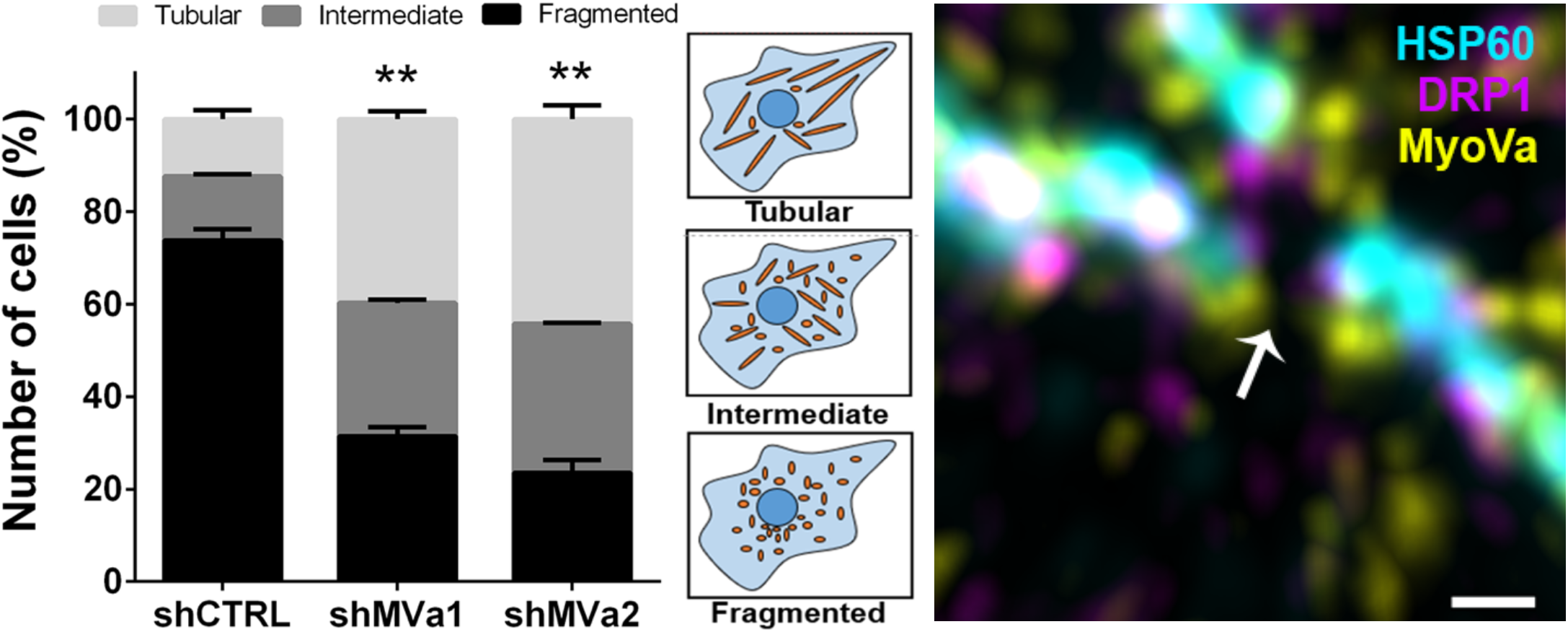

## INTRODUCTION

The reprogramming of cellular metabolism in favor of aerobic glycolysis is a hallmark of cancer, called the Warburg effect. In melanoma cells this is exemplified by the oncogenic mutation of BRAF^V600E^, which, among other changes, exacerbates mitochondrial fragmentation^4,5^. Mitochondrial fusion and fission are coordinated by dynamin-related GTPase proteins, the endoplasmic reticulum (ER), and the actin cytoskeleton. Fission is mediated by DRP1^6^ and DRP1 protein adaptors, such as Fis1^7,8^. Fusion is mainly promoted by Mfn1 and Mfn2, acting on the outer mitochondrial membrane (OMM), and Opa1, on the inner mitochondrial membrane (IMM)^9^. However, growing evidence indicates that alterations of many protein players and pathways act to trigger mitochondrial fragmentation as an adaptive change in cancer cells, in a process not completely understood.

Myosin-Va plays a central role in the transport of melanosomes in melanocytes, but a widespread subcellular distribution and other lines of evidence have paved the view that it has a much wider function. For instance, 10 11 12 myosin-Va is concentrated in microtubule-rich domains^10,11^, associates with nuclear speckles^12^, and with the majority of cytoplasmic organelles, including the ER, Golgi, endosomes, peroxisomes, and mitochondria^13,14^.

Myosin-Va is also upregulated in metastatic melanoma cells and required for manifestation of malignant properties, such as anchorage independent survival, migration, and invasion^15^. In addition, by sequestering the pro-apoptotic factor Bmf^16^, myosin-Va modulates apoptosis triggered by mitochondrial outer membrane permeabilization (MOMP). Interfering in this pathway by overexpressing a small fragment of the myosin-Va tail, which encompasses the DYNLL2 binding site, induces apoptosis in melanoma cells^1^. The myosin-Va ortholog, Myo2p, is required for mitochondrial transport and mitochondrial inheritance in daughter cells in the budding yeast *Saccharomyces cerevisiae*^17^. In mammalian cells, myosin-Va knockdown has also been shown to increase mitochondrial size in neurons^18^, as well as mediate energy metabolism via GLUT4 translocation in adipocytes and muscle cells^19–21^.

The processes of mitochondrial fusion and fission are intertwined with the nutritional and metabolic status of the cell. Many layers of structural and regulatory components influence the fusion and fission effectors, including the cytoskeleton and molecular motors^2,22–31^. Several studies have shown that myosin-V motors interact with and are recruited by the actin nucleating proteins Spire to Rab11-decorated membrane vesicles^32–35^. Recently, the Spire1C isoform that is anchored to the OMM was shown to modulate mitochondrial fission^2^. In view of this and other lines of evidence mentioned above, we postulated that myosin-Va may play a role in mitochondrial dynamics. To investigate this, we examined the effects of depleting myosin-Va on mitochondrial function, morphology and structural integrity in the A375 BRAF^V600E^ mutant metastatic melanoma cell line. We found that myosin-Va depletion in A375 BRAF^V600E^ cells causes increased ROS production, lower mtDNA levels, and mitochondrial elongation. We also show that Spire1C interacts with the myosin-Va tail, and that increased expression of Spire1C recruits myosin-Va to the mitochondrial outer membrane. In addition, we found that myosin-Va localizes to mitochondrial fission sites adjacent to Drp1. Our results reveal a role for myosin-Va in mitochondrial fission, further increasing our understanding of the molecular crosstalk between myosin Va, the actin cytoskeleton, and mitochondrial dynamics.

## RESULTS

### Myosin-Va knockdown increases melanoma cell OXPHOS activity while leading to decreased clonogenic ability

To investigate whether myosin-Va plays a role in the mitochondrial function and melanoma cell metabolism, we knocked down MYO5A in A375 BRAF^V600E^ mutant cells, using two short hairpin RNAs to human myosin-Va (shMVa1, shMVa2) with 60–85% efficiency (Figure 1A and 1B), and a non-targeted shRNA as control (shCTRL). First, we analyzed oxygen consumption of the cells, and found that myosin-Va knockdown cells displayed increased oxygen consumption rates on basal, State III (phosphorylation state – OXPHOS capacity), proton leak (L) and maximal respiratory capacity (E – mitochondrial electron transfer system) states (Figure 1C-G)^36^. We also analyzed the clonogenic ability of these cells, and we found that myosin-Va knockdown cells had lower clonogenic capacity compared to control cells (Figure 1H). Accordingly, myosin-Va knockdown cells showed lower uptake of glucose, lower lactate secretion, and higher ROS production than control cells (Figure 1I-K). To understand if the alterations in mitochondrial function affected the expression of genes and their respective protein products involved in mitochondrial biogenesis and dynamics, we performed mRNA and protein expression analysis. DRP1 mRNA and Drp1 proteins, both total Drp1 and p(S616)Drp1, were unchanged (Figure S1A-C). Mfn1 and Mfn2 mRNA levels were slightly decreased, but no alteration in protein levels were observed (Figure Figure S1A-C).

**Figure 1.**
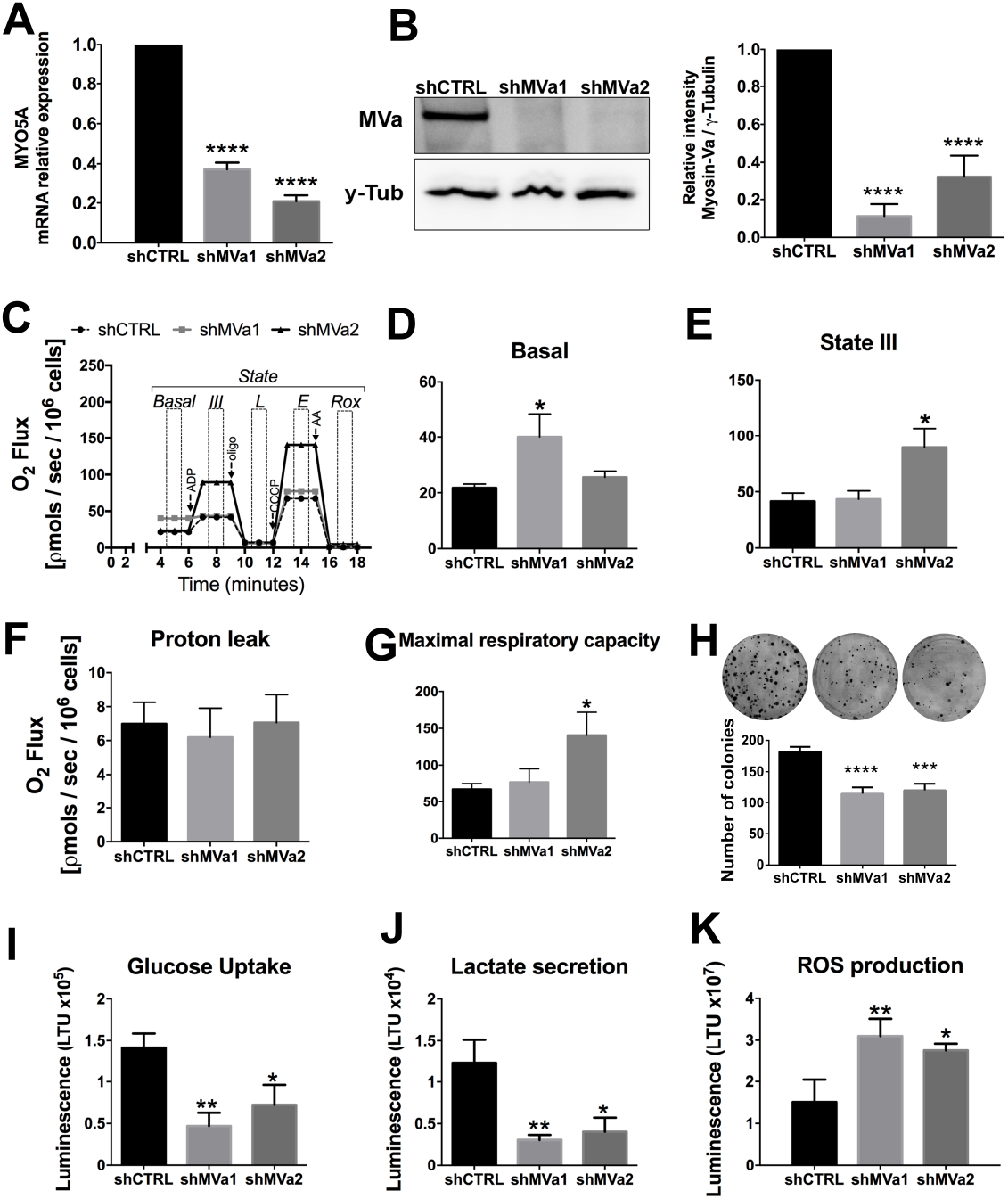
Knockdown of myosin-Va in A375 cells increases oxidative phosphorylation, increases ROS production, and lowers the glucose uptake levels and lactate secretion levels. (A) MYO5A mRNA expression in A375 cells stably transduced with two different shRNAs (shMVa1 and shMVa2). The values plotted are relative to shCTRL. The GAPDH gene was used as housekeeping with the formula 2-ΔΔCt to normalize the expression level. Data are mean ±SD (n=3), **** P< 0.0001. (B) Western blots containing total lysate of transduced A375 cells (line 1-shCTRL; line 2-shMVa1 and line 3-shMVa2) probed with affinity-purified rabbit polyclonal antibody raised to bacterially-expressed GST-myosin-Va medial tail (MVa) and a commercially-available monoclonal antibody to γ-Tubulin (γ-Tub) used as protein load normalizer. The densitometry quantification of myosin-Va relative to γ-Tubulin is shown below. Data are mean ±SD (n=3), **** P< 0.0001. (C) Mitochondrial respiratory parameters of A375 mysoin-Va knockdown cells (shMVa1 and shMVa2) and control (shCTRL) – Representative oxygraph traces of showing respiratory activity of digitonin-permeabilized cells in the presence of substrate for complex I (pyruvate, glutamate and malate). Arrows indicate the additions of 240 μM ADP, 0.5 μg/mL oligomycin (oligo), 0.25 μM CCCP and 0.5 μg/mL antimycin A (AA) used to determine the States of Basal, State III or phosphorylating (P), proton leak (L), maximal respiratory capacity (E) and residual (Rox). (D) Respiratory rates in States Basal, (E) State II, (F) Proton leak, and (G) maximal respiratory capacity. The values represent the mean ± S.E.M from five independent experiments (one 100cm dish plate was used for each experiment, approximately 1×106 cells). One-way ANOVA test followed by post hoc Turkey test was used for data analysis. *P<0.05. (H) Clonogenic assay showing a representative figure of the colonies for A375 myosin-Va knockdown cells and control, and quantification of the number of colonies for each cell. The values represent the mean ± S.D. from three independent experiments. One-way ANOVA test followed by post hoc Dunnet test was used for data analysis. ***P<0.0005; ****P<0.0001. (I) Glucose uptake quantification, (J) Lactate secretion quantification, and (K) ROS production quantification in A375 myosin-Va knockdown and control cells. The values represent the mean ± S.D from three independent experiments. One-way ANOVA test followed by post hoc Dunnet test was used for data analysis. *P<0.01; **P<0.001

OPA1 mRNA expression was increased, but no alteration of OPA1 protein levels were detected (Fig Figure S1A-C). We also observed an increase in the mRNA levels for PGC1α, a master regulator of mitochondrial biogenesis (Figure S1D). Additionally, myosin-Va knockdown cells had decreased levels of mtDNA (Figure S1E).

### Myosin-Va depletion reduces mitochondrial fission and leads to mitochondrial elongation

A375 BRAF^V600E^ mutant cells possess predominantly fragmented mitochondria, which makes them a useful model for understanding changes in mitochondrial dynamics. To investigate whether knockdown of myosin-Va alters mitochondria in these cells, we imaged myosin-Va knockdown (shMVa2) versus control (shCTRL) cells. We found that knockdown cells displayed longer mitochondria (Figure 2A). Over 40% of the knockdown cells showed elongated mitochondria, whereas only 12% of shCTRL cells showed elongated mitochondria. Consistently, the frequency of cells with fragmented mitochondria was above 70% in shCTRL cells and only about 30% in the myosin-Va-knockdown cells (Figure 2B). Finally, using live timelapse imaging of shMVa2 vs shCTRL cells stained with mitotracker, we found that myosin-Va knockdown cells displayed a reduced number of mitochondrial fission events (Figure 2C and Suppl. Movie 1+2).

**Figure 2.**
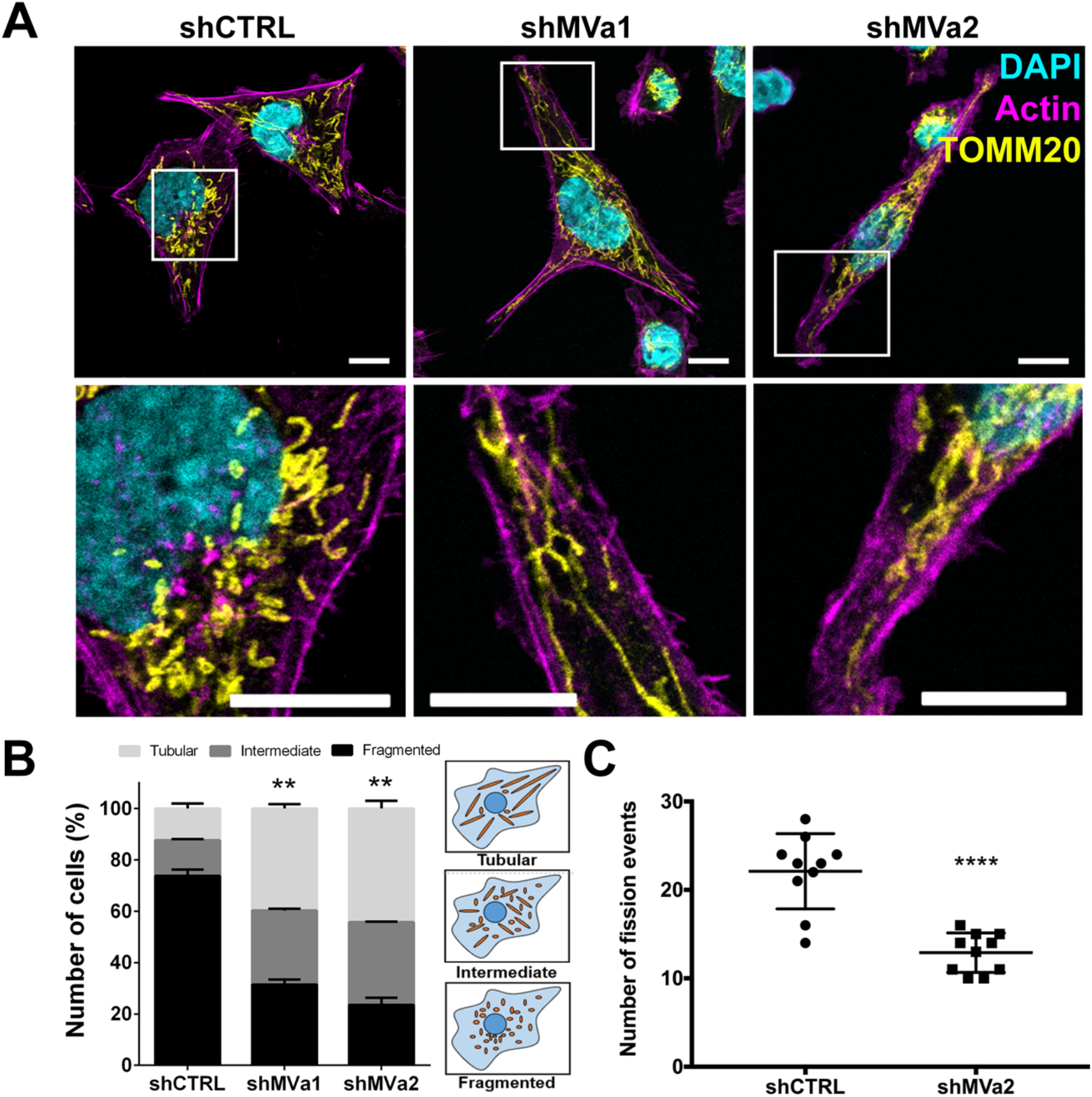
Myosin-Va knockdown promote changes in the mitochondrial network distribution. **(A)** Confocal microscopy images in A375 myosin-Va knockdown cells (shMVa1 and shMVa2) and our control (shCTRL) showing the mitochondrial network immunostained with anti-TOMM20 (yellow) antibody, the F-actin were stained with phalloidin-Alexa-594 (magenta) and the nuclei with DAPI (cian). **(B)** The mitochondrial morphology was quantified by classifying the appearance of mitochondria in A375 myosin-Va knockdown cells; at least 150 cells were counted on three different slides. **(C)** Number of fission events of myosin-Va knockdown cells (shMVa2) compared to control cells (shCTRL), representative movies are shown in supplementary material. Data was analyzed using Unpaired t test, ****P<0.0001 (n=10 cells for each condition). Scale Bar: 10μm and 1μm in the insets. Data are mean ±SD. **P < 0.001.

To better understand the morphological changes in knockdown cells, we employed transmission electron microscopy (TEM). TEM imaging revealed the presence of enlarged mitochondria in the myosin-Va knockdown cells in comparison to shCTRL cells (Figure 3A). Quantification of area (Figure 3B) and perimeter (Figure 3C) confirmed mitochondrial enlargement in knockdown cells. Interestingly, the length of contact regions between endoplasmic reticulum and the mitochondrial surface relative to the total length of the mitochondrial perimeter also decreased in the knockdown cells (Figure 3D).

**Figure 3.**
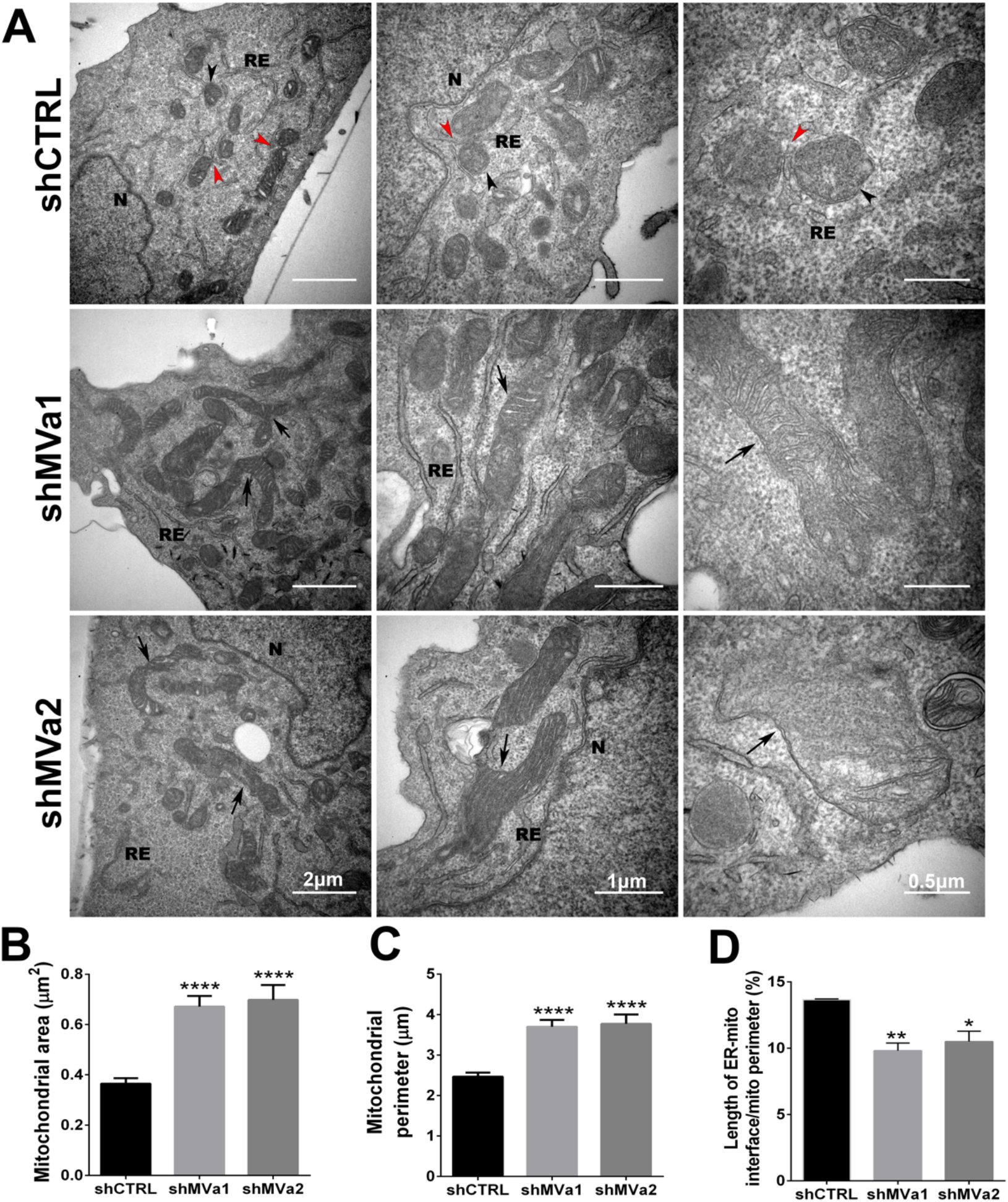
Mitochondrial ultrastructure is affected by MYO5A knockdown in A375 cell line. **(A)** Representative electron micrographs of A375 myosin-Va knockdown cells (shMVa1 and shMVa2) and control (shCTRL) showing the mitochondrial ultrastructure at different magnifications (RE – Endoplasmic reticulum; N – Nucleus). The red arrowheads show the contact sites between the RE and the mitochondrial membrane, the black arrowheads indicate fragmented mitochondria (shCTRL); and the black arrows show enlarged mitochondria (shMVa1 and shMVa2). **(B)** Quantification of mitochondrial area (μm^2^), **(C)** quantification of mitochondrial perimeter (μm), and **(D)** percentage of the length of ER-mitochondria interface, normalized by mitochondrial perimeter. Data are mean ±SD (at least 90 mitochondria per condition). *P<0.01; **P<0.001; ****P<0.0001. Scale Bar = 2 μm, 1 μm, 0.5 μm with magnification of 20x (left panels); with 40x (middle panels) and 80x (right panels).

### Myosin-Va localizes to discrete clusters on the mitochondrial surface

To assess whether myosin-Va plays a direct or indirect role in mitochondrial fission, we analyzed myosin-Va localization relative to the mitochondrial network. To this end, we used two different antibodies to the myosin-Va medial tail for co-staining of myosin-Va and mitochondria. Using a rat anti-myosin-Va and anti-TOMM20, we found that a fraction of the myosin-Va staining colocalized with the mitochondrial network in every cell (Pearson’s correlation = 0.244±0.05, Figure S2A) and, by examining the orthogonal projection, we observed that myosin-Va clusters often surround the mitochondrial network (Figure S2B and S2C). This colocalization was not due to randomly colocalized pixels, since when we rotated the image of myosin-Va staining 180° relative to the TOMM20 image and reanalyzed^37^, the Pearson’s correlation coefficient was then as low as 0.005±0.06. Similar colocalization was obtained using rabbit anti-myosin-Va and anti-Cytc co-staining (Figure 4A). Also, structured illumination microscopy showed that myosin-Va localizes to discrete clusters associated with TÛMM20 stained mitochondria (Figure 4B). The degree of colocalization seen here is consistent with the one previously observed by our group in melanoma cells using immunogold staining^38^.

**Figure 4.**
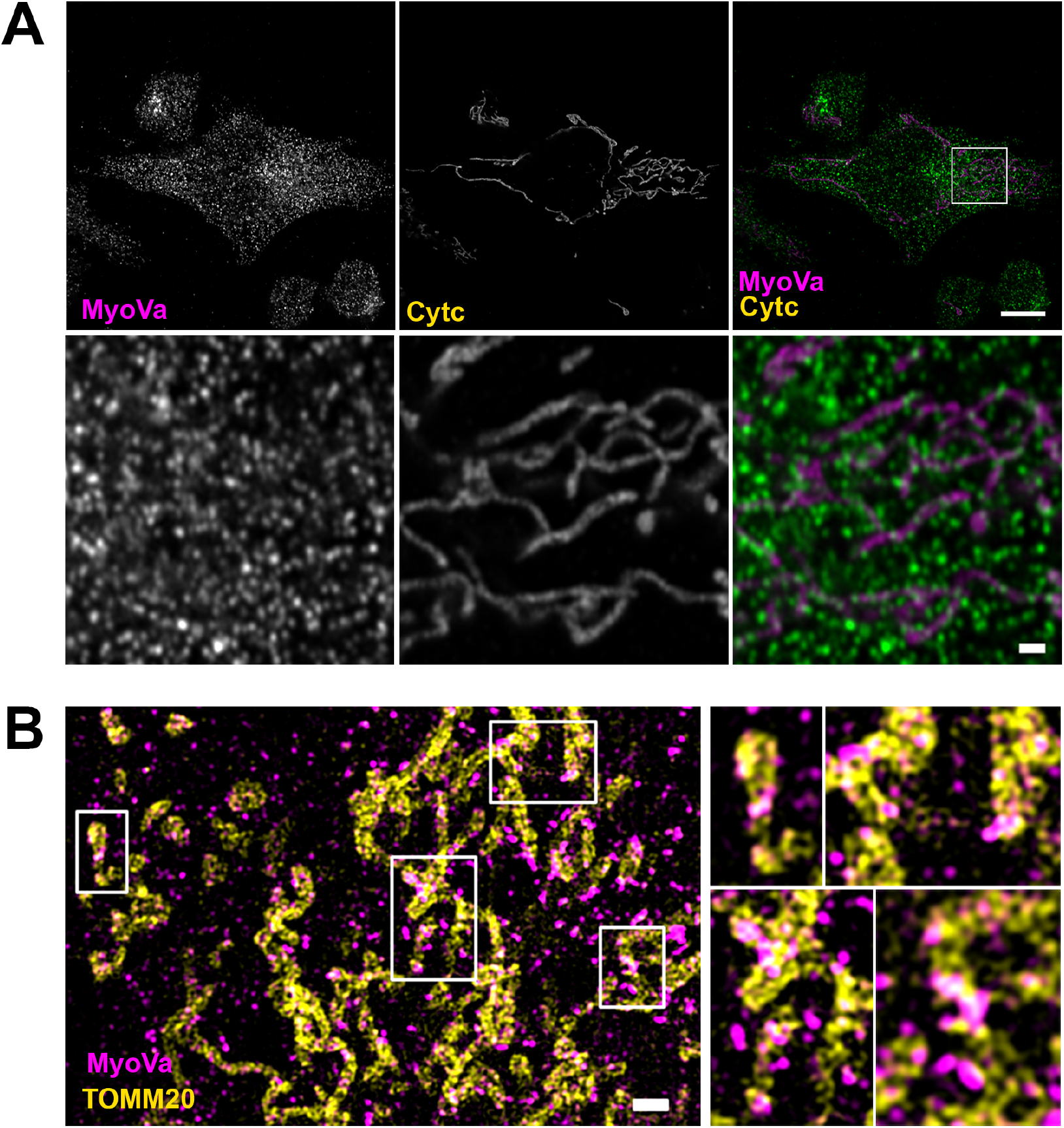
Myosin-Va colocalizes with mitochondrial network in A375 cell line. **(A)** High resolution Airyscan time lapse images of A375 cells showing the localization of myosin-Va, stained with a rabbit polyclonal antibody, and the mitochondrial network mitochondrial network immunostained with anticytochrome c antibody. Scale bar: 10μm and 1μm in the insets. **(B)** Superresolution images of A375 cells showing the immunolocalization of myosin-Va (red) and TOMM20. Images were obtained using an ELYRA S.1 microscope (Carl Zeiss Microimaging). Scale bar 1μm.

### Myosin-Va interacts with and is recruited by Spire1C to mitochondria

We performed affinity chromatography with the medial-globular tail domain of myosin-Va (MVa-MGT) and the cargo binding domain of myosin-VI to identify potential novel binding partners for those proteins. Samples were analyzed using mass spectrometry, and the candidate binding proteins were sorted according to^39^. When we ranked the candidate proteins, spire1 was a top hit, with clear specificity for myosin-Va over myosin-VI (Suppl. Table 1). This result is supported by previous demonstrations that myosin-V motors bind other splice isoforms of the Spire family^32–35^. To validate our result, we expressed myc-tagged Spire1C in HEK293 cells and performed mass spectrometry on myc-Spire immunoprecipitates. We found considerable enrichment of myosin-Va in myc-Spire1C immunoprecipitates compared to control (pcDNA empty) and, interestingly, we found that nearly double the amount of myosin-Va was pulled down with the mitochondrial isoform of Spire1 (Spire1C) compared to the non-mitochondrial isoform (Spire1ΔC) (Figure 5B). To further test whether there is an interaction between Spire1C and the myosin-Va tail, we used a rabbit reticulocyte system for *in vitro* translation of myc-tagged Spire1C and a myosin-Va tail fragment containing the medial and globular domains. Using this assay, we showed that Spire 1C co-purifies with the myosin-Va tail (Figure 5C).

**Figure 5.**
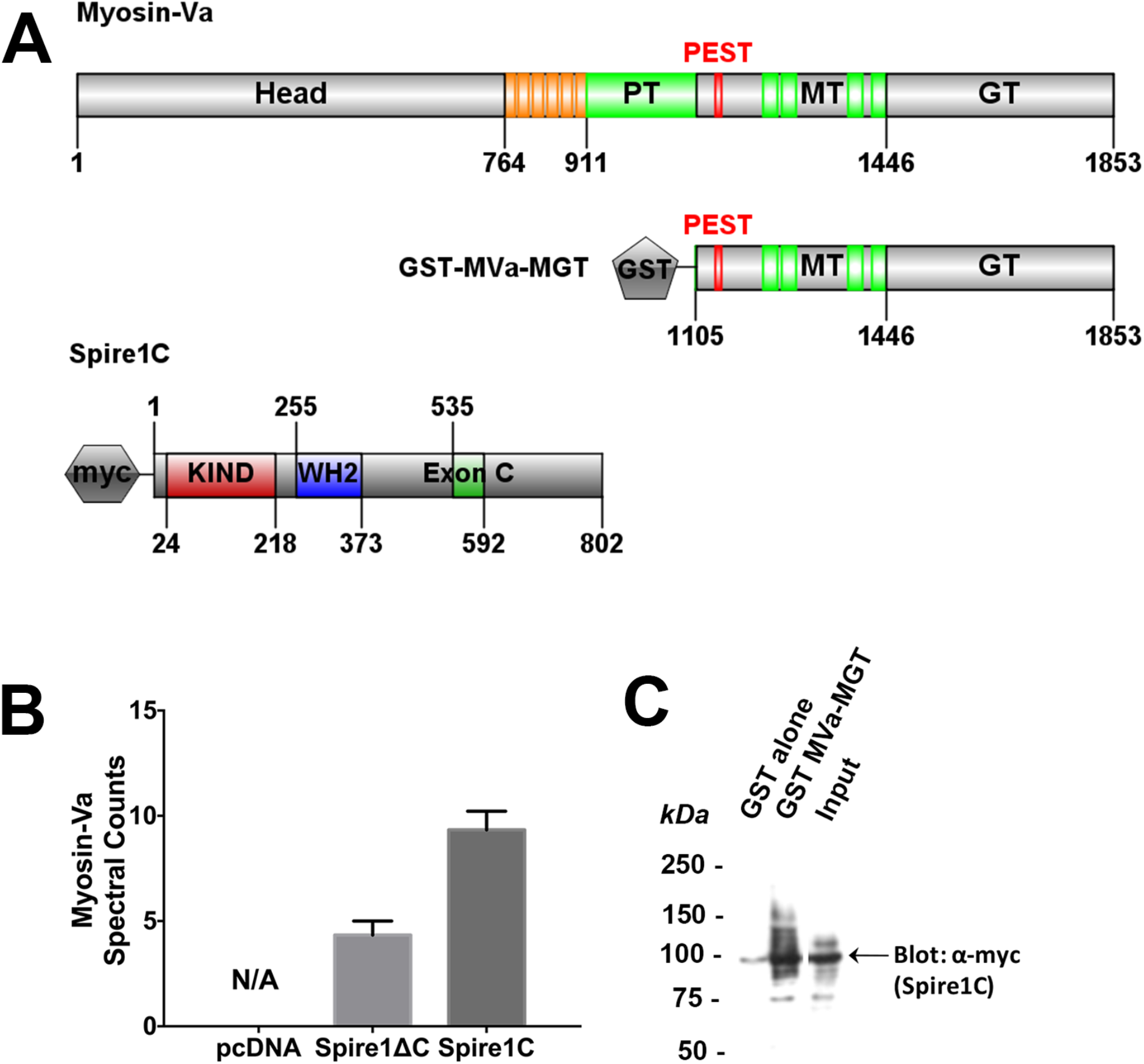
Myosin-Va interacts with Spire1C. **(A)** Schematic drawing representing the primary structure of the myosin-Va heavy chain. The position of the structural sub-domains are shown in: Head; Neck (IQ-motifs); PT, proximal tail; MT, medial tail; GT, globular tail; MGT, medial globular tail. Figure adapted from Izidoro-Toledo et al., 2013^1^; GST-MVa-MGT structure is shown below, and it was used for the *in vitro* interaction with Spire1 proteins. Full length Spire1C domain structure is shown, with the subdomains also represented, KIND, WH2 and ExonC. Figure adapted from Manor et al., 2015^2^. IBS freely available online was used for the schemes^3^. **(B)** Semi-quantitative proteomics by spectral counting demonstrates enrichment of Myosin-Va by Spire1C full length. Error bars represent the standard deviation of three biological triplicates. **(C)** In vitro translated Spire1C interacts with purified GST-tagged myosin-Va globular tail domain, but not GST alone.

To test the hypothesis that myosin-Va is recruited to the OMM by Spire1C via interaction with the myosin-Va tail domain, we overexpressed myc-Spire1C and GFP-tagged chicken myosin-Va tail fragments (full tail; medial-globular tail; globular tail) in mammalian A375 cells (Figure 6). We found that all GFP-tagged myosin-Va tail constructs used (Figure 6), but not the GFP tag alone (Figure S3A), localizes to mitochondria much more strongly in cells overexpressing myc-Spire1C than in control cells (Figure 6; Figure S3B-E). These results support the hypothesis that Spire1C recruits myosin-Va to the OMM by interacting with the myosin-Va tail domain.

**Figure 6.**
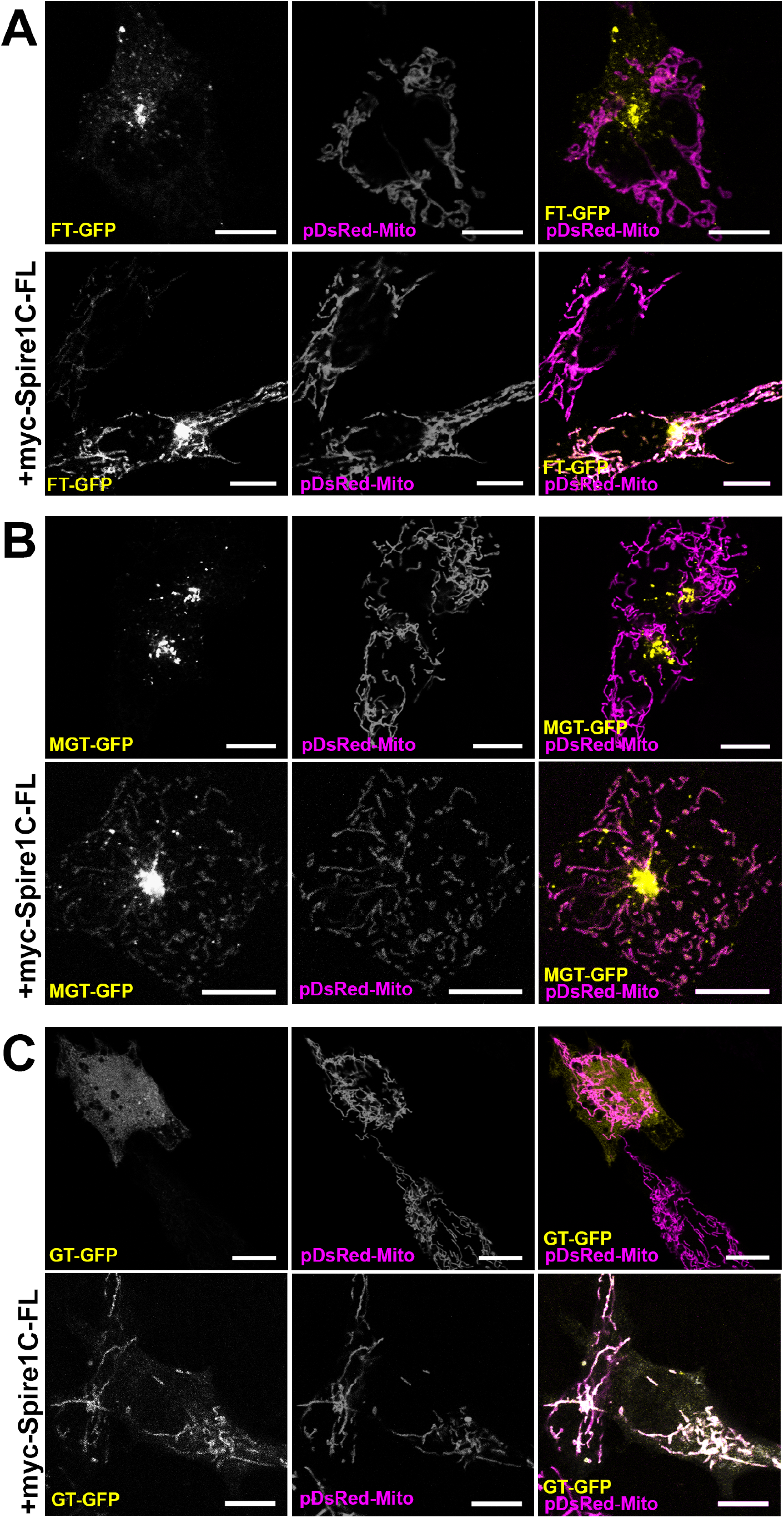
Myosin-Va tail is recruited to mitochondria by Spire1C. **(A)** FT-GFP (chicken brain myosin-Va full tail); **(B)** MGT-GFP (chicken brain myosin-Va medial-globular tail); **(C)** GT-GFP (chicken brain myosin-Va globular tail) expression along with pDsRed-Mito in A375 cells, as control for normal localization of FT-GFP; MGT-GFP or GT-GFP, and below the combined expression with myc-Spire1C-FL showing FT-GFP; MGT-GFP or GT-GFP recruitment to the mitochondrial network. Scale bar 10μm.

### Myosin-Va is located at mitochondrial fission sites along with Drp1

To better understand the potential role of myosin-Va in mitochondrial fission, we co-expressed the full length GFP-myosin-Va with a mitochondrial marker in U2OS cells and used high resolution Airyscan timelapse imaging to monitor mitochondrial dynamics (Figure 7, Suppl. Movie 3 and Movie 4). We found that 58% of the fission events counted had myosin-Va accumulation (Figure 7C), often at the tips of “daughter” mitochondria (Figure 7A and 7B). We also induced fission by treating the cells with ionomycin and observed a significant increase in fission events containing myosin-Va (in ~74% of fission events counted) (Figure 7B and 7C). To confirm the role of endogenous myosin-Va in the fission events, we co-stained U2OS cells with anti-myosin-Va, anti-Drp1, and anti-HSP60. We observed a significant fraction of myosin-Va localized to mitochondrial constriction sites, often adjacent to Drp1 at the ends of “daughter” mitochondria, suggesting that while these two proteins may not directly interact, they may both be localizing to active fission sites (Figure 8A). To further test the putative role of myosin-Va in fission, we induced fission by treating cells with ionomycin, then stained for myosin-Va, Drp1, and HSP60. As expected, Drp1 localization to mitochondria significantly increased by ~1.4x (from 43.5% to 60.6%) in cells treated with ionomycin (Figure 8C). In support of our hypothesis that myosin-Va plays a role in mitochondrial fission, myosin-Va localization to mitochondria increased by ~1.3x (from 37.3% to 48.3%) in ionomycin treated cells (Figure 8D). High magnification views of mitochondrial constriction sites revealed a high frequency of myosin-Va and Drp1 puncta localized at fission sites (Figure 8B).

**Figure 7.**
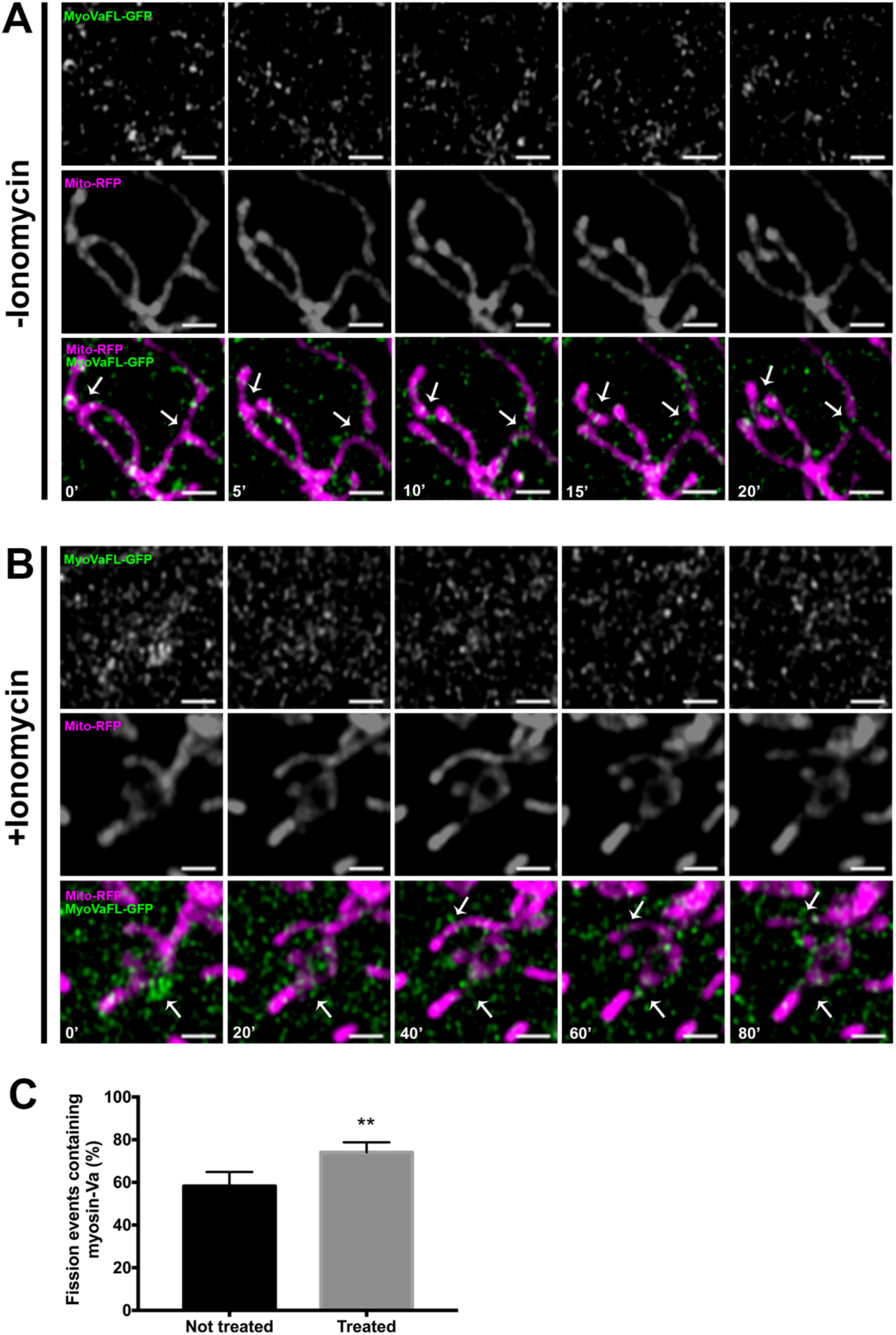
Myosin-Va localizes at mitochondrial fission sites. **(A)** High resolution Airscan time lapse images of U20S cells showing the localization of MyoVaFL-GFP (full length human myosin-Va) with a mitochondrial marker (Mito-RFP) in a control condition, and **(B)** treated with ionomycin. **(C)** Quantification of fission events that myosin-Va was present, either in the constriction of the mitochondria and in the tips of newly formed mitochondria. Fission events were counted from at least 5 timelapses for each condition. Unpaired t test was used for the analysis, data are mean ± SD, **P < 0.001. Scale Bar: lμm

To further characterize whether myosin-Va knockdown affects the localization of Drp1 and pDrp1 (pS616) to the mitochondria in A375 cells, we co-stained anti-Drpl and anti-pDrp1 (pS616) with anti-Cytc, as a mitochondrial marker. We observed that Drp1 (Figure 8E and 8F) and pDrp1 (pS616) (Figure 8G and 8H) localization to the mitochondrial network was decreased in myosin-Va knockdown cells compared to control cells (shCTRL). Since DRP1 gene and protein (Drp1 and pDrp1 (S616)) expression were not affected by the knockdown of myosin-Va in A375 cells (Fig S1A-C), we presume that the decreased mitochondrial localization of Drp1 is due to diminished recruitment.

**Figure 8.**
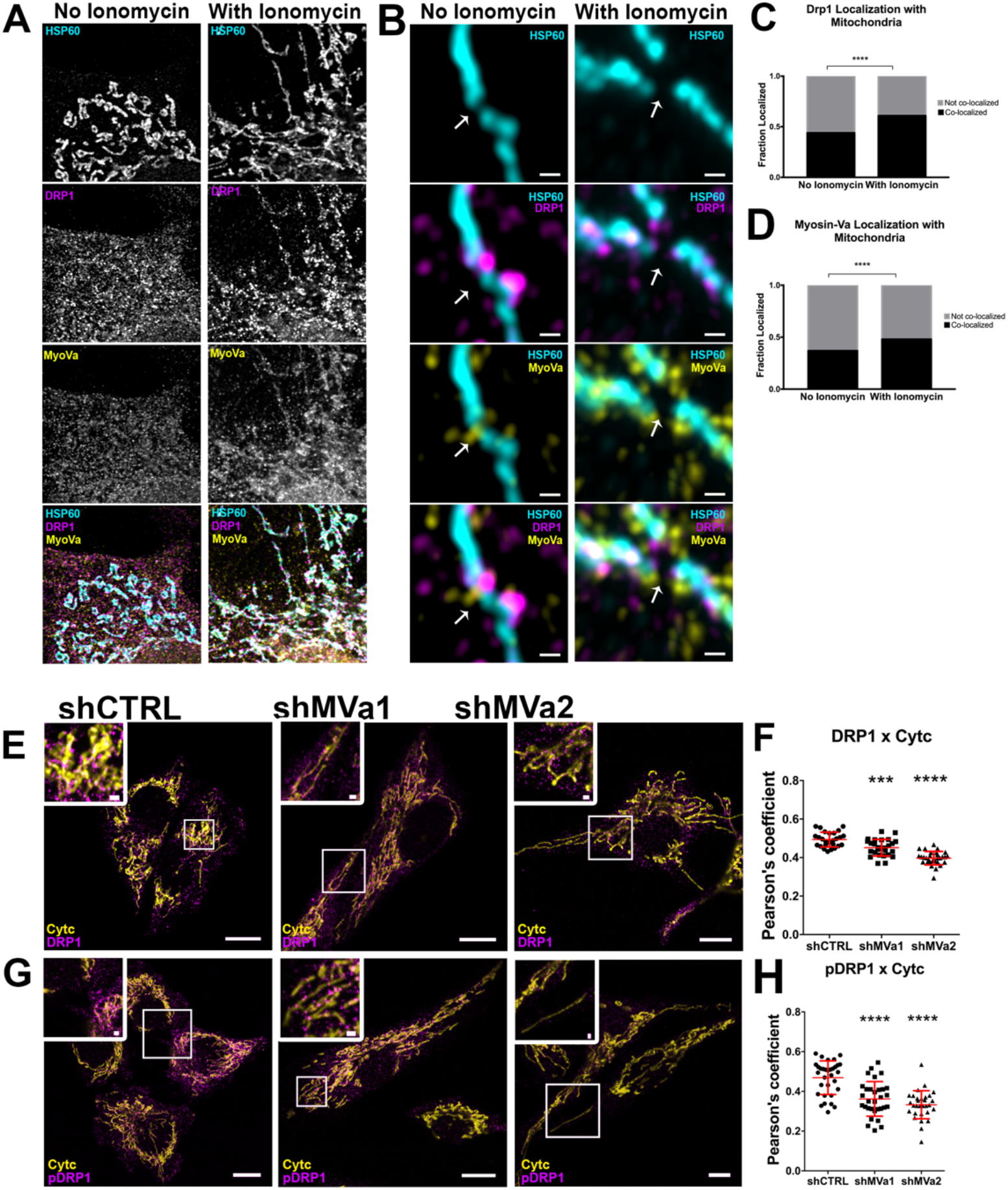
Myosin-Va coexists with DRP1 at fission sites and mediates DRP1 localization to the mitochondria. **(A)** High resolution Airscan images of U2OS cells showing the immunostaining of the mitochondrial network with anti-Hsp60 in cyan, the endogenous myosin-Va in yellow, and endogenous Drp1 in magenta, prior to ionomycin treatment (left panels), and after ionomycin treatment (right panels). **(B)** High magnification views of mitochondrial constriction sites of U2OS cells showing the immunostaining of the mitochondrial network with anti-Hsp60 in blue, the endogenous myosin-Va in green, and endogenous Drp1 in red, prior to ionomycin treatment (left panels), and after ionomycin treatment (right panels). Scale bar: 2μm. The graphs are representative of **(C)** Drp1 colocalization to mitochondria and of **(D)** myosin-Va colocalization to mitochondria with and without 4μM of ionomycin. Data was analyzed using two-wayANOVA ****P<0.0001. Confocal images in A375 myosin-Va knockdown cells (shMVa1 and shMVa2) and control (shCTRL) showing the mitochondrial network immunostained with anti-cytochrome c antibody, and antibodies against **(E)** DRP1 and **(G)** pDRP1(S616). **(F, H)** Pearson’s correlation coefficient showing the colocalization of DRP1 and pDRP1(S616), respectively, to the mitochondrial network. The data compare myosin knockdown cells shMVa1 (n=27 or 31 cells in the images of DRP1 and pDRP1(S616) respectively) and shMVa2 (n=30 or 30 cells in the images of DRP1 and pDRP1(S616) respectively, one replicate) to control cells shCTRL (n=26 or 32 cells in the images of DRP1 and pDRP1(S616) respectively, one replicate). Data was analyzed using one-way ANOVA, mean ± SD, **P < 0.001 and *** P< 0.0005, ****P<0.0001.

Interestingly, our affinity chromatography results also identified Drp1 as a candidate binding protein for myosin-Va, also with specificity for myosin-Va over myosin-VI (Suppl. Table 1), supporting the notion that myosin-Va, Spire1C, and Drp1 might function as a protein complex to mediate mitochondrial fission.

## DISCUSSION

Oncogenic transformation by expression of BRAF^V600E^, the most frequent oncogenic driver in melanoma, typically leads to metabolic reprogramming that favors aerobic glycolysis in detriment of OXPHOS, the Warburg effect^40,41^. This metabolic reprogramming of cancer cells is accompanied by mitochondrial fragmentation, which is likely linked to changes in their activity^42–44^. The human metastatic melanoma cell line A375 harbors a BRAF^V600E^ mutation and possesses predominantly fragmented mitochondria, which makes them a useful model for understanding mitochondrial dynamics in the context of cancer^5,40^. It is also worth underscoring that melanoma cells exhibit relatively high MYO5A gene expression^15^, thus provide a unique opportunity for studying MYO5A functions. Our data show that myosin-Va knockdown in A375 cells leads to higher rates of mitochondrial respiration, lower glucose consumption and lactate secretion, and higher ROS production. Such alterations are accompanied by reduced mitochondrial fission, culminating in mitochondrial elongation. These data support the conclusion that the morphological alterations triggered by loss of myosin-Va function perturb the Warburg effect caused by MAPK pathway activation in melanoma cells. It is now widely known that the Warburg effect favors biomass building of proliferative tissues^45^. Therefore, the reduced clonogenic ability of myosin-Va knockdown cells is consistent with our metabolic data and corroborates previous findings^1,15^ Exploring the possibility that myosin-Va functions as an effector molecule of the MAPK pathway to enhance mitochondrial fragmentation induced by oncogenic transformation is an intriguing possibility worthy of further exploration. However, the scope of the present study was directed toward understanding the mechanistic role of myosin-Va on the fission process itself, and a major finding was the demonstration that myosin-Va molecular motors interact with Spire1C through their tail domain and are recruited by Spire1C to the OMM. Moreover, myosin-Va localizes to mitochondrial fission sites, adjacent to Drp1, and its knockdown reduces Drp1 and pDrp1 localization to the mitochondria.

The mitochondrial elongation triggered by myosin-Va knockdown in A375 melanoma cells is consistent with previously reported increase in mitochondrial size following knockdown of myosin-Va in neurons^18^. The diminished number of fission events observed in A375 melanoma cells is consistent to a role of myosin-Va in mitochondrial fission, as it has been widely shown that a deficiency in the fission process results in mitochondrial elongation upon inactivation of key proteins of the fission machinery, such as myosin II^26^, IFN2^25,26^ and Drp1^46^. Intriguingly, likewise for myosin-Va knockdown, Drp1 knockdown has also been shown to result in diminished tumorigenesis and features, such as increased OXPHOS and low clonogenic ability^4,5,47^.

Our data show that myosin-Va interacts with Spire1C and overexpression of Spire1C increases myosin-Va localization to mitochondria. Recently, non-mitochondrial Spire splice isoforms have been shown to interact with myosin-V motors^32^. Since Spire1C is a modulator of mitochondrial fission^2^, the interaction between Spire1C and myosin-Va gives new insight towards the mechanism by which both myosin-Va and Spire1C regulate mitochondrial morphology. Whether myosin-Va transports any other fission modulating proteins to mitochondria remains an open question. But it should be noted that Rab11a, which interacts with both Spire and myosin-Va, has been suggested to play a role in mitochondrial fission. Finally, the induction of fission with ionomycin causes increased accumulation of myosin-Va on mitochondria, showing that myosin-Va localization to mitochondria is dynamically coupled to fission. Interestingly, following ionomycin stimuli, Drp1 accumulation at fission sites is mirrored by myosin-Va co-accumulation. Moreover, myosin-Va knockdown in A375 cells, resulting in mitochondrial elongation, is associated with a decrease in the MAPK activated form of Drp1 (Drp1-pS616) on the mitochondrial surface. Taken together, these findings reinforce a role for myosin-Va in the Drp1-mediated fission pathway.

Given the results in our ionomycin experiments, it is interesting to consider that myosin-Va activity is calcium-dependent^38,48,49^. Among other mechanisms, it has been shown that myosin-Va associates to the ER and binds to the ryanodine calcium receptor channels on the ER membrane, and also responds to ER-derived calcium by associating with vesicles and directing them to specific sites^50^. More recently, myosin-Va was shown to tether synaptic vesicles to the plasma membrane near Ca+^2^ influx sites and help to increase the retention of vesicles at the release site^51^. The decrease in the ER-mitochondrial contact surface, observed here in myosin-Va knockdown cells, is also consistent with a calcium triggered-role for myosin-Va at the ER-mitochondrial interface. Taken together, these findings suggest myosin-Va helps regulate mitochondrial fission by responding to local cytoplasmic calcium. This is a tempting hypothesis, as calcium can also increase actin polymerization and mitochondrial fission^52,53^.

Although we have not yet elucidated the biophysical mechanisms underlying mitochondrial elongation triggered by loss of myosin-Va, our data are indicative of a decreased number of fission events in the knockdown cells. The diminished extent of Drp1 localization to mitochondria in knockdown cells, and the localization of myosin-Va to fission sites in normal cells implicates myosin-Va as an enhancing factor for mitochondrial fission. Indeed, myosin-Va is an actin-based molecular motor, and as shown previously by multiple studies, actin and the ER are essential for Drp1-mediated mitochondrial fission^2,24–29,53,54^ Moreover, as Drp1 is as a candidate binding protein for myosin-Va, an interaction between those proteins could be a mechanistic explanation for the myosin-Va role in mitochondrial fission. In addition to Drp1, we postulate that other GTPase molecules are probably involved in myosin-Va mediated fission, given previous evidence that Rab7a and Rab11a can interact with and activate mitochondrial fission machinery^55,56^. Interestingly, Spire family proteins interact with both myosin-Va and Rab11a to regulate vesicle motility, suggesting a conserved molecular mechanism by which myosin-Va, Rab GTPase proteins, and actin regulatory proteins cooperate to regulate membrane-bound organelle dynamics. Furthermore, previous studies have shown a role for myosin-Va in regulating ER dynamics^57^, providing one more potential mechanistic link between myosin-Va activity and mitochondrial fission. That myosin-Va is a motor protein specialized for directed motility and transport, along with the localization of myosin-Va at the separating ends of daughter mitochondria, suggests that myosin-Va may play a role in moving newly separated mitochondria apart to complete the fission process and/or to avoid that they fuse back. Concluding, the results presented here prompt many questions to be pursued in future studies regarding the molecular partners of myosin-Va that are required to promote the fission process, and how its function is regulated, both in normal and cancer cells.

## MATERIALS AND METHODS

### Cell culture

Cell lines A-375 (ATCC^®^ CRL-1619™) and HEK-293T were cultured in Dulbecco’s Modified Eagle’s Medium (DMEM – Thermo Fisher Scientific, Waltham, MA, EUA) supplemented with fetal bovine serum (FBS) to a final concentration of 10% (v/v) and 100 mg/ml of streptomycin. The cultures were maintained in a cell incubator (Waltham, MA, USA) under 95% air and 5% CO2 at 37 °C. Cells were harvested and passaged for experiments with 0.25% trypsin and 0.025% EDTA, and viability was monitored by trypan blue exclusion.

### Immunoblotting

Cells were grown in 6-well plates, and after 24h were washed and lysed in lysis buffer (1% Triton-X100, 50 mM Tris ph 8.0, 150 mM NaCl, 0,1% SDS, 1:100 dilution of Sigma-Aldrich cocktail proteinase inhibitor, 20 mM NaF, 2 mM Na_3_VO_4_, 5 mM MgCl_2_ and 10mM ATP, pH 7.4). Lysates were passed through a 19-gauge syringe needle 10 times, and supernatants were collected after centrifugation at 11,000 *×g* at 4°C for 10 min and protein was quantification using the Bradford method (Bradford, 1976). 40 μg of total protein was loaded in a 10% polyacrylamide gel, then separated by SDS–PAGE. Immunoblotting was conducted on nitrocellulose membrane (Bio-Rad, Richmond, CA, USA) with respective primary antibodies and visualized by HRP-conjugated secondary antibodies by enhanced chemiluminescence (GE Life Sciences, Buckinghamshire, England).

### Lentivirus production and cell transduction

For lentiviral production, HEK-293T cells were cultured in suitable medium and transfected by calcium phosphate method with the following plasmids: 10 μg of pLKO.1 (Sigma-Aldrich, St Louis, MO, USA – SHCLNG-NM_000259), containing the shRNA cassette; 3.5 μg pVSV-G, 6.5 μg pΔ8.9, with culture medium with serum and without antibiotic. After sixteen hours, the transfection medium was changed to complete culture medium. Collection of the supernatant was performed on day 2 or 3 post-transfection. The supernatant was filtered through a 0.22 μm membrane and added to the cells of interest for the transduction. After 48 hours, the culture medium was changed to 10% DMEM medium containing puromycin (1 μg/ml) for selection of transduced cells. The sequences of the shRNA used here are in Table 1.

**Table 1.**
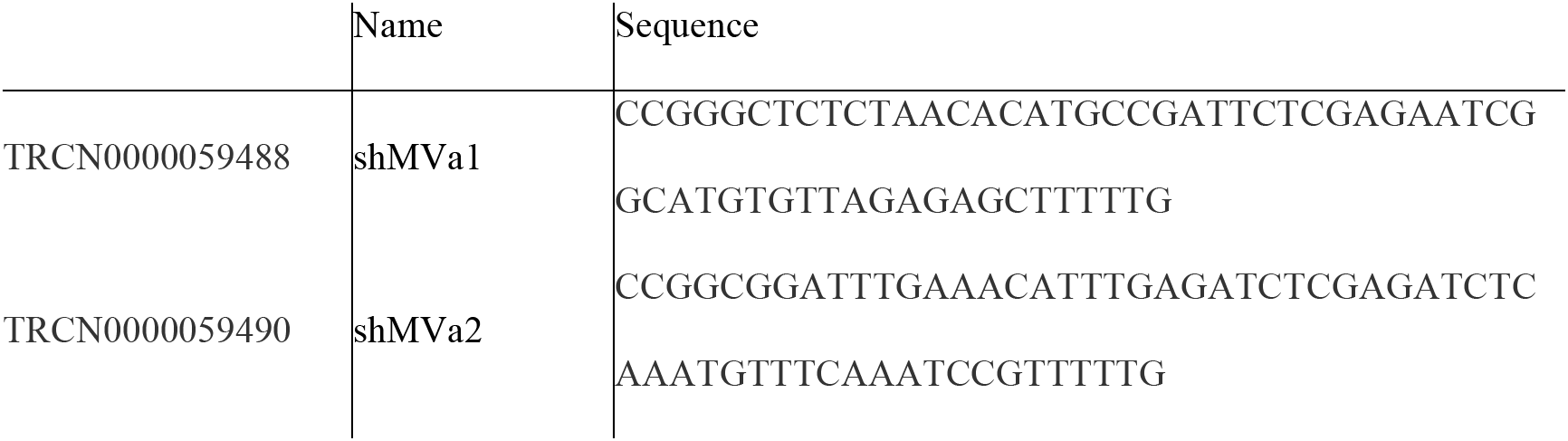
Sequences of the shRNAs used to knockdown MYO5A gene.

### DNA, RNA isolation and qRT-PCR

Genomic DNA (gDNA) was extracted using the ReliaPrep ™ gDNA Tissue Miniprep System kit (Promega) according to the manufacturer’s specifications. gDNA was amplified using primers according to a method previously described^58^. The sequences analyzed are in Table 1. Relative quantification of mtDNA expression was performed using the 2-ΔΔCt method, and normalized with nDNA.

Total RNA was extracted from the cells using the TRIZOL reagent (Invitrogen, Carlsbad, CA, USA) according to the manufacturer’s specifications. The RNA pellets were suspended in DEPC water and the samples were quantified at 260/280 nm in a spectrophotometer (NanoVue – GE Life Sciences, Buckinghamshire, England).

For the quantitative RT-PCR, all RNA samples (1 μg) were converted to cDNA using the High-Capacity cDNA Reverse Transcription Kit (Applied Biosystems). The reaction was incubated at 25 °C for 5 minutes, 42 °C for 30 minutes and the reverse transcriptase was inactivated by heating at 85 °C for 5 minutes. For amplifications, SYBR Green Power Mix reagent (Applied Biosystems) was used in reactions with a final volume of 12 μl containing 6 μl of SYBR Green Power Mix, 2 μl of a sense and antisense primer mixture (400 nM of each primer in the reaction) and 4 μl of cDNA. Amplification reactions and analysis were performed on an ABI PRISM 7500 Sequence Detection System (Applied Biosystems) using the ABI 7500 Real-Time PCR SDS 1.2 software (Applied Biosystems). Relative quantification of mRNA expression was done using the 2^-ΔΔCt^ method. The primers for the analyzed genes were designed flanking intron regions to avoid the amplification of unwanted products. GAPDH and TBP genes were used as reference genes. The sequences of the primers are shown in the table below (Table 2). The real-time PCR data were analyzed using the unpaired t-test, with P<0.05 considered statistically significant.

**Table 2.**
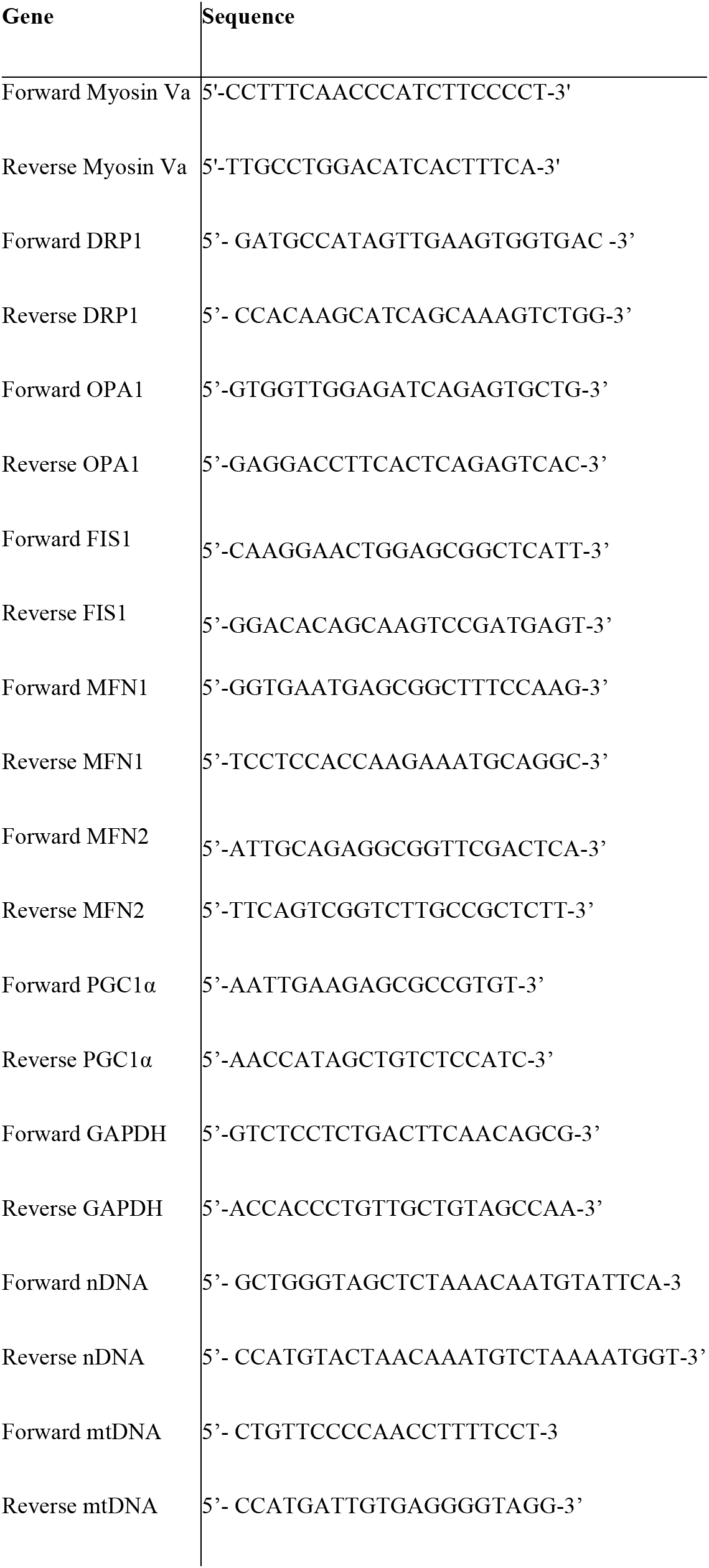
Sequences of the primers used for qRT-PCR.

### Exogenous protein overexpression

For exogenous protein overexpression assays, 5 × 10^4^ A375 cells were incubated overnight on 13 mm diameter cover slips (Knittel, Braunschweig, Germany) in 24-well plates (Corning incorporated Costar^®^). The cells were transfected using Lipofectamine 2000 (Thermo Fisher Scientific, Carlsbad, CA)^®^. Transfections were carried out in a final volume of 250μL medium following the manufactures’ instructions. After 48h of incubation, cells were washed with PBS, fixed with 4% *(v/v)* paraformaldehyde pH 7.4 for 15 min, washed once more with PBS and processed for immunofluorescence and data acquisition as described above. For these assays, cells were imaged using a Leica CTR 6000 microscope (Leica).

The following constructions were used: mCherry-DRP1 (AddGene Plasmid #49152); pDsRed-Mito (AddGene Plasmid #55838); full length Spire1C (myc-SpireFL)^2^; mouse myosin-Va-Full Length (MyoVaFL-GFP)^59^; chicken brain globular tail of myosin-Va (GT-GFP – residues 1423–1830); chicken brain medial tail (partial) and globular tail of myosin-Va (MGT-GFP – resiidues 1377–1830); chicken brain full tail of myosin-Va (FT-GFP – residues 899 – 1830) (Figure S4 for a schematic of the constructs).

### Immunocytochemistry

Cells (6.0×10^4^) were grown in 24 wells plates on coverslips in appropriate culture medium, as described above. After 18-24h they were washed and fixed with 2% paraformaldehyde (w/v) in PBS, pH 7.4, for 15min. They were then permeabilized with 0.3% (v/v) Triton X-100, blocked with 100 mM glycine, and then 3% (w/v) BSA-PBS at room temperature. Cells were then probed with desired primary antibody. Coverslips without primary antibody were used as a negative control and incubated overnight at 4°C in a humidified chamber. Cells were washed extensively with PBS and probed with 1 μg/mL of secondary antibody Alexa Fluor^®^ donkey anti-rabbit 594 (Abcam: ab150073) end/or donkey anti-mouse 488 (W4021 – Promega, Madison, WI, USA) IgG. The coverslips were mounted on ProLong^®^ Diamond Antifade Mountant medium with DAPI (P36962 – Thermo Fisher Scientific, Waltham, MA, EUA). When indicated, cells were treated with 4μM of ionomycin for 5 minutes before fixation and then treated as described above.

Antibodies used: rabbit polyclonal antibody, affinity purified, recognizing myosin Va and rat polyclonal antibody, affinity purified, recognizing myosin Va^60,61^; TOMM20 [EPR15581] – mitochondrial marker (Abcam: ab186734) (Rabbit); Cytochrome C – Mouse monoclonal antibody (BD Pharmingen: #556432); phosphoDRP1(S616) – (Cell Signaling: #3455) Rabbit mAb; DRP1 D6C7 (Cell Signaling: #8570); MFN1 D6E2S (Cell Signaling: #14739) Rabbit mAb; MFN2 D2D10 (Cell Signaling: #9482) Rabbit mAb; Alexa Fluor 594 Phalloidin (ThermoFisher Scientific: # A12381); p44/42 MAPK (Erk1/2) (Cell Signaling: #137f5) Rabbit mAb; Phospho p44/42 MAPK (Erk1/2) (Thr202/Tyr204) (Cell Signaling: #137f5) Mouse mAb.

### Structured Illumination Microscopy (SR-SIM)

The immunofluorescence stained cells were subjected to SR-SIM in an ELYRA S.1 microscope (Carl Zeiss Microimaging), equipped with a 488-nm laser (100 mW), a 561-nm laser (100 mW) and an Andor EM-CCD camera (iXon DU 885). Thin (0·1 mm) Z-stacks of high-resolution image frames were collected in 5 rotations by utilizing an alpha Plan-Apochromat 63×/1·46 oil DIC M27ELYRA objective. Image acquisition, reconstruction, and alignment for structured illumination microscopy were performed using the Zeiss ZEN 2012 SP1 software (black edition, version 8.1.5.484), as previously described^62^.

### Transmission electron microscopy (TEM)

Cells (2.0×10^5^) were grown in a 6-well plate in appropriate cell medium as described above. After 18-24h cells were fixed with 2% glutaraldehyde, 2% formaldehyde, 0,5% CaCl_2_ in PBS with Ca+^2^ and Mg^+2^ for 2 hours at 37°C. Then cells were washed with 0.1M cacodylate buffer (pH 7.4) for 1h at 4°C. The fixed cells were post-fixed with 1% OsO_4_ for 2 hours at 4°C, washed with H_2_O, and dehydrated in a graded series of ethanol (30% to 100%), infiltrated with propylene oxide and embedded in Embed 812 resin, polymerized for 72 hours at 60°C. Thin sections were stained with uranyl acetate and lead citrate for 10min and examined with an electron microscope (Jeol JEM-100 CXII). A unique experiment with two grades were analyzed.

### Metabolite Detection Assays

ROS-Glo™ H_2_O_2_ assay (Promega cat. #G8820) was used to the level of hydrogen peroxide (H_2_O_2_) produced by the cells. For this, 10,000 cells were plated on 96-well plates the day before the experiment. To perform the assay, H_2_O_2_ was added for 4 hours, and the manufacturer’s recommendations were followed.

Glucose Uptake-Glo™ assay (Promega cat. #J1341) was used to measure the glucose uptake in the cell culture. For this, 20,000 cells were plated on 96-well plates the day before the experiment, and the manufacturer’s recommendations followed.

Lactate secretion was measured in cell media of cultured cells using the Lactate-Glo™ assay (Promega cat. # J5021). For this, 50,000 cells were plated on 24-well plates, media was collected 48h after, and the manufacturer’s recommendations were followed.

### Clonogenic assay

300 cells were transferred to 6 well plates and maintained for 10 days in a cell incubator under 95% air and 5% CO2 at 37 °C. After this period, the cells were fixed with 4% paraformaldehyde (w/v) in PBS (pH 7.4) and stained with 2% violet crystal (w/v) in 0.2% ethanol (v/v) for visualization of the colonies. The dishes were photographed and the colonies number were counted.

### Respiratory rates

Oxygen consumption was monitored using a High-Resolution Respirometer (OROBOROS Oxygraph) equipped with DatLab 5.0 (Oroboros, Innsbruk, Austria). Cells were cultured in 100mm dishes, the FBS was removed 24h prior the experiment. In the day of the experiment, the cells were detached and counted 1×10^6^ cells for each condition, that were incubated in 2 mL air saturated MiR05 respiration buffer (20 mM HEPES, 10 mM KH2PO4, 110 mM sucrose, 20 mM taurine, 60 mM K-lactobionate, 0.5 mM EGTA, 3 mM MgCl2, 1 g/L fatty acid-free BSA, pH 7.1) at 37°C, 300 rpm stirring. The cell membrane permeabilization was performed with the addition of 25 μM digitonin in the beginning of the experiment. Respiratory States were determined in the presence of: exogenous respiratory substrates 1.5 mM pyruvate, 0.28 mM malate and 1.3 mM glutamate (Basal State); 240 μM ADP (phosphorylating or State P); 0.5 μg/mL oligomycin (proton leak, non-phosphorylating or State L); 2 additions of 0.25 μM (titration) of carbonyl cyanide 3-chlorophenylhydrazone (CCCP) (maximal respiratory capacity, or State E); and 0.5 μg/mL antimycin A (non-mitochondrial respiration, residual or Rox).

### Myosin-Va affinity chromatography and cloning of candidate genes

Performed by R. Taranath and M. Stern according to the methods of Finan, Hartman and Spudich with the following modifications^39^. Briefly, the melanocyte isoform (Roland et al., 2009) of mouse myosin-Va globular tail domain containing some of the coiled-coil region (to aid in dimerization and stability) was purified and used to construct an affinity column according to previous protocols^39^. The cargo binding domain of myosin-VI was used as a comparison and internal control. Brain extract from 6-8 week-old BL6 mice was flowed onto the column, and proteins were eluted with increasing salt washes. Proteins and protein complexes binding most tightly to the MVa column were eluted with 1M KCl, and analyzed using SDS-PAGE and mass spectrometry.

### Confirmation of binding of Spire1C to myosin-Va

Direct binding assays were performed according to Finan, Hartman, and Spudich^39^. Briefly, a myc-Spire1C construct lacking the N-terminal KIND domain was translated *in vitro* using rabbit reticulocyte lysates. Binding reactions were performed by combining translated myc-Spire1C products with purified myosin-Va or myosin-VI tail domains attached to glutathione-agarose; myosin VI was used as a control for non-specific binding. Proteins co-purifying with myosin-Va or myosin-VI were isolated and analyzed with immunoblots. An HRP-conjugated mouse anti-myc antibody (Abcam) was used at 1:10000 prior to developing the blot with the SuperSignal West Pico Chemiluminescent Substrate kit (Pierce).

### Spire1C immunoprecipitation

Myc-tagged Spire1C full-length and ΔExonC constructs (or empty pcDNA3.1(+) vector) were transfected into a 10-cm dish of HEK293T cells using Lipofectamine 2000 according to manufacturer’s protocol. 48 hours posttransfection, cells were harvested and lysed in RIPA buffer (Thermo #89901) supplemented with Roche complete protease inhibitor cocktail tablet and 1 mM PMSF. Cells were lysed on ice for 20 min followed by centrifugation at 20,000×g for 20 min at 4°C to remove cell debris. Cell lysates were incubated with myc-tag antibody (Cell Signaling 2276S) at 4°C overnight. The next morning, pre-washed protein A/G agarose beads (Santa Cruz Biotechnology, sc-2003) were added and rotated at 4°C for 2 hours. Bound proteins were washed four times with 1×TBST and eluted with 1×SDS loading dye. The eluents were then processed for proteomics.

### Mass Spectrometry and Data Analysis

The digested samples were analyzed on a Q Exactive mass spectrometer (Thermo). The digest was injected directly onto a 30 cm, 75 μm ID column packed with BEH 1.7 μm C18 resin (Waters). Samples were separated at a flow rate of 200 nl/min on a nLC 1000 (Thermo). Buffer A and B were 0.1% formic acid in water and acetonitrile, respectively. A gradient of 5-40% B over 110 min, an increase to 50% B over 10min, an increase to 90% B over another 10min and held at 90% B for a final 10 min of washing was used for 140 min total run time. Column was re-equilibrated with 20 μl of buffer A prior to the injection of sample. Peptides were eluted directly from the tip of the column and nanosprayed directly into the mass spectrometer by application of 2.5 kV voltage at the back of the column. The Q Exactive was operated in a data dependent mode. Full M\S^63^ scans were collected in the Orbitrap at 70K resolution with a mass range of 400 to 1800 m/z and an AGC target of 5e^6^. The ten most abundant ions per scan were selected for MS/MS analysis with HCD fragmentation of 25NCE, an AGC target of 5e^6^ and minimum intensity of 4e^3^. Maximum fill times were set to 60 ms and 120 ms for MS and MS/MS scans respectively. Quadrupole isolation of 2.0 m/z was used, dynamic exclusion was set to 15 sec and unassigned charge states were excluded.

Protein and peptide identification were done with Integrated Proteomics Pipeline – IP2 (Integrated Proteomics Applications). Tandem mass spectra were extracted from raw files using RawConverter^63^ and searched with ProLuCID^64^ against human UniProt database. The search space included all fully-tryptic and half-tryptic peptide candidates with maximum of two missed cleavages. Carbamidomethylation on cysteine was counted as a static modification. Data was searched with 50 ppm precursor ion tolerance and 50 ppm fragment ion tolerance. Data was filtered to 10 ppm precursor ion tolerance post search. Identified proteins were filtered using DTASelect^65^and utilizing a target-decoy database search strategy to control the false discovery rate to 1% at the protein level.

### Acquisition, processing and analysis of confocal images

Confocal images were collected using Leica SP5 or Leica SP8 confocal microscopes, with a oil 63x 1.4NA oil immersion objective. When indicated, the images were imaged with a 63x 1.4NA oil objective on a ZEISS 880 LSM Airyscan confocal system with an inverted stage and heated incubation system with 5% CO2 control, also used for the timelapses. The images were processed and analyzed using Icy-Bioimage and NIH-developed Image J software (Wayne Rasband; National Institutes of Health, Bethesda, MD; available at https://imagej.net). The colocalization was analyzed in the Icy-Bioimage with the Colocalization Studio plugin (Lagache et al., 2015) used to calculate the Pearson coefficients, in addition one of the channels used for colocalization was rotated 180° and resubmitted to the analysis being used as negative control (random colocalization). Electron micrographs were collected in a transmission microscopy Jeol JEM-100 CXII. All the compared images were acquired with the same parameters. For visualization purposes, color, brightness and contrast, when necessary, were modified, always maintaining the equivalent proportions. The raw files were used for all quantifications.

## ACKNOWLEDGMENTS

We are grateful to Prof. Dr. Marcos Roberto Chiaratti for the antibodies against mitochondrial dynamics proteins and to Silmara Banzi Reis and Benedita Oliveira Souza for the technical assistance. We also thank the Multiuser Laboratories on Confocal and Multiphoton Microscopy of the Department of Cell and Molecular Biology, Faculty of Medicine of Ribeirão Preto, University of São Paulo; the Department of Physics and Chemistry, Faculty of Pharmaceutical Sciences of Ribeirão Preto. We are also grateful to the Centro Nacional de Bioimagem (CENABIO) for the use of the super-resolution microscopy facility. This work was supported by São Paulo Research Foundation (FAPESP grants 2014/18189-5 and 2018/04017-9 (to E.M.E.); and fellowships 2016/10862-8 (to J.S.A); 2012/13900-7 (to C.L.S.P.); (2019/00849-2) and (2014/03989-6) to (R.M.P.S.J.);; FAPESP grant to multiuser facilities (2004/08868-0; 2009/54014-7); National Council for Scientific and Technological Development (CNPq grants 457603/2013-5 and 309187/2015-0 (E.M.E). T.Z., M.W., C.S., and U.M. were all supported by the Waitt Foundation and Core Grant application NCI CCSG (CA014195.

## AUTHOR CONTRIBUTIONS

Cell culture, shRNA knockdown, immunocytochemistry, RT-qPCR, metabolite assays, MET imaging (J.S.A); confocal imaging and imaging quantifications (J.S.A, R.M.P.S.J., T.Z., M.W., C.S., U.M.); Respiratory rates (A.O.S., L.C.A.) conception and design (J.S.A., C.L.S.P, U.M., E.M.E.); data analysis and interpretation (all authors); manuscript writing (J.S.A., R.M.P.S.J., U.M., and E.M.E.); figure preparation (J.S.A.); manuscript revision (all authors).

---

**Figure S1.**
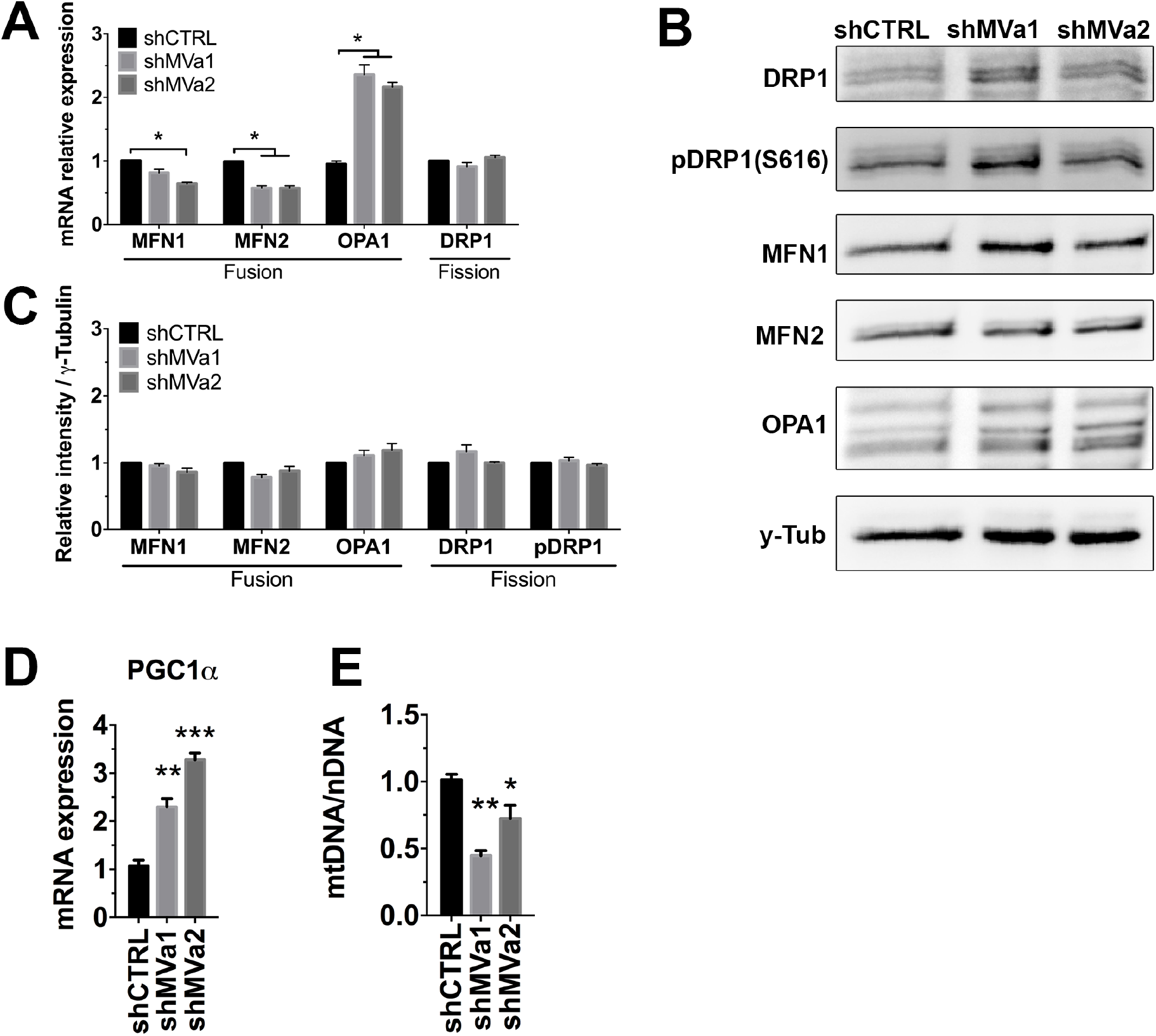
Myosin-Va knockdown changes the expression genes related to biogenesis and mitochondrial dynamics, and alters mtDNA copy number. **(A)** Genes related to mitochondrial dynamics. The GAPDH gene was used as housekeeping. **(B)** Western blots containing total lysate of A375 cells (line 1-shCTRL; line 2-shMVa1 and line 3-shMVa2) probed with antibodies against DRP1, p(S616)DRP1, MFN1, MFN2, OPA1 and γ-Tubulin (γ-Tub). γ-Tubulin was utilized as protein load normalizer. **(C)** Representative plot of the densitometry quantification of the proteins from (B) relative to γ-Tubulin. **(D)** qRT-PCR to determine the level of expression of PGC1α. The GAPDH gene was used as housekeeping. **(E)** qRT-PCR to determine the number of copies of mitochondrial DNA in A375 cells. gDNA was amplified using primers specific for nuclear DNA (nDNA) and primers specific for mitochondrial DNA (mtDNA) (Phillips et al., 2014). The formula 2^-ΔΔCt^ was used for normalizing the mRNA expression level and mtDNA. Data are mean ±SD of three independents experiments. *P<0.01; **P<0.001; ***P<0.0005.

**Figure S2.**
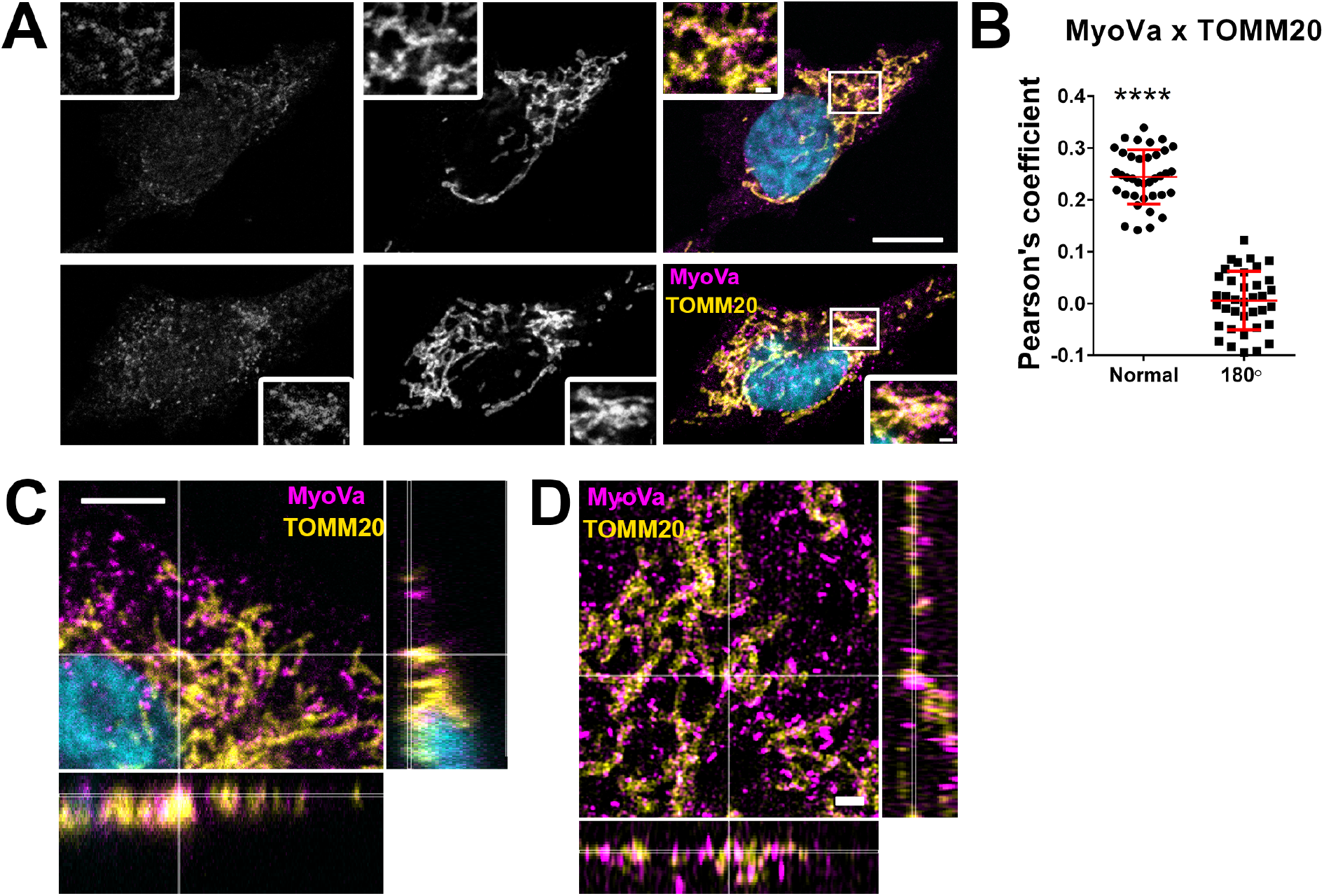
Myosin-Va colocalizes with mitochondrial network in A375 cell line. **(A)** Confocal microscopy images in A375 cells showing the immunolocalization of myosin-Va (magenta) and TOMM20 (yellow), a marker for mitochondrial network. Scale bar: 10μm and 1μm in the insets. **(B)** Pearson’s correlation coefficients support the partial colocalization of Myosin-Va and TOMM20 labeling, (n=37 cells from two independent experiments). The correlation between Myosin-Va (rotated 180°) and TOMM20 channels was used as negative control (random colocalization). **(C)** Orthogonal projections of z-stacks of A375 cells showing the immunolocalization of myosin-Va (magenta) and TOMM20 (yellow), and small clusters colocalized with the mitochondrial network in the OMM. Scale bar: 10 μm or 1 μm in the inserts. Images obtained using Leica SP8 confocal microscope stacks (z=7.14 μm in 34 stacks) and **(D)** images obtained using SR-SIM in an ELYRA S.1 microscope (Carl Zeiss Microimaging) (z=2.64 μm in 24 stacks).

**Figure S3.**
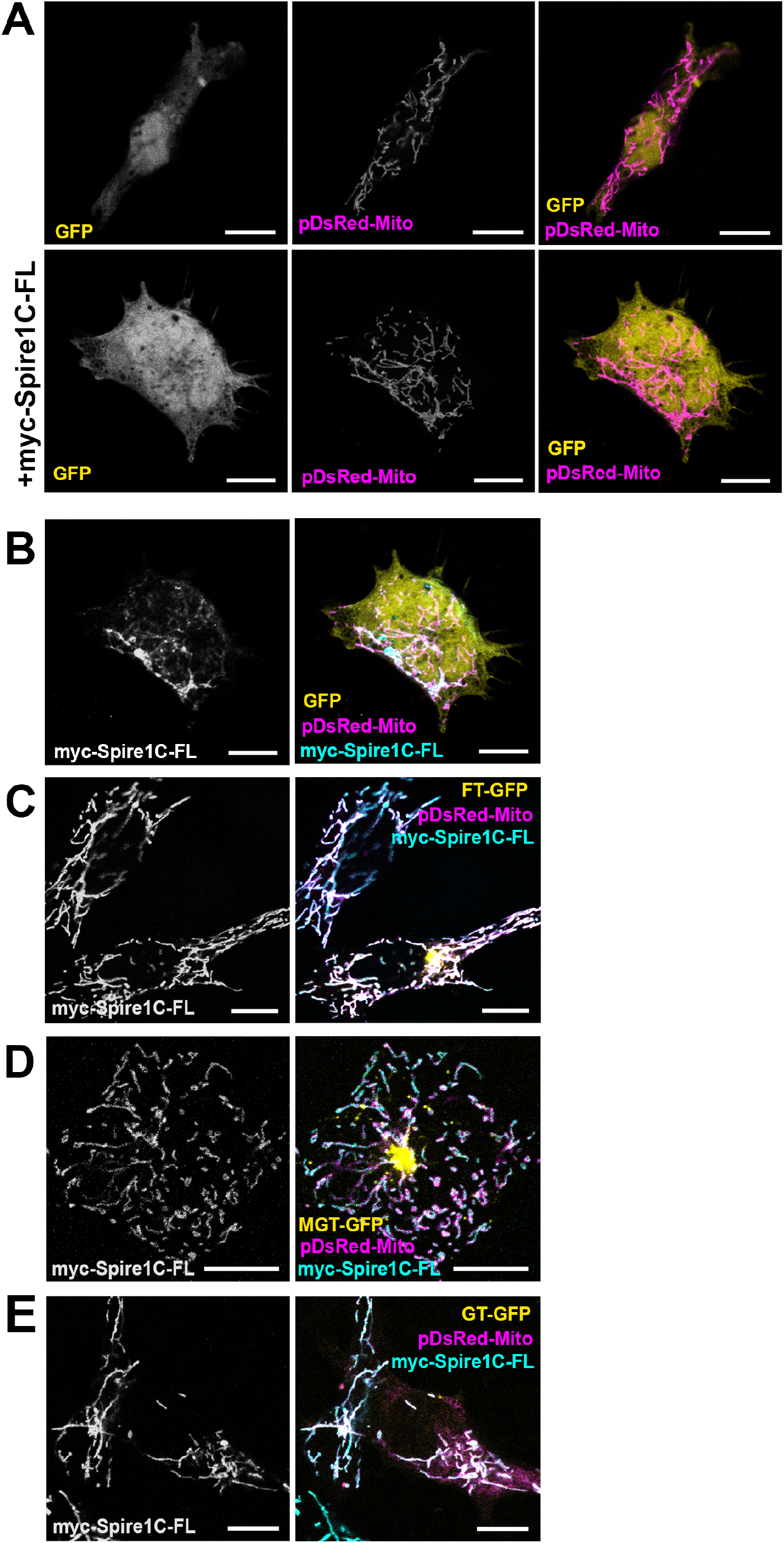
**(A)** pEGFP (GFP) expression along with pDsRed-Mito in A375 cells, as control for tail expressions shown in Figure 6, below the combined expression with myc-Spire1C-FL showing that GFP is not recruited to the mitochondrial network. Representative image of myc-Spire1C-FL overexpression (blue) coexpressed with **(A)** FT-GFP (chicken brain myosin-Va full tail); **(B)** MGT-GFP (chicken brain myosin-Va medial-globular tail); **(C)** GT-GFP (chicken brain myosin-Va globular tail) expression along with pDsRed-Mito in A375 cells showing recruitment of the myosin-Va tails to the mitochondrial network. Scale bar: 10μm and 1μm in the insets

**Figure S4.**
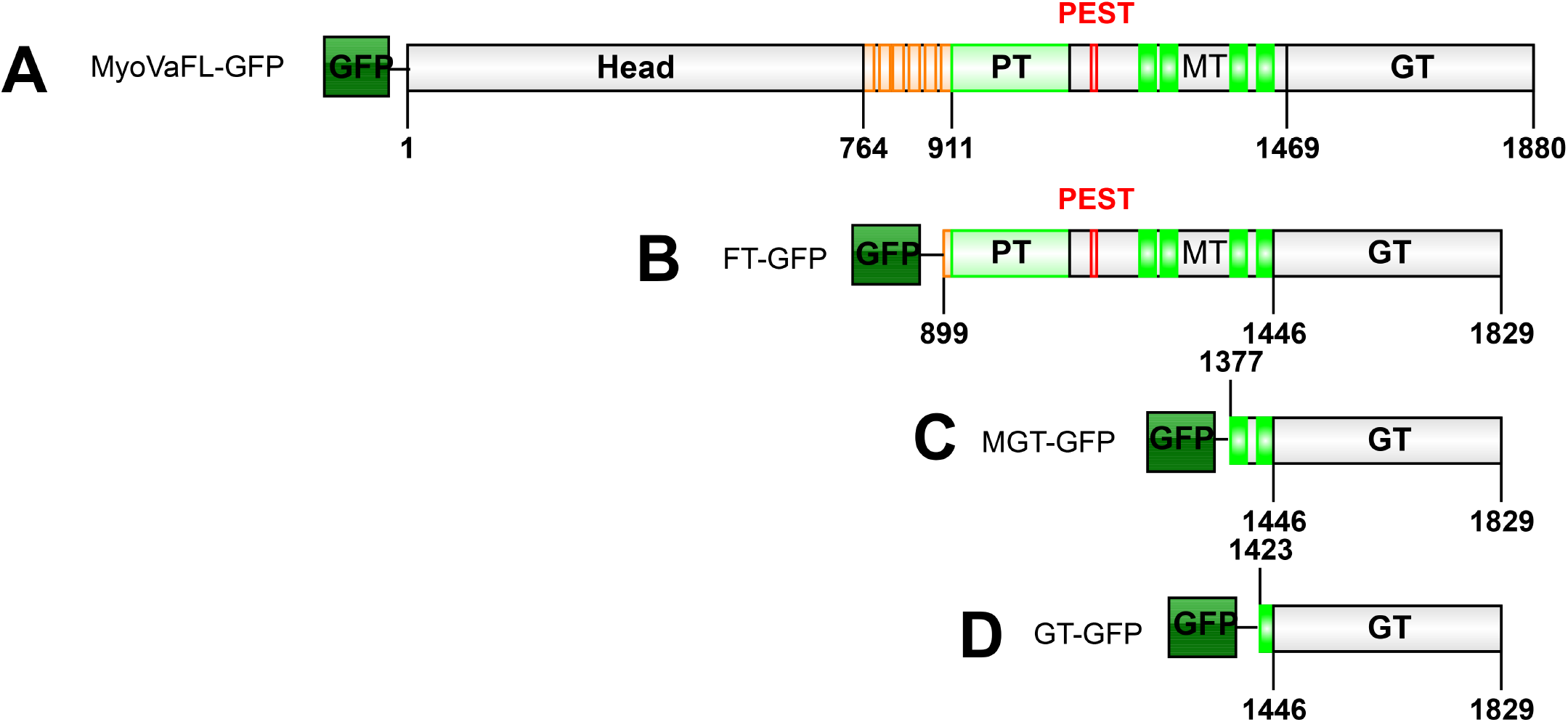
Schematic diagram of myosin-Va constructs. **(A)** Representation of the full length murine myosin-Va fused with GFP tag. **(B, C, D)** Representation of chicken myosin-Va tail constructs fused with GFP tag: (B) full tail, (C) medial globular tail, (D) globular tail. The position of the structural sub-domains are shown in: Head; Neck (IQ-motifs); PT, proximal tail; MT, medial tail; GT, globular tail; MGT, medial domain fragment and globular tail. IBS freely available online was used for the schemes^3^.

**Supplementary Movie 1. Live cell imaging of mitotracker in A375 control cells**. Airscan live imaging of A375 transduced with shCTRL showing the dynamics of the mitochondrial network. Relative to Figure 2C.

**Supplementary Movie 2. Live cell imaging of mitotracker in myosin-Va knockdown cells**. Airscan live imaging of A375 transduced with shMVa2 showing the dynamics of the mitochondrial network. Relative to Figure 2C.

**Supplementary Movie 3. Live cell imaging of myosin-Va at fission sites**. Relative to Figure 7A. Airscan live imaging of U2OS cells showing the localization of MyoVaFL-GFP (green) (full length human myosin-Va) with a mitochondrial marker (magenta) (Mito-RFP) in a control condition.

**Supplementary Movie 4. Live cell imaging of myosin-Va at fission sites upon ionomycin treatment**. Relative to figure 7B. Airscan live imaging of U2OS cells showing the localization of MyoVaFL-GFP (green) (full length human myosin-Va) with a mitochondrial marker (magenta) (Mito-RFP) upon treatment with 4μM of ionomycin.

## REFERENCES

1 Izidoro-Toledo, T. C. et al. A myosin-Va tail fragment sequesters dynein light chains leading to apoptosis in melanoma cells. Cell Death Dis 4, e547, doi:10.1038/cddis.2013.45 (2013).

2 Manor, U. et al. A mitochondria-anchored isoform of the actin-nucleating spire protein regulates mitochondrial division. eLife 4, doi:10.7554/eLife.08828 (2015).

3 Liu, W. et al. IBS: an illustrator for the presentation and visualization of biological sequences. Bioinformatics 31, 3359–3361, doi:10.1093/bioinformatics/btv362 (2015).

4 Serasinghe, M. N. et al. Mitochondrial division is requisite to RAS-induced transformation and targeted by oncogenic MAPK pathway inhibitors. Mol Cell 57, 521–536, doi:10.1016/j.molcel.2015.01.003 (2015).

5 Kashatus, J. A. et al. Erk2 phosphorylation of Drp1 promotes mitochondrial fission and MAPK-driven tumor growth. Mol Cell 57, 537–551, doi:10.1016/j.molcel.2015.01.002 (2015).

6 Otera, H., Ishihara, N. & Mihara, K. New insights into the function and regulation of mitochondrial fission. Biochim Biophys Acta 1833, 1256–1268, doi:10.1016/j.bbamcr.2013.02.002 (2013).

7 Zhao, J. et al. Human MIEF1 recruits Drp1 to mitochondrial outer membranes and promotes mitochondrial fusion rather than fission. Embo j 30, 2762–2778, doi:10.1038/emboj.2011.198 (2011).

8 Loson, O. C., Song, Z., Chen, H. & Chan, D. C. Fis1, Mff, MiD49, and MiD51 mediate Drp1 recruitment in mitochondrial fission. Mol Biol Cell 24, 659–667, doi:10.1091/mbc.E12-10-0721 (2013).

9 MacVicar, T. & Langer, T. OPA1 processing in cell death and disease – the long and short of it. J Cell Sci 129, 2297–2306, doi:10.1242/jcs.159186 (2016).

10 Espreafico, E. M. et al. Localization of myosin-V in the centrosome. Proc Natl Acad Sci U S A 95, 8636–8641 (1998).

11 Assis, L. H. et al. The molecular motor Myosin Va interacts with the cilia-centrosomal protein RPGRIP1L. Sci Rep 7, 43692, doi:10.1038/srep43692 (2017).

12 Pranchevicius, M. C. et al. Myosin Va phosphorylated on Ser1650 is found in nuclear speckles and redistributes to nucleoli upon inhibition of transcription. Cell Motil Cytoskeleton 65, 441–456, doi:10.1002/cm.20269 (2008).

13 Nascimento, A. A., Amaral, R. G., Bizario, J. C., Larson, R. E. & Espreafico, E. M. Subcellular localization of myosin-V in the B16 melanoma cells, a wild-type cell line for the dilute gene. Mol Biol Cell 8, 1971–1988 (1997).

14 Takagishi, Y. et al. The dilute-lethal (dl) gene attacks a Ca2+ store in the dendritic spine of Purkinje cells in mice. Neurosci Lett 215, 169–172 (1996).

15 Alves, C. P. et al. Myosin-Va contributes to manifestation of malignant-related properties in melanoma cells. J Invest Dermatol 133, 2809–2812, doi:10.1038/jid.2013.218 (2013).

16 Puthalakath, H. et al. Bmf: a proapoptotic BH3-only protein regulated by interaction with the myosin V actin motor complex, activated by anoikis. Science 293, 1829–1832, doi:10.1126/science.1062257 (2001).

17 Boldogh, I. R., Ramcharan, S. L., Yang, H. C. & Pon, L. A. A type V myosin (Myo2p) and a Rab-like G-protein (Ypt11p) are required for retention of newly inherited mitochondria in yeast cells during cell division. Mol Biol Cell 15, 3994–4002, doi:10.1091/mbc.E04-01-0053 (2004).

18 Pathak, D., Sepp, K. J. & Hollenbeck, P. J. Evidence that myosin activity opposes microtubule-based axonal transport of mitochondria. J Neurosci 30, 8984–8992, doi:10.1523/JNEUROSCI.1621-10.2010 (2010).

19 Sun, Y., Chiu, T. T., Foley, K. P., Bilan, P. J. & Klip, A. Myosin Va mediates Rab8A-regulated GLUT4 vesicle exocytosis in insulin-stimulated muscle cells. Mol Biol Cell 25, 1159–1170, doi:10.1091/mbc.E13-08-0493 (2014).

20 Chen, Y. et al. Rab10 and myosin-Va mediate insulin-stimulated GLUT4 storage vesicle translocation in adipocytes. J Cell Biol 198, 545–560, doi:10.1083/jcb.201111091 (2012).

21 Yoshizaki, T. et al. Myosin 5a is an insulin-stimulated Akt2 (protein kinase Bbeta) substrate modulating GLUT4 vesicle translocation. Mol Cell Biol 27, 5172–5183, doi:10.1128/MCB.02298-06 (2007).

22 Li, S. et al. Transient assembly of F-actin on the outer mitochondrial membrane contributes to mitochondrial fission. J Cell Biol 208, 109–123, doi:10.1083/jcb.201404050 (2015).

23 Hatch, A. L., Ji, W. K., Merrill, R. A., Strack, S. & Higgs, H. N. Actin filaments as dynamic reservoirs for Drp1 recruitment. Mol Biol Cell 27, 3109–3121, doi:10.1091/mbc.E16-03-0193 (2016).

24 Ji, W., Hatch, A. L., Merrill, R. A., Strack, S. & Higgs, H. N. in eLife Vol. 4 (2015).

25 Korobova, F., Ramabhadran, V. & Higgs, H. N. An actin-dependent step in mitochondrial fission mediated by the ER-associated formin INF2. Science 339, 464–467, doi:10.1126/science.1228360 (2013).

26 Korobova, F., Gauvin, T. J. & Higgs, H. N. A role for myosin II in mammalian mitochondrial fission. Curr Biol 24, 409–414, doi:10.1016/j.cub.2013.12.032 (2014).

27 De Vos, K. J., Allan, V. J., Grierson, A. J. & Sheetz, M. P. Mitochondrial function and actin regulate dynamin-related protein 1-dependent mitochondrial fission. Curr Biol 15, 678–683, doi:10.1016/j.cub.2005.02.064 (2005).

28 Schiavon, C. et al. Actin chromobody imaging reveals sub-organellar actin dynamics. doi:10.1101/639278 (2019).

29 Rehklau, K. et al. Cofilin1-dependent actin dynamics control DRP1-mediated mitochondrial fission. Cell death & disease 8, e3063, doi:10.1038/cddis.2017.448 (2017).

30 Yang, C. & Svitkina, T. M. Ultrastructure and dynamics of the actin-myosin II cytoskeleton during mitochondrial fission. Nat Cell Biol 21, 603–613, doi:10.1038/s41556-019-0313-6 (2019).

31 Moore, A. S. & Holzbaur, E. L. Dynamic recruitment and activation of ALS-associated TBK1 with its target optineurin are required for efficient mitophagy. Proc Natl Acad Sci U S A 113, E3349–3358, doi:10.1073/pnas.1523810113 (2016).

32 Pylypenko, O. et al. Coordinated recruitment of Spir actin nucleators and myosin V motors to Rab11 vesicle membranes. Elife 5, doi:10.7554/eLife.17523 (2016).

33 Montaville, P. et al. Spire and Formin 2 synergize and antagonize in regulating actin assembly in meiosis by a ping-pong mechanism. PLoS biology 12, e1001795, doi:10.1371/journal.pbio.1001795 (2014).

34 Pfender, S., Kuznetsov, V., Pleiser, S., Kerkhoff, E. & Schuh, M. Spire-type actin nucleators cooperate with Formin-2 to drive asymmetric oocyte division. Curr Biol 21, 955–960, doi:10.1016/j.cub.2011.04.029 (2011).

35 Schuh, M. An actin-dependent mechanism for long-range vesicle transport. Nat Cell Biol 13, 1431–1436, doi:10.1038/ncb2353 (2011).

36 Makrecka-Kuka, M., Krumschnabel, G. & Gnaiger, E. High-Resolution Respirometry for Simultaneous Measurement of Oxygen and Hydrogen Peroxide Fluxes in Permeabilized Cells, Tissue Homogenate and Isolated Mitochondria. Biomolecules 5, 1319–1338, doi:10.3390/biom5031319 (2015).

37 McDonald, J. H. & Dunn, K. W. Statistical tests for measures of colocalization in biological microscopy. J Microsc 252, 295–302, doi:10.1111/jmi.12093 (2013).

38 Nascimento, A. A., Cheney, R. E., Tauhata, S. B., Larson, R. E. & Mooseker, M. S. Enzymatic characterization and functional domain mapping of brain myosin-V. J Biol Chem 271, 17561–17569 (1996).

39 Finan, D., Hartman, M. A. & Spudich, J. A. Proteomics approach to study the functions of Drosophila myosin VI through identification of multiple cargo-binding proteins. Proc Natl Acad Sci USA 108, 5566–5571, doi:10.1073/pnas.1101415108 (2011).

40 Abildgaard, C. & Guldberg, P. Molecular drivers of cellular metabolic reprogramming in melanoma. Trends Mol Med 21, 164–171, doi:10.1016/j.molmed.2014.12.007 (2015).

41 Warburg, O. On the origin of cancer cells. Science 123, 309–314 (1956).

42 Chen, H. & Chan, D. C. Mitochondrial Dynamics in Regulating the Unique Phenotypes of Cancer and Stem Cells. Cell Metab 26, 39–48, doi:10.1016/j.cmet.2017.05.016 (2017).

43 Guido, C. et al. Mitochondrial fission induces glycolytic reprogramming in cancer-associated myofibroblasts, driving stromal lactate production, and early tumor growth. Oncotarget 3, 798–810, doi:10.18632/oncotarget.574 (2012).

44 Bordi, M., Nazio, F. & Campello, S. The Close Interconnection between Mitochondrial Dynamics and Mitophagy in Cancer. Front Oncol 7, 81, doi:10.3389/fonc.2017.00081 (2017).

45 Liberti, M. V. & Locasale, J. W. The Warburg Effect: How Does it Benefit Cancer Cells? Trends Biochem Sci 41, 211–218, doi:10.1016/j.tibs.2015.12.001 (2016).

46 Hoque, A. et al. Mitochondrial fission protein Drp1 inhibition promotes cardiac mesodermal differentiation of human pluripotent stem cells. Cell Death Discov 4, 39, doi:10.1038/s41420-018-0042-9 (2018).

47 Kitamura, S. et al. Drp1 regulates mitochondrial morphology and cell proliferation in cutaneous squamous cell carcinoma. J Dermatol Sci 88, 298–307, doi:10.1016/j.jdermsci.2017.08.004 (2017).

48 Hammer, J. A., 3rd & Wagner, W. Functions of class V myosins in neurons. J Biol Chem 288, 28428–28434, doi:10.1074/jbc.R113.514497 (2013).

49 Lu, H., Krementsova, E. B. & Trybus, K. M. Regulation of myosin V processivity by calcium at the single molecule level. J Biol Chem 281, 31987–31994, doi:10.1074/jbc.M605181200 (2006).

50 Wada, F. et al. Myosin Va and Endoplasmic Reticulum Calcium Channel Complex Regulates Membrane Export during Axon Guidance. Cell Rep 15, 1329–1344, doi:10.1016/j.celrep.2016.04.021 (2016).

51 Maschi, D., Gramlich, M. W. & Klyachko, V. A. Myosin V functions as a vesicle tether at the plasma membrane to control neurotransmitter release in central synapses. Elife 7, doi:10.7554/eLife.39440 (2018).

52 Wales, P. et al. Calcium-mediated actin reset (CaAR) mediates acute cell adaptations. eLife 5, doi:10.7554/eLife.19850 (2016).

53 Chakrabarti, R. et al. INF2-mediated actin polymerization at the ER stimulates mitochondrial calcium uptake, inner membrane constriction, and division. J Cell Biol 217, 251–268, doi:10.1083/jcb.201709111 (2018).

54 Friedman, J. R. et al. ER tubules mark sites of mitochondrial division. Science 334, 358–362, doi:10.1126/science.1207385 (2011).

55 Wong, Y. C., Ysselstein, D. & Krainc, D. Mitochondria-lysosome contacts regulate mitochondrial fission via RAB7 GTP hydrolysis. Nature 554, 382–386, doi:10.1038/nature25486 (2018).

56 Landry, M. C. et al. A functional interplay between the small GTPase Rab11a and mitochondria-shaping proteins regulates mitochondrial positioning and polarization of the actin cytoskeleton downstream of Src family kinases. J Biol Chem 289, 2230–2249, doi:10.1074/jbc.M113.516351 (2014).

57 Wagner, W., Brenowitz, S. D. & Hammer, J. A., 3rd. Myosin-Va transports the endoplasmic reticulum into the dendritic spines of Purkinje neurons. Nat Cell Biol 13, 40–48, doi:10.1038/ncb2132 (2011).

58 Phillips, N. R., Sprouse, M. L. & Roby, R. K. Simultaneous quantification of mitochondrial DNA copy number and deletion ratio: a multiplex real-time PCR assay. Sci Rep 4, 3887, doi:10.1038/srep03887 (2014).

59 Wu, X. S. et al. Identification of an organelle receptor for myosin-Va. Nat Cell Biol 4, 271–278, doi:10.1038/ncb760 (2002).

60 Espindola, F. S. et al. Biochemical and immunological characterization of p190-calmodulin complex from vertebrate brain: a novel calmodulin-binding myosin. J Cell Biol 118, 359–368, doi:10.1083/jcb.118.2.359 (1992).

61 Espreafico, E. M. et al. Primary structure and cellular localization of chicken brain myosin-V (p190), an unconventional myosin with calmodulin light chains. J Cell Biol 119, 1541–1557 (1992).

62 Rocha, M. R. et al. Annexin A2 overexpression associates with colorectal cancer invasiveness and TGF-ss induced epithelial mesenchymal transition via Src/ANXA2/STAT3. Sci Rep 8, 11285, doi:10.1038/s41598-018-29703-0 (2018).

63 He, L., Diedrich, J., Chu, Y. Y. & Yates, J. R., 3rd. Extracting Accurate Precursor Information for Tandem Mass Spectra by RawConverter. Anal Chem 87, 11361–11367, doi:10.1021/acs.analchem.5b02721 (2015).

64 Xu, T. et al. ProLuCID: An improved SEQUEST-like algorithm with enhanced sensitivity and specificity. J Proteomics 129, 16–24, doi:10.1016/j.jprot.2015.07.001 (2015).

65 Tabb, D. L., McDonald, W. H. & Yates, J. R., 3rd. DTASelect and Contrast: tools for assembling and comparing protein identifications from shotgun proteomics. J Proteome Res 1, 21–26 (2002).

