## Supplementary material for "A novel role for Myosin-Va in mitochondrial fission": Suppl. Table 1

| Protein Name | Accession Number | Molecular Weight | MyoVI-GTD Spectral Counts | MyoVI-GTD Unique Peptides | MyoVa-MGT Spectral Counts | MyoVa-MGT Unique Peptides |
|---|---|---|---|---|---|---|
| Dync1h1 Cytoplasmic dynein 1 heavy chain 1 | IPI00119876 | 532 kDa | 310 | 121 | 515 | 182 |
| Spna2 Spectrin alpha 2 | IPI00757353 | 285 kDa | 853 | 170 | 597 | 149 |
| Myo5a 215 kDa protein | IPI00875222 | 215 kDa | 162 | 47 | 874 | 109 |
| AU042671 hypothetical protein LOC269700 isoform 1 | IPI00762814 | 453 kDa | 2 | 2 | 231 | 104 |
| Spnb2 Isoform 1 of Spectrin beta chain, brain 1 | IPI00319830 | 274 kDa | 505 | 122 | 347 | 100 |
| Dmxl2 Isoform 1 of DmX-like protein 2 | IPI00853932 | 338 kDa | 63 | 38 | 251 | 100 |
| Cltc Clathrin heavy chain 1 | IPI00169916 (+1) | 192 kDa | 1994 | 138 | 565 | 90 |
| Mtap2 12 days embryo spinal cord cDNA, RIKEN full-length enriched library, clone:C530026I | IPI00894724 | 199 kDa | 229 | 82 | 258 | 74 |
| Mtap1a Isoform 1 of Microtubule-associated protein 1A | IPI00408909 (+1) | 300 kDa | 310 | 86 | 214 | 74 |
| Itpr1 Isoform 4 of Inositol 1,4,5-trisphosphate receptor type 1 | IPI00230019 (+3) | 311 kDa | 37 | 18 | 155 | 73 |
| Huwe1 HECT, UBA and WWE domain containing 1 | IPI00463909 (+1) | 483 kDa | 5 | 5 | 91 | 69 |
| Fasn Fatty acid synthase | IPI00113223 | 272 kDa | 24 | 17 | 140 | 68 |
| Usp9x Ubiquitin carboxyl-terminal hydrolase | IPI00798468 | 291 kDa | 68 | 45 | 98 | 65 |
| Lrp1 Prolow-density lipoprotein receptor-related protein 1 precursor | IPI00119063 | 505 kDa | 92 | 53 | 109 | 62 |
| Myh10 Myosin-10 | IPI00515398 (+1) | 229 kDa | 65 | 40 | 98 | 59 |
| Mical1 NEDD9-interacting protein with calponin homology and LIM domains | IPI00116371 | 117 kDa | 2 | 2 | 203 | 57 |
| Plec1 Isoform PLEC-1I of Plectin-1 | IPI00229509 (+10) | 517 kDa | 121 | 62 | 110 | 57 |
| Nbea Isoform 4 of Neurobeachin | IPI00320836 (+1) | 326 kDa | 7 | 7 | 99 | 55 |
| Dnm1 Isoform 3 of Dynamin-1 | IPI00465648 (+1) | 96 kDa | 57 | 35 | 94 | 54 |
| Ap2b1 Isoform 1 of AP-2 complex subunit beta-1 | IPI00119689 (+1) | 105 kDa | 66 | 33 | 164 | 53 |
| Mtap1b microtubule-associated protein 1B | IPI00896700 | 270 kDa | 314 | 78 | 159 | 53 |
| Bcan brevican isoform 1 | IPI00869394 | 96 kDa | 40 | 11 | 378 | 50 |
| Tnr Isoform 1 of Tenascin-R precursor | IPI00227126 | 150 kDa | 89 | 27 | 327 | 48 |
| Spnb3 Adult male brain UNDEFINED_CELL_LINE cDNA, RIKEN full-length enriched library, clc | IPI00134344 | 271 kDa | 239 | 62 | 152 | 47 |
| Spire1 Isoform 2 of Protein spire homolog 1 | IPI00420800 | 69 kDa | 7 | 7 | 261 | 45 |
| Ptprz1 protein tyrosine phosphatase, receptor type Z, polypeptide 1 | IPI00627008 | 254 kDa | 48 | 20 | 271 | 43 |
| Ubr4 Isoform 1 of E3 ubiquitin-protein ligase UBR4 | IPI00378681 (+3) | 572 kDa | 14 | 14 | 51 | 39 |
| Ncan Neurocan core protein precursor | IPI00135563 | 137 kDa | 34 | 15 | 243 | 38 |
| Hsp90aa1 Heat shock protein HSP 90-alpha | IPI00330804 | 85 kDa | 44 | 28 | 62 | 38 |
| sp_K2C1_HUMAN | IPIsp_K2C1_HUMAN | 66 kDa | 166 | 33 | 110 | 36 |
| Ddb1 DNA damage-binding protein 1 | IPI00316740 | 127 kDa | 101 | 34 | 100 | 36 |
| Hspa8 Heat shock cognate 71 kDa protein | IPI00323357 | 71 kDa | 159 | 41 | 78 | 35 |
| Ank2 Adult retina cDNA, RIKEN full-length enriched library, clone:A930028N13 product:simi | IPI00227235 | 131 kDa | 96 | 29 | 116 | 34 |
| Ap2a1 Isoform A of AP-2 complex subunit alpha-1 | IPI00108780 (+1) | 108 kDa | 59 | 32 | 75 | 34 |
| Nsf Vesicle-fusing ATPase | IPI00656325 (+1) | 83 kDa | 39 | 30 | 48 | 33 |
| sp_K1C10_HUMAN | IPIsp_K1C10_HUMAN | 60 kDa | 105 | 27 | 76 | 32 |
| Dhrs4 NADPH-dependent retinol dehydrogenase/reductase isoform 1 | IPI00318750 | 30 kDa | 10 | 8 | 182 | 31 |

| | | | | | | |
|---|---|---|---|---|---|---|
| Vcp Transitional endoplasmic reticulum ATPase | IPI00622235 | 89 kDa | 23 | 19 | 44 | 31 |
| Hsph1 Isoform HSP105-alpha of Heat shock protein 105 kDa | IPI00123802 (+2) | 96 kDa | 20 | 17 | 37 | 31 |
| Zzef1 Isoform 2 of Zinc finger ZZ-type and EF-hand domain-containing protein 1 | IPI00340123 (+1) | 324 kDa | 0 | 0 | 35 | 31 |
| Tubb2a Tubulin beta-2A chain | IPI00338039 | 50 kDa | 773 | 32 | 419 | 30 |
| Atp1a3 Sodium/potassium-transporting ATPase subunit alpha-3 | IPI00122048 (+1) | 112 kDa | 67 | 24 | 103 | 30 |
| Ubash3b Isoform 1 of Ubiquitin associated and SH3 domain-containing protein B | IPI00331539 | 71 kDa | 5 | 4 | 68 | 30 |
| Atp6v1a Isoform 1 of Vacuolar ATP synthase catalytic subunit A | IPI00407692 | 68 kDa | 29 | 20 | 50 | 30 |
| Tuba1b Tubulin alpha-1B chain | IPI00117348 | 50 kDa | 320 | 31 | 271 | 29 |
| Cnp Isoform CNPI of 2',3'-cyclic-nucleotide 3'-phosphodiesterase | IPI00229598 (+1) | 45 kDa | 116 | 24 | 108 | 29 |
| Hk1 Hk1 protein | IPI00762858 | 102 kDa | 28 | 21 | 40 | 29 |
| Nf1 Isoform 2 of Neurofibromin | IPI00108783 (+1) | 320 kDa | 0 | 0 | 36 | 29 |
| sp_K22E_HUMAN | IPIsp_K22E_HUMAN | 66 kDa | 96 | 24 | 74 | 28 |
| Dpysl2 Dihydropyrimidinase-related protein 2 | IPI00114375 | 62 kDa | 45 | 26 | 73 | 28 |
| Pabpc1 Polyadenylate-binding protein 1 | IPI00124287 (+1) | 71 kDa | 6 | 6 | 56 | 28 |
| Hspa4l Isoform 1 of Heat shock 70 kDa protein 4L | IPI00317710 | 94 kDa | 21 | 17 | 41 | 28 |
| Hspa9 heat shock protein 9 | IPI00880839 | 73 kDa | 68 | 34 | 40 | 28 |
| Syn1 Isoform Ia of Synapsin-1 | IPI00649886 | 74 kDa | 67 | 26 | 155 | 27 |
| Ncam1 Isoform N-CAM 180 of Neural cell adhesion molecule 1, 180 kDa isoform precursor | IPI00122971 | 119 kDa | 108 | 23 | 127 | 27 |
| Plxnb1 Plexin-B1 precursor | IPI00229992 | 231 kDa | 3 | 3 | 70 | 27 |
| Hadha Trifunctional enzyme subunit alpha, mitochondrial precursor | IPI00223092 | 83 kDa | 7 | 6 | 48 | 27 |
| Nfasc Neurofascin precursor | IPI00329927 | 138 kDa | 28 | 18 | 40 | 27 |
| Actg1 Actin, cytoplasmic 2 | IPI00874482 | 42 kDa | 59 | 20 | 122 | 26 |
| Cyfip2 Cytoplasmic FMR1-interacting protein 2 | IPI00405625 | 146 kDa | 28 | 19 | 42 | 26 |
| Hspa4 Heat shock 70 kDa protein 4 | IPI00331556 | 94 kDa | 34 | 25 | 38 | 26 |
| Pi4ka Phosphatidylinositol 4-kinase, catalytic, alpha polypeptide | IPI00115875 | 231 kDa | 5 | 3 | 37 | 26 |
| Htt huntingtin | IPI00271166 | 345 kDa | 0 | 0 | 34 | 26 |
| Wdr7 WD repeat domain 7 | IPI00380997 (+1) | 163 kDa | 25 | 19 | 51 | 25 |
| Vps13c Chorein | IPI00473340 (+3) | 415 kDa | 0 | 0 | 40 | 25 |
| Copa Coatomer subunit alpha | IPI00229834 | 138 kDa | 14 | 12 | 29 | 25 |
| Vcan versican | IPI00875672 | 367 kDa | 20 | 7 | 140 | 24 |
| Hapln1 Hyaluronan and proteoglycan link protein 1 precursor | IPI00131995 | 40 kDa | 4 | 4 | 73 | 24 |
| Nrxn1 Isoform 2 of Neurexin-1-alpha precursor | IPI00230050 (+2) | 165 kDa | 0 | 0 | 63 | 24 |
| Ap2a2 adaptor protein complex AP-2, alpha 2 subunit | IPI00753468 | 104 kDa | 43 | 20 | 61 | 24 |
| Epb4.1l3 Isoform 1 of Band 4.1-like protein 3 | IPI00125501 (+2) | 103 kDa | 55 | 28 | 60 | 24 |
| sp_K1C9_HUMAN | IPIsp_K1C9_HUMAN | 62 kDa | 86 | 23 | 43 | 24 |
| Stxbp1 Isoform 1 of Syntaxin-binding protein 1 | IPI00415402 | 68 kDa | 36 | 25 | 39 | 24 |
| Sept11 Isoform 3 of Septin-11 | IPI00420385 (+2) | 49 kDa | 27 | 18 | 32 | 24 |
| Nefm Neurofilament medium polypeptide | IPI00323800 | 96 kDa | 134 | 44 | 46 | 23 |
| Cntn1 Contactin-1 precursor | IPI00123058 | 113 kDa | 5 | 5 | 40 | 23 |
| Hspa5 78 kDa glucose-regulated protein precursor | IPI00319992 | 72 kDa | 59 | 29 | 39 | 23 |
| Tln2 talin 2 | IPI00229647 | 272 kDa | 5 | 5 | 32 | 23 |

| | | | | | | |
|---|---|---|---|---|---|---|
| Trim32 Tripartite motif-containing protein 32 | IPI00321005 | 72 kDa | 8 | 7 | 29 | 23 |
| Mtap6 microtubule-associated protein 6 isoform 1 | IPI00115833 | 96 kDa | 63 | 25 | 64 | 22 |
| Nefh Neurofilament heavy polypeptide | IPI00114241 (+1) | 117 kDa | 103 | 29 | 49 | 22 |
| Rtn4 Isoform 1 of Reticulon-4 | IPI00469392 | 127 kDa | 56 | 24 | 46 | 22 |
| Synj1 similar to mKIAA0910 protein | IPI00850983 (+3) | 189 kDa | 18 | 12 | 44 | 22 |
| L1cam L1 cell adhesion molecule | IPI00785371 | 141 kDa | 16 | 12 | 43 | 22 |
| Nefl Neurofilament light polypeptide | IPI00554928 | 62 kDa | 69 | 26 | 37 | 22 |
| 5031439G07Rik Novel protein | IPI00464302 | 50 kDa | 0 | 0 | 34 | 22 |
| Ap1b1 Adult male diencephalon cDNA, RIKEN full-length enriched library, clone:9330159F11 | IPI00648683 | 105 kDa | 42 | 4 | 138 | 21 |
| Syn2 Isoform IIa of Synapsin-2 | IPI00469548 | 63 kDa | 51 | 13 | 101 | 21 |
| Ap2m1 AP-2 complex subunit mu-1 | IPI00116356 | 50 kDa | 32 | 13 | 72 | 21 |
| Pdha1 Pyruvate dehydrogenase E1 component subunit alpha, somatic form, mitochondrial p | IPI00337893 | 43 kDa | 15 | 9 | 41 | 21 |
| Idh3b Tumor-related protein | IPI00126635 | 42 kDa | 51 | 23 | 40 | 21 |
| Sept7 cell division cycle 10 homolog | IPI00224626 | 51 kDa | 26 | 14 | 37 | 21 |
| Sbf1 Putative uncharacterized protein | IPI00752390 (+1) | 211 kDa | 2 | 2 | 26 | 21 |
| Lrpprc Leucine-rich PPR motif-containing protein, mitochondrial precursor | IPI00420706 | 157 kDa | 0 | 0 | 23 | 21 |
| Snap91 Isoform Long of Clathrin coat assembly protein AP180 | IPI00122409 (+2) | 92 kDa | 46 | 19 | 48 | 20 |
| Atp6v1b2 Vacuolar ATP synthase subunit B, brain isoform | IPI00119113 | 57 kDa | 11 | 7 | 29 | 20 |
| Atp5b ATP synthase subunit beta, mitochondrial precursor | IPI00468481 | 56 kDa | 19 | 14 | 27 | 20 |
| Cand1 TBP-interacting protein isoform 1 | IPI00420562 (+1) | 159 kDa | 16 | 16 | 20 | 20 |
| Mbp Isoform 4 of Myelin basic protein | IPI00223377 | 22 kDa | 297 | 17 | 228 | 19 |
| Gcdh glutaryl-Coenzyme A dehydrogenase | IPI00788331 | 49 kDa | 0 | 0 | 44 | 19 |
| Ppp2r1a Serine/threonine-protein phosphatase 2A 65 kDa regulatory subunit A alpha isofor | IPI00310091 | 65 kDa | 31 | 19 | 32 | 19 |
| Dctn1 Dctn1 protein | IPI00856365 | 140 kDa | 43 | 31 | 31 | 19 |
| Tcp1 Isoform 1 of T-complex protein 1 subunit alpha B | IPI00459493 | 60 kDa | 16 | 10 | 27 | 19 |
| Nrcam 123 kDa protein | IPI00876075 | 123 kDa | 11 | 10 | 25 | 19 |
| Atp5a1 ATP synthase subunit alpha, mitochondrial precursor | IPI00130280 | 60 kDa | 16 | 13 | 24 | 19 |
| ENSMUSG00000070490;Gapdh;100041325;ENSMUSG00000072451;LOC100048291;100040 | IPI00273646 (+1) | 36 kDa | 80 | 15 | 88 | 18 |
| Hsp90ab1 Heat shock protein 84b | IPI00229080 | 83 kDa | 43 | 12 | 59 | 18 |
| Icam5 intercellular adhesion molecule 5, telencephalin | IPI00877299 | 97 kDa | 8 | 7 | 36 | 18 |
| Hnrnpm Isoform 1 of Heterogeneous nuclear ribonucleoprotein M | IPI00132443 (+1) | 78 kDa | 15 | 14 | 28 | 18 |
| Aldoa Fructose-bisphosphate aldolase A | IPI00221402 | 39 kDa | 7 | 7 | 27 | 18 |
| Cct8 T-complex protein 1 subunit theta | IPI00469268 | 60 kDa | 10 | 9 | 23 | 18 |
| Gpd2 Glycerol phosphate dehydrogenase 2, mitochondrial | IPI00331182 (+1) | 83 kDa | 0 | 0 | 22 | 18 |
| Pkm2 Isoform M1 of Pyruvate kinase isozymes M1/M2 | IPI00845840 | 58 kDa | 7 | 5 | 21 | 18 |
| Cadps Isoform 1 of Calcium-dependent secretion activator 1 | IPI00668903 (+1) | 153 kDa | 6 | 5 | 21 | 18 |
| Kif21a Isoform 1 of Kinesin-like protein KIF21A | IPI00454081 (+4) | 187 kDa | 0 | 0 | 21 | 18 |
| Ryr2 ryanodine receptor 2, cardiac | IPI00338309 | 565 kDa | 0 | 0 | 21 | 18 |
| Ctnna2 Visual cortex cDNA, RIKEN full-length enriched library, clone:K530006G14 product:ca | IPI00230751 | 102 kDa | 26 | 23 | 18 | 18 |
| Itih3 69 kDa protein | IPI00876246 | 69 kDa | 0 | 0 | 68 | 17 |
| Ank2 ankyrin 2, brain isoform 2 | IPI00228697 | 117 kDa | 37 | 12 | 50 | 17 |

| | | | | | | |
|---|---|---|---|---|---|---|
| Camk2a Isoform Alpha CaMKII of Calcium/calmodulin-dependent protein kinase type II alph | IPI00621806 | 54 kDa | 66 | 19 | 48 | 17 |
| Ap1m1 AP-1 complex subunit mu-1 | IPI00119680 | 49 kDa | 0 | 0 | 35 | 17 |
| Ywhae 14-3-3 protein epsilon | IPI00118384 | 29 kDa | 46 | 18 | 31 | 17 |
| LOC100044138 similar to CDCrel-1AI isoform 1 | IPI00850539 (+1) | 42 kDa | 21 | 12 | 31 | 17 |
| Cct3 T-complex protein 1 subunit gamma | IPI00116283 | 61 kDa | 19 | 15 | 30 | 17 |
| Slc25a4 ADP/ATP translocase 1 | IPI00115564 | 33 kDa | 33 | 12 | 29 | 17 |
| Hapln4 Hyaluronan and proteoglycan link protein 4 precursor | IPI00229184 | 43 kDa | 0 | 0 | 26 | 17 |
| Glud1 Glutamate dehydrogenase 1, mitochondrial precursor | IPI00114209 | 61 kDa | 48 | 19 | 23 | 17 |
| Dlat Dihydrolipoyllysine-residue acetyltransferase component of pyruvate dehydrogenase c | IPI00153660 | 68 kDa | 17 | 12 | 23 | 17 |
| Atp6v1h Vacuolar proton pump subunit H | IPI00311461 | 56 kDa | 8 | 6 | 21 | 17 |
| Ddx5 Probable ATP-dependent RNA helicase DDX5 | IPI00420363 | 69 kDa | 6 | 6 | 21 | 17 |
| Ddx3x ATP-dependent RNA helicase DDX3X | IPI00230035 | 73 kDa | 6 | 5 | 20 | 17 |
| Myh9 Myosin-9 | IPI00123181 (+1) | 226 kDa | 17 | 7 | 30 | 16 |
| Rab10 Ras-related protein Rab-10 | IPI00130118 | 23 kDa | 7 | 3 | 29 | 16 |
| Myrip Rab effector MyRIP | IPI00322509 | 95 kDa | 0 | 0 | 26 | 16 |
| Acly Adult male testis cDNA, RIKEN full-length enriched library, clone:4922505F07 product:A | IPI00126248 (+1) | 121 kDa | 2 | 2 | 25 | 16 |
| Elavl2 ELAV (Embryonic lethal, abnormal vision, Drosophila)-like 2 | IPI00648092 (+1) | 43 kDa | 8 | 7 | 23 | 16 |
| Gls glutaminase isoform 1 | IPI00464317 | 74 kDa | 4 | 4 | 23 | 16 |
| Cdkl5 Isoform 1 of Cyclin-dependent kinase-like 5 | IPI00381416 | 105 kDa | 2 | 2 | 22 | 16 |
| Ubr1 ubiquitin protein ligase E3 component n-recognin 1 | IPI00856455 | 200 kDa | 111 | 43 | 20 | 16 |
| Aldh1l1 10-formyltetrahydrofolate dehydrogenase | IPI00153317 | 99 kDa | 0 | 0 | 19 | 16 |
| Pcx Activated spleen cDNA, RIKEN full-length enriched library, clone:F830201B12 product:py | IPI00114710 | 130 kDa | 13 | 13 | 17 | 16 |
| Madd MAP-kinase activating death domain | IPI00330606 (+4) | 175 kDa | 3 | 3 | 16 | 16 |
| Tubb4 Tubulin beta-4 chain | IPI00109073 | 50 kDa | 677 | 15 | 366 | 15 |
| Glul Glutamine synthetase | IPI00626790 | 42 kDa | 16 | 9 | 29 | 15 |
| Gnb1 Guanine nucleotide-binding protein G(I)/G(S)/G(T) subunit beta-1 | IPI00120716 | 37 kDa | 15 | 9 | 26 | 15 |
| Ssb Lupus La protein homolog | IPI00134300 | 48 kDa | 0 | 0 | 23 | 15 |
| Ap3d1 AP-3 complex subunit delta-1 | IPI00117811 | 135 kDa | 3 | 2 | 21 | 15 |
| Ogdh Isoform 3 of 2-oxoglutarate dehydrogenase E1 component, mitochondrial precursor | IPI00845652 | 118 kDa | 13 | 9 | 17 | 15 |
| Hspa12a Heat shock 70 kDa protein 12A | IPI00279443 | 75 kDa | 6 | 6 | 15 | 15 |
| Pdhb Pyruvate dehydrogenase E1 component subunit beta, mitochondrial precursor | IPI00132042 | 39 kDa | 33 | 13 | 50 | 14 |
| Gnao1 Isoform Alpha-2 of Guanine nucleotide-binding protein G(o) subunit alpha | IPI00115546 | 40 kDa | 31 | 9 | 35 | 14 |
| Atp2a2 Isoform SERCA2B of Sarcoplasmic/endoplasmic reticulum calcium ATPase 2 | IPI00338964 | 115 kDa | 24 | 10 | 31 | 14 |
| Sept3 Isoform 2 of Neuronal-specific septin-3 | IPI00808312 (+1) | 39 kDa | 16 | 7 | 25 | 14 |
| Cap1 Adenylyl cyclase-associated protein 1 | IPI00137331 (+1) | 52 kDa | 7 | 5 | 24 | 14 |
| Crmp1 Crmp1 protein | IPI00312527 | 74 kDa | 15 | 7 | 23 | 14 |
| Tnc Isoform 1 of Tenascin precursor | IPI00403938 (+3) | 232 kDa | 0 | 0 | 23 | 14 |
| Syn3 Synapsin-3 | IPI00463761 | 63 kDa | 0 | 0 | 22 | 14 |
| Hnrnpul1 Isoform 1 of Heterogeneous nuclear ribonucleoprotein U-like protein 1 | IPI00123501 | 96 kDa | 0 | 0 | 19 | 14 |
| Clip2 Isoform 1 of CAP-Gly domain-containing linker protein 2 | IPI00467854 | 116 kDa | 40 | 33 | 18 | 14 |
| Hadhb Trifunctional enzyme subunit beta, mitochondrial precursor | IPI00115607 | 51 kDa | 4 | 4 | 18 | 14 |

| | | | | | | |
|---|---|---|---|---|---|---|
| Canx Calnexin precursor | IPI00119618 | 67 kDa | 19 | 14 | 17 | 14 |
| Birc6 baculoviral IAP repeat-containing 6 | IPI00134095 | 529 kDa | 0 | 0 | 17 | 14 |
| Mgea5 Isoform 3 of Bifunctional protein NCOAT | IPI00406624 (+1) | 107 kDa | 0 | 0 | 17 | 14 |
| Lmna Isoform A of Lamin-A/C | IPI00620256 | 74 kDa | 15 | 14 | 16 | 14 |
| Sdha Succinate dehydrogenase [ubiquinone] flavoprotein subunit, mitochondrial precursor | IPI00230351 | 73 kDa | 5 | 5 | 16 | 14 |
| Plxna4 212 kDa protein | IPI00876097 | 212 kDa | 0 | 0 | 16 | 14 |
| Ubr2 Isoform 1 of E3 ubiquitin-protein ligase UBR2 | IPI00468701 (+1) | 199 kDa | 52 | 24 | 15 | 14 |
| Opa1 Isoform 1 of Dynamin-like 120 kDa protein, mitochondrial precursor | IPI00117657 (+1) | 111 kDa | 19 | 18 | 15 | 14 |
| Cct4 T-complex protein 1 subunit delta | IPI00116277 | 58 kDa | 9 | 7 | 15 | 14 |
| Plg Plasminogen precursor | IPI00322936 | 91 kDa | 40 | 26 | 14 | 14 |
| Slc1a2 Isoform Glt-1 of Excitatory amino acid transporter 2 | IPI00470184 | 62 kDa | 70 | 9 | 73 | 13 |
| Spire2 Isoform 1 of Protein spire homolog 2 | IPI00173139 | 80 kDa | 0 | 0 | 29 | 13 |
| Sept8 Septin 8 | IPI00875717 | 56 kDa | 21 | 11 | 26 | 13 |
| Necab1 N-terminal EF-hand calcium-binding protein 1 | IPI00221430 | 41 kDa | 0 | 0 | 25 | 13 |
| Snap25 Isoform SNAP-25b of Synaptosomal-associated protein 25 | IPI00125635 | 23 kDa | 32 | 14 | 23 | 13 |
| Pura;LOC100045958 Transcriptional activator protein Pur-alpha | IPI00118447 | 35 kDa | 8 | 7 | 22 | 13 |
| Hdac11 Histone deacetylase 11 | IPI00127900 | 39 kDa | 0 | 0 | 22 | 13 |
| Ina Alpha-internexin | IPI00135965 | 56 kDa | 39 | 15 | 21 | 13 |
| Bsn Isoform 1 of Protein bassoon | IPI00134093 (+1) | 419 kDa | 11 | 6 | 21 | 13 |
| Cct7 T-complex protein 1 subunit eta | IPI00331174 | 60 kDa | 11 | 8 | 19 | 13 |
| Sf3b2 Bone marrow macrophage cDNA, RIKEN full-length enriched library, clone:I830028H0 | IPI00349401 | 98 kDa | 4 | 4 | 19 | 13 |
| Hnrnpa2b1 Isoform 1 of Heterogeneous nuclear ribonucleoproteins A2/B1 | IPI00828488 | 37 kDa | 2 | 2 | 19 | 13 |
| Aldoc Fructose-bisphosphate aldolase C | IPI00119458 | 39 kDa | 7 | 7 | 18 | 13 |
| Cct6a T-complex protein 1 subunit zeta | IPI00116281 | 58 kDa | 5 | 5 | 18 | 13 |
| Camkv CaM kinase-like vesicle-associated protein | IPI00122486 | 55 kDa | 17 | 13 | 17 | 13 |
| Cct2 T-complex protein 1 subunit beta | IPI00320217 | 57 kDa | 12 | 11 | 17 | 13 |
| Nckap1 Isoform 1 of Nck-associated protein 1 | IPI00319320 (+3) | 129 kDa | 7 | 4 | 17 | 13 |
| Rps3 40S ribosomal protein S3 | IPI00134599 | 27 kDa | 2 | 2 | 17 | 13 |
| Csnk2a1;ENSMUSG00000057808 casein kinase 2, alpha 1 polypeptide | IPI00408176 | 45 kDa | 2 | 2 | 16 | 13 |
| Psmd2 26S proteasome non-ATPase regulatory subunit 2 | IPI00123494 | 100 kDa | 21 | 16 | 15 | 13 |
| Psmd1 26S proteasome non-ATPase regulatory subunit 1 | IPI00267295 (+1) | 106 kDa | 6 | 6 | 15 | 13 |
| Hspe1 10 kDa heat shock protein, mitochondrial | IPI00263863 | 11 kDa | 8 | 4 | 66 | 12 |
| Add1 Isoform 1 of Alpha-adducin | IPI00136000 (+3) | 81 kDa | 29 | 14 | 23 | 12 |
| Mdh2 Malate dehydrogenase, mitochondrial precursor | IPI00323592 | 36 kDa | 10 | 9 | 23 | 12 |
| Ywhaz 14-3-3 protein zeta/delta | IPI00116498 | 28 kDa | 31 | 12 | 21 | 12 |
| Gnb2l1 Guanine nucleotide-binding protein subunit beta-2-like 1 | IPI00317740 | 35 kDa | 2 | 2 | 21 | 12 |
| Amph Amphiphysin | IPI00400180 | 75 kDa | 30 | 18 | 20 | 12 |
| Ckb Creatine kinase B-type | IPI00136703 | 43 kDa | 9 | 7 | 20 | 12 |
| Agrn Agrin | IPI00378698 (+2) | 205 kDa | 0 | 0 | 18 | 12 |
| Rph3a Rabphilin-3A | IPI00111151 | 75 kDa | 9 | 8 | 16 | 12 |
| Prmt5 Protein arginine N-methyltransferase 5 | IPI00229845 | 73 kDa | 14 | 10 | 15 | 12 |

| | | | | | | |
|---|---|---|---|---|---|---|
| Got2 Aspartate aminotransferase, mitochondrial precursor | IPI00117312 | 47 kDa | 5 | 4 | 15 | 12 |
| Gramd1b Isoform 4 of GRAM domain-containing protein 1B | IPI00227757 (+2) | 101 kDa | 0 | 0 | 15 | 12 |
| Actn1 Alpha-actinin-1 | IPI00380436 | 103 kDa | 23 | 19 | 14 | 12 |
| Alb Serum albumin precursor | IPI00131695 | 69 kDa | 15 | 13 | 14 | 12 |
| Ppp1ca Serine/threonine-protein phosphatase PP1-alpha catalytic subunit | IPI00130185 | 38 kDa | 17 | 12 | 14 | 12 |
| Rps4x 40S ribosomal protein S4, X isoform | IPI00331092 | 30 kDa | 8 | 5 | 14 | 12 |
| Tufm Isoform 1 of Elongation factor Tu, mitochondrial precursor | IPI00274407 (+1) | 50 kDa | 20 | 14 | 13 | 12 |
| Uso1 Isoform 1 of General vesicular transport factor p115 | IPI00128071 | 107 kDa | 6 | 6 | 13 | 12 |
| Vps53 Isoform 1 of Vacuolar protein sorting-associated protein 53 homolog | IPI00387427 | 94 kDa | 0 | 0 | 13 | 12 |
| Psmd3 26S proteasome non-ATPase regulatory subunit 3 | IPI00314439 (+1) | 61 kDa | 0 | 0 | 13 | 12 |
| Acsl6 acyl-CoA synthetase long-chain family member 6 isoform 4 | IPI00262693 (+1) | 78 kDa | 0 | 0 | 13 | 12 |
| Frap1 Isoform 1 of FKBP12-rapamycin complex-associated protein | IPI00268673 | 289 kDa | 0 | 0 | 12 | 12 |
| Tubb3 Tubulin beta-3 chain | IPI00112251 | 50 kDa | 471 | 13 | 273 | 11 |
| Cspg5 Isoform 1 of Chondroitin sulfate proteoglycan 5 precursor | IPI00454159 | 60 kDa | 26 | 8 | 72 | 11 |
| Eef1a1 Elongation factor 1-alpha 1 | IPI00307837 | 50 kDa | 31 | 7 | 33 | 11 |
| Sept6 Isoform V of Septin-6 | IPI00226602 (+2) | 49 kDa | 14 | 5 | 31 | 11 |
| Atp2b1 plasma membrane calcium ATPase 1 | IPI00556827 | 135 kDa | 17 | 6 | 27 | 11 |
| Rchy1 RING finger and CHY zinc finger domain-containing protein 1 | IPI00133161 | 30 kDa | 9 | 6 | 25 | 11 |
| Ncdn Isoform 1 of Neurochondrin | IPI00331299 (+1) | 79 kDa | 13 | 5 | 25 | 11 |
| Ybx1 Nuclease-sensitive element-binding protein 1 | IPI00120886 (+1) | 36 kDa | 0 | 0 | 23 | 11 |
| Actr1a Alpha-centractin | IPI00113895 | 43 kDa | 9 | 8 | 20 | 11 |
| Mapk1 Mitogen-activated protein kinase 1 | IPI00119663 | 41 kDa | 0 | 0 | 18 | 11 |
| Trove2 60 kDa SS-A/Ro ribonucleoprotein | IPI00116360 | 60 kDa | 0 | 0 | 18 | 11 |
| Capzb Isoform 2 of F-actin-capping protein subunit beta | IPI00269481 (+2) | 31 kDa | 9 | 5 | 17 | 11 |
| Prkaca Isoform 1 of cAMP-dependent protein kinase catalytic subunit alpha | IPI00230005 | 41 kDa | 6 | 5 | 17 | 11 |
| Bdh1 3-hydroxybutyrate dehydrogenase, type 1 | IPI00890322 | 38 kDa | 27 | 12 | 16 | 11 |
| Capza2 F-actin-capping protein subunit alpha-2 | IPI00111265 | 33 kDa | 10 | 8 | 15 | 11 |
| Ptrf Polymerase I and transcript release factor | IPI00117689 | 44 kDa | 0 | 0 | 15 | 11 |
| Prkar2b cAMP-dependent protein kinase type II-beta regulatory subunit | IPI00224570 | 46 kDa | 5 | 5 | 14 | 11 |
| Pcca Propionyl-CoA carboxylase alpha chain, mitochondrial precursor | IPI00330523 | 80 kDa | 3 | 3 | 14 | 11 |
| Add2 Isoform 1 of Beta-adducin | IPI00323122 | 81 kDa | 19 | 14 | 13 | 11 |
| LOC100046594 similar to heterogeneous nuclear ribonucleoprotein U-like 2 | IPI00849047 | 85 kDa | 7 | 7 | 13 | 11 |
| Atp6v1c1 Vacuolar proton pump subunit C 1 | IPI00130186 | 44 kDa | 4 | 4 | 13 | 11 |
| Eprs Bifunctional aminoacyl-tRNA synthetase | IPI00339916 (+1) | 170 kDa | 3 | 3 | 13 | 11 |
| Ap3b2 Isoform 1 of AP-3 complex subunit beta-2 | IPI00420426 | 119 kDa | 15 | 11 | 12 | 11 |
| Slc3a2 CD98 heavy chain | IPI00114641 | 59 kDa | 14 | 11 | 12 | 11 |
| Vars Valyl-tRNA synthetase | IPI00130353 | 140 kDa | 5 | 5 | 12 | 11 |
| Dnm1l Isoform 2 of Dynamin-1-like protein | IPI00172221 (+3) | 80 kDa | 3 | 3 | 11 | 11 |
| Ide insulin degrading enzyme | IPI00828796 | 118 kDa | 0 | 0 | 11 | 11 |
| Fus RNA-binding protein FUS | IPI00117063 (+1) | 53 kDa | 48 | 11 | 42 | 10 |
| Sv2a Synaptic vesicle glycoprotein 2A | IPI00465810 | 83 kDa | 40 | 9 | 38 | 10 |

| | | | | | | |
|---|---|---|---|---|---|---|
| Camk2b Calcium/calmodulin-dependent protein kinase type II beta chain | IPI00474502 (+2) | 60 kDa | 39 | 9 | 31 | 10 |
| Mtap4 Isoform 1 of Microtubule-associated protein 4 | IPI00408119 (+1) | 117 kDa | 19 | 11 | 29 | 10 |
| Ldhb L-lactate dehydrogenase B chain | IPI00229510 | 37 kDa | 7 | 6 | 26 | 10 |
| Dnaja1 DnaJ homolog subfamily A member 1 | IPI00132208 (+1) | 45 kDa | 22 | 11 | 24 | 10 |
| #NAME? | IPI00831055 | 16 kDa | 23 | 10 | 19 | 10 |
| Cct5 T-complex protein 1 subunit epsilon | IPI00116279 | 60 kDa | 12 | 8 | 18 | 10 |
| Dld Dihydrolipoyl dehydrogenase, mitochondrial precursor | IPI00874456 | 54 kDa | 3 | 3 | 18 | 10 |
| Hnrnpc Bone marrow macrophage cDNA, RIKEN full-length enriched library, clone:I830120C | IPI00130343 (+2) | 37 kDa | 10 | 8 | 17 | 10 |
| Ppp2ca Serine/threonine-protein phosphatase 2A catalytic subunit alpha isoform | IPI00120374 | 36 kDa | 12 | 9 | 15 | 10 |
| Ogdhl oxoglutarate dehydrogenase-like | IPI00342603 | 117 kDa | 9 | 2 | 15 | 10 |
| Kpnb1 Importin subunit beta-1 | IPI00323881 (+1) | 97 kDa | 14 | 12 | 14 | 10 |
| Gfap Isoform 1 of Glial fibrillary acidic protein | IPI00117042 (+1) | 50 kDa | 15 | 8 | 14 | 10 |
| Rock2 Rho-associated protein kinase 2 | IPI00108150 (+1) | 161 kDa | 7 | 5 | 14 | 10 |
| Sh3gl2 Endophilin-A1 | IPI00331110 (+1) | 40 kDa | 4 | 4 | 14 | 10 |
| Hnrnpl heterogeneous nuclear ribonucleoprotein L | IPI00620362 | 64 kDa | 0 | 0 | 14 | 10 |
| Rilpl2 Isoform 1 of RILP-like protein 2 | IPI00116819 | 22 kDa | 0 | 0 | 14 | 10 |
| Hnrnpk Isoform 1 of Heterogeneous nuclear ribonucleoprotein K | IPI00223253 (+2) | 51 kDa | 24 | 14 | 13 | 10 |
| Upf1 Isoform 1 of Regulator of nonsense transcripts 1 | IPI00420949 (+1) | 124 kDa | 5 | 5 | 13 | 10 |
| Dnajc6 Isoform 1 of Putative tyrosine-protein phosphatase auxilin | IPI00330269 (+2) | 102 kDa | 17 | 11 | 12 | 10 |
| Myo18a Isoform 3 of Myosin-XVIIIa | IPI00619995 (+3) | 233 kDa | 11 | 9 | 12 | 10 |
| BC010304 hypothetical protein LOC218236 | IPI00830478 | 122 kDa | 8 | 7 | 12 | 10 |
| Psmd6 26S proteasome non-ATPase regulatory subunit 6 | IPI00319965 | 46 kDa | 7 | 6 | 12 | 10 |
| Psmc6 26S protease regulatory subunit S10B | IPI00125971 | 44 kDa | 16 | 10 | 11 | 10 |
| Adam23 Isoform Alpha of ADAM 23 precursor | IPI00129175 (+3) | 92 kDa | 5 | 5 | 11 | 10 |
| Mink1 misshapen-like kinase 1 isoform 2 | IPI00124753 (+1) | 147 kDa | 5 | 4 | 11 | 10 |
| LOC100045332;Rpsa 40S ribosomal protein SA | IPI00123604 (+1) | 33 kDa | 3 | 3 | 11 | 10 |
| 1110014N23Rik CRL-1722 L5178Y-R cDNA, RIKEN full-length enriched library, clone:I730034 | IPI00652882 | 86 kDa | 0 | 0 | 11 | 10 |
| Sorl1 Adult male brain UNDEFINED_CELL_LINE cDNA, RIKEN full-length enriched library, clor | IPI00776230 | 247 kDa | 3 | 3 | 10 | 10 |
| Eif4g1 Isoform 1 of Eukaryotic translation initiation factor 4 gamma 1 | IPI00421179 (+2) | 176 kDa | 0 | 0 | 10 | 10 |
| Slc1a3 Excitatory amino acid transporter 1 | IPI00114279 | 60 kDa | 43 | 5 | 54 | 9 |
| Pabpc4 Poly A binding protein, cytoplasmic 4 | IPI00172364 (+3) | 68 kDa | 0 | 0 | 31 | 9 |
| Hspa1b Heat shock 70 kDa protein 1B | IPI00346073 (+1) | 70 kDa | 42 | 14 | 21 | 9 |
| Rab3a Ras-related protein Rab-3A | IPI00122965 | 25 kDa | 12 | 6 | 20 | 9 |
| Dpysl3 Dihydropyrimidinase-related protein 3 | IPI00122349 (+2) | 62 kDa | 0 | 0 | 20 | 9 |
| Ywhag 14-3-3 protein gamma | IPI00230707 | 28 kDa | 25 | 6 | 19 | 9 |
| Krt17 Keratin, type I cytoskeletal 17 | IPI00230365 | 48 kDa | 20 | 4 | 19 | 9 |
| Tardbp TAR DNA-binding protein 43 | IPI00121758 | 45 kDa | 4 | 3 | 19 | 9 |
| Pclo piccolo isoform 1 | IPI00225140 (+4) | 551 kDa | 29 | 16 | 17 | 9 |
| Psma1 Proteasome subunit alpha type-1 | IPI00283862 | 30 kDa | 21 | 7 | 17 | 9 |
| Kif5c Visual cortex cDNA, RIKEN full-length enriched library, clone:K430357F23 product:kine | IPI00753326 | 109 kDa | 14 | 12 | 15 | 9 |
| Atp6v1e1 Vacuolar proton pump subunit E 1 | IPI00119115 | 26 kDa | 24 | 9 | 15 | 9 |

| | | | | | | |
|---|---|---|---|---|---|---|
| Vps35 Vacuolar protein sorting-associated protein 35 | IPI00111181 | 92 kDa | 4 | 4 | 15 | 9 |
| Lancl2 LanC-like protein 2 | IPI00121949 (+1) | 51 kDa | 0 | 0 | 15 | 9 |
| Sfpq Splicing factor, proline- and glutamine-rich | IPI00129430 (+1) | 75 kDa | 19 | 15 | 14 | 9 |
| Tom1;EG545878 Isoform 1 of Target of Myb protein 1 | IPI00380814 | 54 kDa | 233 | 31 | 13 | 9 |
| Aak1 Isoform 2 of AP2-associated protein kinase 1 | IPI00356608 (+1) | 95 kDa | 27 | 19 | 13 | 9 |
| Actr2 Actin-related protein 2 | IPI00177038 | 45 kDa | 2 | 2 | 13 | 9 |
| Clip1 Isoform 1 of CAP-Gly domain-containing linker protein 1 | IPI00123063 (+4) | 156 kDa | 17 | 8 | 12 | 9 |
| Idh3a Isoform 1 of Isocitrate dehydrogenase [NAD] subunit alpha, mitochondrial precursor | IPI00459725 | 40 kDa | 6 | 6 | 12 | 9 |
| Caskin1 Isoform 1 of Caskin-1 | IPI00411109 (+2) | 150 kDa | 3 | 3 | 12 | 9 |
| Fem1c Protein fem-1 homolog C | IPI00378461 | 69 kDa | 0 | 0 | 12 | 9 |
| Ctnnb1 Catenin beta-1 | IPI00125899 (+1) | 85 kDa | 19 | 13 | 11 | 9 |
| Hsp90b1 Endoplasmin precursor | IPI00129526 | 92 kDa | 14 | 11 | 11 | 9 |
| Dclk1 Isoform 1 of Serine/threonine-protein kinase DCLK1 | IPI00468380 | 84 kDa | 6 | 6 | 11 | 9 |
| Fech Ferrochelatase | IPI00228343 (+1) | 47 kDa | 7 | 5 | 11 | 9 |
| Raly Isoform 2 of RNA-binding protein Raly | IPI00130147 (+3) | 33 kDa | 3 | 3 | 11 | 9 |
| Cntnap1 Contactin-associated protein 1 precursor | IPI00338983 (+1) | 156 kDa | 0 | 0 | 11 | 9 |
| Lmnb1 Lamin-B1 | IPI00230394 | 67 kDa | 8 | 8 | 10 | 9 |
| Klc2 kinesin light chain 2 | IPI00753928 (+1) | 68 kDa | 7 | 7 | 10 | 9 |
| Kars Lysyl-tRNA synthetase | IPI00620145 (+1) | 68 kDa | 5 | 4 | 10 | 9 |
| Eef2 Elongation factor 2 | IPI00466069 | 95 kDa | 4 | 4 | 10 | 9 |
| Psmd12 Bone marrow macrophage cDNA, RIKEN full-length enriched library, clone:I830057( | IPI00133066 | 53 kDa | 4 | 3 | 10 | 9 |
| Acad9 very-long-chain acyl-CoA dehydrogenase VLCAD homolog | IPI00759881 | 69 kDa | 0 | 0 | 10 | 9 |
| Acsbg1 Long-chain-fatty-acid--CoA ligase ACSBG1 | IPI00453834 | 80 kDa | 0 | 0 | 10 | 9 |
| Immt Isoform 1 of Mitochondrial inner membrane protein | IPI00228150 (+2) | 84 kDa | 24 | 17 | 9 | 9 |
| Mccc1 Methylcrotonoyl-CoA carboxylase subunit alpha, mitochondrial precursor | IPI00320850 | 79 kDa | 2 | 2 | 9 | 9 |
| Homer1 Isoform 2 of Homer protein homolog 1 | IPI00172234 (+2) | 42 kDa | 0 | 0 | 9 | 9 |
| Dars Aspartyl-tRNA synthetase, cytoplasmic | IPI00122743 (+1) | 57 kDa | 0 | 0 | 9 | 9 |
| Dnajc10 DnaJ (Hsp40) homolog, subfamily C, member 10 | IPI00229600 | 91 kDa | 0 | 0 | 9 | 9 |
| Atp1a2 Sodium/potassium-transporting ATPase subunit alpha-2 precursor | IPI00420569 (+1) | 112 kDa | 54 | 3 | 80 | 8 |
| Atp1a1 Sodium/potassium-transporting ATPase subunit alpha-1 precursor | IPI00311682 | 113 kDa | 46 | 6 | 70 | 8 |
| Krt5 Keratin, type II cytoskeletal 5 | IPI00139301 (+1) | 62 kDa | 45 | 3 | 33 | 8 |
| Ddx17 Isoform 1 of Probable ATP-dependent RNA helicase DDX17 | IPI00396797 (+1) | 72 kDa | 5 | 2 | 21 | 8 |
| Hnrnph1 Heterogeneous nuclear ribonucleoprotein H | IPI00133916 (+1) | 49 kDa | 10 | 6 | 20 | 8 |
| Purb Transcriptional activator protein Pur-beta | IPI00128867 | 34 kDa | 5 | 3 | 17 | 8 |
| Macf1 Microtubule-actin crosslinking factor 1b | IPI00884482 | 832 kDa | 15 | 8 | 15 | 8 |
| Luc7l2 Isoform 1 of Putative RNA-binding protein Luc7-like 2 | IPI00380309 (+2) | 47 kDa | 4 | 4 | 15 | 8 |
| Ahcyl2 Isoform 1 of Putative adenosylhomocysteinase 3 | IPI00308446 (+1) | 67 kDa | 44 | 15 | 14 | 8 |
| Psmb5 Proteasome subunit beta type-5 precursor | IPI00317902 | 29 kDa | 19 | 11 | 14 | 8 |
| Rps8;ENSMUSG00000070446 40S ribosomal protein S8 | IPI00466820 | 24 kDa | 2 | 2 | 14 | 8 |
| Rasgrf1 Ras-specific guanine nucleotide-releasing factor 1 | IPI00118835 | 144 kDa | 0 | 0 | 14 | 8 |
| Kcnab2 Voltage-gated potassium channel subunit beta-2 | IPI00315359 | 41 kDa | 10 | 5 | 13 | 8 |

| | | | | | | |
|---|---|---|---|---|---|---|
| Elavl1 ELAV (embryonic lethal, abnormal vision, Drosophila)-like 1 | IPI00466032 | 36 kDa | 4 | 4 | 13 | 8 |
| Ywhaq Isoform 1 of 14-3-3 protein theta | IPI00408378 (+2) | 28 kDa | 19 | 6 | 12 | 8 |
| Eef1g Elongation factor 1-gamma | IPI00318841 | 50 kDa | 6 | 5 | 12 | 8 |
| Gnb5 Isoform 2 of Guanine nucleotide-binding protein subunit beta-5 | IPI00378017 | 39 kDa | 6 | 4 | 12 | 8 |
| Crbn cereblon isoform 1 | IPI00123335 (+1) | 51 kDa | 0 | 0 | 12 | 8 |
| Poldip3 Polymerase delta-interacting protein 3 | IPI00229721 (+1) | 46 kDa | 0 | 0 | 12 | 8 |
| Psma7 Proteasome subunit alpha type-7 | IPI00131406 | 28 kDa | 16 | 8 | 11 | 8 |
| Rbm14 Isoform 1 of RNA-binding protein 14 | IPI00404707 | 69 kDa | 8 | 7 | 11 | 8 |
| Tpm3 Isoform 2 of Tropomyosin alpha-3 chain | IPI00230044 | 29 kDa | 7 | 5 | 11 | 8 |
| Wdfy3 Isoform 1 of WD repeat and FYVE domain-containing protein 3 | IPI00227110 (+1) | 392 kDa | 3 | 3 | 11 | 8 |
| Spock2 Testican-2 precursor | IPI00111600 | 47 kDa | 3 | 3 | 11 | 8 |
| LOC637606;LOC100043752 hypothetical protein | IPI00604967 (+3) | 30 kDa | 2 | 2 | 11 | 8 |
| Acat1 Acetyl-CoA acetyltransferase, mitochondrial precursor | IPI00154054 | 45 kDa | 2 | 2 | 11 | 8 |
| OTTMUSG00000025606;LOC100044916;Rpl13;OTTMUSG00000009186 60S ribosomal prote | IPI00224505 | 24 kDa | 0 | 0 | 11 | 8 |
| Hspa12b Heat shock 70 kDa protein 12B | IPI00224776 | 76 kDa | 0 | 0 | 11 | 8 |
| Hyou1 Hypoxia up-regulated protein 1 precursor | IPI00123342 | 111 kDa | 13 | 11 | 10 | 8 |
| Wdr77 Methylosome protein 50 | IPI00114819 | 37 kDa | 5 | 4 | 10 | 8 |
| Pde2a Visual cortex cDNA, RIKEN full-length enriched library, clone:K230012M18 product:pl | IPI00380799 (+2) | 106 kDa | 4 | 3 | 10 | 8 |
| Sec23a Protein transport protein Sec23A | IPI00123349 | 86 kDa | 0 | 0 | 10 | 8 |
| Mrps27 Mitochondrial 28S ribosomal protein S27 | IPI00222514 | 48 kDa | 0 | 0 | 10 | 8 |
| Ctnnd2 Isoform 1 of Catenin delta-2 | IPI00136135 (+3) | 135 kDa | 28 | 22 | 9 | 8 |
| Gphn Gphn protein | IPI00816946 | 84 kDa | 7 | 6 | 9 | 8 |
| Hnrnpr Bone marrow macrophage cDNA, RIKEN full-length enriched library, clone:I830045N | IPI00128441 | 71 kDa | 5 | 5 | 9 | 8 |
| Sept2 Septin-2 | IPI00114945 | 42 kDa | 4 | 4 | 9 | 8 |
| Fbxo2 F-box only protein 2 | IPI00153176 (+1) | 34 kDa | 3 | 3 | 9 | 8 |
| Dhx9 Isoform 2 of ATP-dependent RNA helicase A | IPI00339468 (+1) | 150 kDa | 3 | 3 | 9 | 8 |
| Usp24 Ubiquitin carboxyl-terminal hydrolase | IPI00123410 | 294 kDa | 0 | 0 | 9 | 8 |
| Hspg2 Basement membrane-specific heparan sulfate proteoglycan core protein precursor | IPI00113824 (+1) | 398 kDa | 0 | 0 | 9 | 8 |
| Ascc3l1 Activating signal cointegrator 1 complex subunit 3-like 1 | IPI00420329 | 245 kDa | 0 | 0 | 9 | 8 |
| Tpp2 Isoform Short of Tripeptidyl-peptidase 2 | IPI00227843 (+1) | 138 kDa | 0 | 0 | 9 | 8 |
| Matr3 Matrin-3 | IPI00453826 | 95 kDa | 27 | 20 | 8 | 8 |
| Slc25a12 Calcium-binding mitochondrial carrier protein Aralar1 | IPI00308162 | 75 kDa | 7 | 7 | 8 | 8 |
| Psmc2 Adult male testis cDNA, RIKEN full-length enriched library, clone:4930525G09 produc | IPI00270326 | 53 kDa | 3 | 3 | 8 | 8 |
| Ccdc132 Isoform 3 of Coiled-coil domain-containing protein 132 | IPI00869473 | 108 kDa | 0 | 0 | 8 | 8 |
| Herc1 hect (homologous to the E6-AP (UBE3A) carboxyl terminus) domain and RCC1 (CHC1)- | IPI00676574 | 533 kDa | 0 | 0 | 8 | 8 |
| Trim3 Tripartite motif-containing protein 3 | IPI00129020 (+1) | 81 kDa | 0 | 0 | 8 | 8 |
| sp_HBB_HUMAN | IPIsp_HBB_HUMAN | 16 kDa | 28 | 5 | 33 | 7 |
| Hspa2 heat shock protein 2 | IPI00387494 | 70 kDa | 75 | 16 | 28 | 7 |
| LOC100048669;Rps27a;EG619900 ribosomal protein S27a | IPI00470152 | 18 kDa | 82 | 7 | 28 | 7 |
| Dnm3 Isoform 1 of Dynamin-3 | IPI00227432 (+1) | 97 kDa | 13 | 2 | 28 | 7 |
| Epb4.1l2 7 days neonate cerebellum cDNA, RIKEN full-length enriched library, clone:A73006 | IPI00227267 (+5) | 91 kDa | 14 | 5 | 24 | 7 |

| | | | | | | |
|---|---|---|---|---|---|---|
| Clta clathrin, light polypeptide (Lca) isoform c | IPI00648681 (+1) | 26 kDa | 195 | 20 | 20 | 7 |
| Elavl3 Isoform HuC-L of ELAV-like protein 3 | IPI00122451 | 40 kDa | 0 | 0 | 20 | 7 |
| Epb4.1l1 erythrocyte protein band 4.1-like 1 isoform a | IPI00465812 (+1) | 98 kDa | 14 | 6 | 19 | 7 |
| Ywhah 14-3-3 protein eta | IPI00227392 | 28 kDa | 24 | 10 | 13 | 7 |
| Atp5c1 ATP synthase, H+ transporting, mitochondrial F1 complex, gamma subunit isoform b | IPI00750074 (+3) | 30 kDa | 11 | 6 | 13 | 7 |
| Dnaja2 DnaJ homolog subfamily A member 2 | IPI00136251 | 46 kDa | 6 | 5 | 13 | 7 |
| Mdh1 Malate dehydrogenase, cytoplasmic | IPI00336324 | 37 kDa | 4 | 4 | 13 | 7 |
| Lgi1 Leucine-rich glioma-inactivated protein 1 precursor | IPI00120302 | 64 kDa | 4 | 3 | 12 | 7 |
| Nrxn2 neurexin II | IPI00467429 | 184 kDa | 0 | 0 | 12 | 7 |
| Rtn1 Reticulon-1 | IPI00395193 | 84 kDa | 18 | 11 | 11 | 7 |
| Slc25a22 Mitochondrial glutamate carrier 1 | IPI00109275 | 35 kDa | 15 | 8 | 11 | 7 |
| Sept9 Isoform 1 of Septin-9 | IPI00457611 (+2) | 66 kDa | 11 | 8 | 11 | 7 |
| Stx1b Syntaxin-1B | IPI00113149 (+1) | 33 kDa | 31 | 7 | 11 | 7 |
| Psmd11 26S proteasome non-ATPase regulatory subunit 11 | IPI00222515 | 47 kDa | 6 | 5 | 11 | 7 |
| Ckmt1 Creatine kinase, ubiquitous mitochondrial precursor | IPI00128296 | 47 kDa | 4 | 4 | 11 | 7 |
| Ppp3ca Isoform 1 of Serine/threonine-protein phosphatase 2B catalytic subunit alpha isofor | IPI00121545 (+1) | 59 kDa | 8 | 3 | 11 | 7 |
| Kctd12 BTB/POZ domain-containing protein KCTD12 | IPI00421206 | 36 kDa | 3 | 2 | 11 | 7 |
| Rps9 40S ribosomal protein S9 | IPI00420726 | 23 kDa | 0 | 0 | 11 | 7 |
| Vps13a Isoform 1 of Vacuolar protein sorting-associated protein 13A | IPI00420394 (+1) | 359 kDa | 0 | 0 | 11 | 7 |
| Herc2 Isoform 2 of Probable E3 ubiquitin-protein ligase HERC2 | IPI00309574 (+1) | 523 kDa | 0 | 0 | 11 | 7 |
| Cops4 COP9 signalosome complex subunit 4 | IPI00131871 | 46 kDa | 14 | 11 | 10 | 7 |
| D630045J12Rik hypothetical protein LOC330286 | IPI00623897 (+2) | 209 kDa | 3 | 3 | 10 | 7 |
| Plxna1 Plexin-A1 precursor | IPI00137311 | 211 kDa | 0 | 0 | 10 | 7 |
| Strn Striatin, calmodulin binding protein | IPI00554884 | 86 kDa | 38 | 28 | 9 | 7 |
| Slc25a3 Phosphate carrier protein, mitochondrial precursor | IPI00124771 (+1) | 40 kDa | 12 | 6 | 9 | 7 |
| Psma5 Proteasome subunit alpha type-5 | IPI00131407 | 26 kDa | 9 | 6 | 9 | 7 |
| Sgip1 Isoform 1 of SH3-containing GRB2-like protein 3-interacting protein 1 | IPI00465788 (+2) | 86 kDa | 6 | 6 | 9 | 7 |
| Mapre2 Isoform 1 of Microtubule-associated protein RP/EB family member 2 | IPI00403682 (+2) | 37 kDa | 8 | 5 | 9 | 7 |
| Hnrnpu Osteoclast-like cell cDNA, RIKEN full-length enriched library, clone:I420039N16 proc | IPI00458583 | 88 kDa | 6 | 5 | 9 | 7 |
| Csde1 Cold shock domain-containing protein E1 | IPI00274747 | 89 kDa | 2 | 2 | 9 | 7 |
| Prpf8 Pre-mRNA-processing-splicing factor 8 | IPI00121596 (+1) | 274 kDa | 0 | 0 | 9 | 7 |
| Mrps22 Mitochondrial 28S ribosomal protein S22 | IPI00110918 | 41 kDa | 0 | 0 | 9 | 7 |
| Rpl18 60S ribosomal protein L18 | IPI00555113 | 22 kDa | 0 | 0 | 9 | 7 |
| Prmt1 Isoform 1 of Protein arginine N-methyltransferase 1 | IPI00120495 (+3) | 42 kDa | 0 | 0 | 9 | 7 |
| Kif1a Adult male brain UNDEFINED_CELL_LINE cDNA, RIKEN full-length enriched library, clor | IPI00626501 (+1) | 192 kDa | 0 | 0 | 9 | 7 |
| Hnrnpa3 Isoform 1 of Heterogeneous nuclear ribonucleoprotein A3 | IPI00269661 (+1) | 40 kDa | 0 | 0 | 9 | 7 |
| Rabep1 Isoform 1 of Rab GTPase-binding effector protein 1 | IPI00131692 (+4) | 100 kDa | 16 | 13 | 8 | 7 |
| Psma3 Proteasome subunit alpha type-3 | IPI00331644 | 28 kDa | 15 | 8 | 8 | 7 |
| Pacsin1 Protein kinase C and casein kinase substrate in neurons protein 1 | IPI00123613 | 51 kDa | 9 | 8 | 8 | 7 |
| Akap12 Isoform 1 of A-kinase anchor protein 12 | IPI00123709 (+1) | 181 kDa | 8 | 7 | 8 | 7 |
| Gprin1 G protein-regulated inducer of neurite outgrowth 1 | IPI00138232 | 95 kDa | 8 | 6 | 8 | 7 |

| | | | | | | |
|---|---|---|---|---|---|---|
| Sirt2 Isoform 1 of NAD-dependent deacetylase sirtuin-2 | IPI00110265 (+1) | 43 kDa | 6 | 6 | 8 | 7 |
| Aifm1 Apoptosis-inducing factor 1, mitochondrial precursor | IPI00129577 (+1) | 67 kDa | 4 | 4 | 8 | 7 |
| Adam22 Isoform 19 of ADAM 22 precursor | IPI00378796 (+20) | 94 kDa | 3 | 2 | 8 | 7 |
| Phgdh D-3-phosphoglycerate dehydrogenase | IPI00225961 | 57 kDa | 2 | 2 | 8 | 7 |
| Ptprd protein tyrosine phosphatase, receptor type, D isoform B | IPI00465836 (+2) | 170 kDa | 0 | 0 | 8 | 7 |
| Cspg4 Isoform 1 of Chondroitin sulfate proteoglycan 4 precursor | IPI00128915 | 252 kDa | 0 | 0 | 8 | 7 |
| 1300010F03Rik hypothetical protein LOC219189 | IPI00125180 (+1) | 213 kDa | 0 | 0 | 8 | 7 |
| Dpp9 Isoform 1 of Dipeptidyl peptidase 9 | IPI00462461 | 98 kDa | 0 | 0 | 8 | 7 |
| Pex5l 73 kDa protein | IPI00875944 | 73 kDa | 34 | 23 | 7 | 7 |
| sp_KRHB6_HUMAN | IPIsp_KRHB6_HUMAN | 53 kDa | 9 | 8 | 7 | 7 |
| Psmd13 Isoform 1 of 26S proteasome non-ATPase regulatory subunit 13 | IPI00126048 | 43 kDa | 8 | 8 | 7 | 7 |
| Cfl1 Cofilin-1 | IPI00890117 | 19 kDa | 8 | 7 | 7 | 7 |
| Cask Isoform 5 of Peripheral plasma membrane protein CASK | IPI00119517 (+3) | 104 kDa | 8 | 7 | 7 | 7 |
| Wasf1 Wiskott-Aldrich syndrome protein family member 1 | IPI00471372 | 62 kDa | 6 | 6 | 7 | 7 |
| EG433182;Eno1;LOC100044223 Alpha-enolase | IPI00462072 | 47 kDa | 5 | 4 | 7 | 7 |
| Wdr48 Isoform 1 of WD repeat-containing protein 48 | IPI00336198 (+1) | 76 kDa | 2 | 2 | 7 | 7 |
| Copb1 Coatomer subunit beta | IPI00120503 | 107 kDa | 2 | 2 | 7 | 7 |
| Tssc1 Protein TSSC1 | IPI00321310 | 43 kDa | 0 | 0 | 7 | 7 |
| Sec24c SEC24 related gene family, member C | IPI00229483 (+1) | 119 kDa | 0 | 0 | 7 | 7 |
| Usp7 Isoform 2 of Ubiquitin carboxyl-terminal hydrolase 7 | IPI00463367 (+2) | 133 kDa | 0 | 0 | 7 | 7 |
| Poldip2 Polymerase delta-interacting protein 2 | IPI00126634 | 42 kDa | 0 | 0 | 7 | 7 |
| Kank3 KN motif and ankyrin repeat domain-containing protein 3 | IPI00130324 | 84 kDa | 0 | 0 | 7 | 7 |
| Spire1 MKIAA1135 protein (Fragment) | IPI00460722 | 44 kDa | 0 | 0 | 131 | 6 |
| Slc25a5 ADP/ATP translocase 2 | IPI00127841 | 33 kDa | 36 | 5 | 29 | 6 |
| Ywhab Isoform Long of 14-3-3 protein beta/alpha | IPI00230682 | 28 kDa | 25 | 5 | 15 | 6 |
| Acan Aggrecan core protein precursor | IPI00119035 | 222 kDa | 0 | 0 | 15 | 6 |
| Pabpn1 Isoform 1 of Polyadenylate-binding protein 2 | IPI00136169 (+1) | 32 kDa | 0 | 0 | 14 | 6 |
| Bin1 Isoform 1 of Myc box-dependent-interacting protein 1 | IPI00114352 | 64 kDa | 20 | 9 | 11 | 6 |
| Vdac1 Isoform Pl-VDAC1 of Voltage-dependent anion-selective channel protein 1 | IPI00122549 (+1) | 32 kDa | 7 | 6 | 11 | 6 |
| Gnai1 guanine nucleotide binding protein, alpha inhibiting 1 | IPI00467152 | 40 kDa | 10 | 3 | 11 | 6 |
| Ckap5 Cytoskeleton associated protein 5 | IPI00317134 (+2) | 219 kDa | 5 | 3 | 11 | 6 |
| Idh3g Isocitrate dehydrogenase [NAD] subunit gamma, mitochondrial precursor | IPI00109169 | 43 kDa | 4 | 4 | 10 | 6 |
| Nptn 44 kDa protein | IPI00757771 | 44 kDa | 4 | 3 | 10 | 6 |
| Igsf8 Immunoglobulin superfamily member 8 precursor | IPI00321348 | 65 kDa | 0 | 0 | 10 | 6 |
| Msra Peptide methionine sulfoxide reductase | IPI00331442 | 26 kDa | 3 | 3 | 9 | 6 |
| Aco2 Aconitate hydratase, mitochondrial precursor | IPI00116074 | 85 kDa | 3 | 2 | 9 | 6 |
| Acaca Isoform 1 of Acetyl-CoA carboxylase 1 | IPI00474783 (+1) | 265 kDa | 2 | 2 | 9 | 6 |
| Pccb Propionyl Coenzyme A carboxylase, beta polypeptide | IPI00468653 (+1) | 58 kDa | 0 | 0 | 9 | 6 |
| Dab1 Isoform DAB588 of Disabled homolog 1 | IPI00123898 (+2) | 64 kDa | 16 | 7 | 8 | 6 |
| Psmb1 Proteasome subunit beta type-1 precursor | IPI00113845 | 26 kDa | 14 | 7 | 8 | 6 |
| G3bp2 Isoform A of Ras GTPase-activating protein-binding protein 2 | IPI00124245 | 54 kDa | 9 | 7 | 8 | 6 |

| | | | | | | |
|---|---|---|---|---|---|---|
| Prdx1 Peroxiredoxin-1 | IPI00121788 | 22 kDa | 9 | 7 | 8 | 6 |
| Hspd1 Isoform 1 of 60 kDa heat shock protein, mitochondrial precursor | IPI00308885 | 61 kDa | 12 | 6 | 8 | 6 |
| Psmd7 26S proteasome non-ATPase regulatory subunit 7 | IPI00114667 | 37 kDa | 8 | 5 | 8 | 6 |
| Eef1d eukaryotic translation elongation factor 1 delta isoform b | IPI00515654 | 31 kDa | 7 | 5 | 8 | 6 |
| Syt1 Synaptotagmin-1 | IPI00129618 (+1) | 47 kDa | 4 | 3 | 8 | 6 |
| Tceb2 Transcription elongation factor B polypeptide 2 | IPI00131224 | 13 kDa | 3 | 3 | 8 | 6 |
| Prkcc Protein kinase C gamma type | IPI00122069 | 78 kDa | 4 | 2 | 8 | 6 |
| Rps11 Adult male cerebellum cDNA, RIKEN full-length enriched library, clone:1500004H08 p | IPI00117569 (+1) | 19 kDa | 0 | 0 | 8 | 6 |
| Cul2 Isoform 1 of Cullin-2 | IPI00387216 | 87 kDa | 0 | 0 | 8 | 6 |
| Rbm39 Isoform 1 of RNA-binding protein 39 | IPI00127763 (+2) | 59 kDa | 0 | 0 | 8 | 6 |
| Cap2 Adenylyl cyclase-associated protein 2 | IPI00112001 | 53 kDa | 0 | 0 | 8 | 6 |
| Strn3 Striatin-3 | IPI00229990 | 87 kDa | 57 | 21 | 7 | 6 |
| Tollip Toll-interacting protein | IPI00136618 | 30 kDa | 95 | 15 | 7 | 6 |
| AW555464 Isoform 1 of Protein KIAA0284 | IPI00380953 | 171 kDa | 9 | 8 | 7 | 6 |
| Dctn2 Dynactin subunit 2 | IPI00116112 | 44 kDa | 9 | 6 | 7 | 6 |
| Sirpa 56 kDa protein | IPI00775779 (+1) | 56 kDa | 8 | 6 | 7 | 6 |
| Necab2 Isoform 1 of N-terminal EF-hand calcium-binding protein 2 | IPI00131287 | 43 kDa | 7 | 6 | 7 | 6 |
| Atp6v1d Vacuolar proton pump subunit D | IPI00118787 | 28 kDa | 10 | 5 | 7 | 6 |
| Rplp0 60S acidic ribosomal protein P0 | IPI00314950 | 34 kDa | 5 | 5 | 7 | 6 |
| Cops5 COP9 signalosome complex subunit 5 | IPI00135087 | 38 kDa | 5 | 5 | 7 | 6 |
| Trio triple functional domain | IPI00604947 (+1) | 348 kDa | 3 | 2 | 7 | 6 |
| LOC100048637;Tmod2 Isoform 1 of Tropomodulin-2 | IPI00402982 | 40 kDa | 2 | 2 | 7 | 6 |
| Map2k1 Dual specificity mitogen-activated protein kinase kinase 1 | IPI00466610 | 43 kDa | 2 | 2 | 7 | 6 |
| - 42 kDa protein | IPI00474144 (+1) | 42 kDa | 0 | 0 | 7 | 6 |
| Itih5 Inter-alpha-trypsin inhibitor heavy chain H5 precursor | IPI00222369 | 107 kDa | 0 | 0 | 7 | 6 |
| Arpc1a Actin-related protein 2/3 complex subunit 1A | IPI00127987 | 42 kDa | 0 | 0 | 7 | 6 |
| Dock7 Isoform 1 of Dedicator of cytokinesis protein 7 | IPI00816914 | 241 kDa | 89 | 32 | 6 | 6 |
| Dbn1 Isoform A of Drebrin | IPI00135475 (+1) | 77 kDa | 11 | 9 | 6 | 6 |
| Sucla2 Succinyl-CoA ligase [ADP-forming] subunit beta, mitochondrial precursor | IPI00261627 | 50 kDa | 11 | 7 | 6 | 6 |
| Mag Isoform L-MAG of Myelin-associated glycoprotein precursor | IPI00276515 | 69 kDa | 9 | 7 | 6 | 6 |
| Timm50 Import inner membrane translocase subunit TIM50, mitochondrial precursor | IPI00111045 | 40 kDa | 8 | 7 | 6 | 6 |
| Rab11b Ras-related protein Rab-11B | IPI00135869 | 24 kDa | 6 | 6 | 6 | 6 |
| Ruvbl2 RuvB-like 2 | IPI00123557 | 51 kDa | 6 | 5 | 6 | 6 |
| Scye1 small inducible cytokine subfamily E, member 1 | IPI00132194 | 35 kDa | 6 | 5 | 6 | 6 |
| Stip1 Stress-induced-phosphoprotein 1 | IPI00121514 | 63 kDa | 6 | 5 | 6 | 6 |
| Ppia Peptidyl-prolyl cis-trans isomerase | IPI00554989 | 18 kDa | 6 | 5 | 6 | 6 |
| Ilf2 Interleukin enhancer-binding factor 2 | IPI00318550 (+1) | 43 kDa | 5 | 5 | 6 | 6 |
| Psmc1 26S protease regulatory subunit 4 | IPI00133428 | 49 kDa | 4 | 4 | 6 | 6 |
| Copb2 Coatomer subunit beta' | IPI00115097 | 102 kDa | 3 | 3 | 6 | 6 |
| Cttn Src substrate cortactin | IPI00118143 (+2) | 61 kDa | 3 | 3 | 6 | 6 |
| Atp1b2 Sodium/potassium-transporting ATPase subunit beta-2 | IPI00123704 | 33 kDa | 2 | 2 | 6 | 6 |

| | | | | | | |
|---|---|---|---|---|---|---|
| Flna Isoform 1 of Filamin-A | IPI00131138 (+4) | 281 kDa | 2 | 2 | 6 | 6 |
| Dpysl5 Dihydropyrimidinase-related protein 5 | IPI00624192 | 62 kDa | 0 | 0 | 6 | 6 |
| Eif3d Eukaryotic translation initiation factor 3 subunit D | IPI00762774 (+1) | 64 kDa | 0 | 0 | 6 | 6 |
| Inpp4a Isoform 1 of Type I inositol-3,4-bisphosphate 4-phosphatase | IPI00110426 | 106 kDa | 0 | 0 | 6 | 6 |
| Nrbp1 NOD-derived CD11c +ve dendritic cells cDNA, RIKEN full-length enriched library, clon | IPI00664246 | 61 kDa | 0 | 0 | 6 | 6 |
| Prps1 Ribose-phosphate pyrophosphokinase | IPI00318204 (+4) | 35 kDa | 0 | 0 | 6 | 6 |
| Mkrn2 Makorin-2 | IPI00620251 (+1) | 47 kDa | 0 | 0 | 6 | 6 |
| Ndufa8 NADH dehydrogenase [ubiquinone] 1 alpha subcomplex subunit 8 | IPI00120984 | 20 kDa | 0 | 0 | 6 | 6 |
| Gcs1 Mannosyl-oligosaccharide glucosidase | IPI00330323 | 92 kDa | 0 | 0 | 6 | 6 |
| Ikbkap Elongator complex protein 1 | IPI00380780 | 150 kDa | 0 | 0 | 6 | 6 |
| Ap3m2 AP-3 complex subunit mu-2 | IPI00124286 | 47 kDa | 0 | 0 | 6 | 6 |
| Tubb5 Tubulin beta-5 chain | IPI00117352 | 50 kDa | 725 | 5 | 394 | 5 |
| Ank3 ankyrin 3, epithelial isoform a | IPI00173248 (+5) | 188 kDa | 0 | 0 | 41 | 5 |
| Plp1 Isoform 1 of Myelin proteolipid protein | IPI00263013 | 30 kDa | 19 | 5 | 31 | 5 |
| Gnb2 Guanine nucleotide-binding protein G(I)/G(S)/G(T) subunit beta-2 | IPI00162780 | 37 kDa | 15 | 4 | 22 | 5 |
| Ank1 Isoform Br2 of Ankyrin-1 | IPI00119871 (+8) | 203 kDa | 21 | 3 | 20 | 5 |
| Elavl4 Elavl4 protein | IPI00466120 (+2) | 45 kDa | 8 | 2 | 19 | 5 |
| sp_HBA_HUMAN | IPIsp_HBA_HUMAN | 15 kDa | 17 | 5 | 14 | 5 |
| - 11 kDa protein | IPI00329998 (+9) | 11 kDa | 10 | 5 | 14 | 5 |
| Hsd17b10 Hydroxysteroid (17-beta) dehydrogenase 10 | IPI00626132 (+1) | 28 kDa | 14 | 9 | 11 | 5 |
| Dmxl1 Dmx-like 1 | IPI00674554 | 337 kDa | 0 | 0 | 11 | 5 |
| Vim Vimentin | IPI00227299 | 54 kDa | 17 | 8 | 10 | 5 |
| Klc1 kinesin light chain 1 isoform 1D | IPI00403890 (+3) | 69 kDa | 7 | 4 | 10 | 5 |
| Actr10 Actin-related protein 10 | IPI00137206 | 46 kDa | 2 | 2 | 10 | 5 |
| Psma4 Proteasome subunit alpha type-4 | IPI00277001 | 29 kDa | 9 | 5 | 8 | 5 |
| Sept4 Isoform A of Septin-4 | IPI00123219 (+2) | 55 kDa | 7 | 3 | 8 | 5 |
| Chl1 cell adhesion molecule with homology to L1CAM | IPI00831546 (+1) | 135 kDa | 0 | 0 | 8 | 5 |
| LOC100048210 similar to ribosomal protein L3 isoform 1 | IPI00605077 (+5) | 46 kDa | 0 | 0 | 8 | 5 |
| Stk24 Serine/threonine-protein kinase 24 | IPI00116065 | 48 kDa | 8 | 8 | 7 | 5 |
| Psma6 Proteasome subunit alpha type-6 | IPI00131845 | 27 kDa | 11 | 7 | 7 | 5 |
| Vapb Vesicle-associated membrane protein-associated protein B | IPI00135655 (+1) | 27 kDa | 11 | 7 | 7 | 5 |
| Atp1b1 Sodium/potassium-transporting ATPase subunit beta-1 | IPI00121550 | 35 kDa | 8 | 5 | 7 | 5 |
| Napb Beta-soluble NSF attachment protein | IPI00311515 | 34 kDa | 4 | 4 | 7 | 5 |
| ENSMUSG00000060419 Rps16 protein | IPI00469918 (+1) | 19 kDa | 4 | 4 | 7 | 5 |
| Dbt Lipoamide acyltransferase component of branched-chain alpha-keto acid dehydrogenas | IPI00130535 (+1) | 53 kDa | 4 | 3 | 7 | 5 |
| Sparcl1 SPARC-like protein 1 precursor | IPI00308484 | 72 kDa | 3 | 2 | 7 | 5 |
| Rgs7 Rgs7 protein | IPI00118769 | 55 kDa | 3 | 2 | 7 | 5 |
| Eif3i Eukaryotic translation initiation factor 3 subunit I | IPI00269613 | 36 kDa | 0 | 0 | 7 | 5 |
| Rps3a 40S ribosomal protein S3a | IPI00331345 (+2) | 30 kDa | 0 | 0 | 7 | 5 |
| Ddx6 Probable ATP-dependent RNA helicase DDX6 | IPI00109932 | 54 kDa | 0 | 0 | 7 | 5 |
| Tom1l2 Isoform 1 of TOM1-like protein 2 | IPI00322033 | 56 kDa | 144 | 29 | 6 | 5 |

| | | | | | | |
|---|---|---|---|---|---|---|
| Tpi1 Triosephosphate isomerase | IPI00467833 | 27 kDa | 12 | 7 | 6 | 5 |
| Nudt21 Cleavage and polyadenylation specificity factor subunit 5 | IPI00132473 | 26 kDa | 9 | 7 | 6 | 5 |
| Rps18 40S ribosomal protein S18 | IPI00317590 (+3) | 18 kDa | 6 | 5 | 6 | 5 |
| Syp Syp protein | IPI00756139 | 33 kDa | 6 | 3 | 6 | 5 |
| Arpc2 Actin-related protein 2/3 complex subunit 2 | IPI00661414 | 34 kDa | 3 | 3 | 6 | 5 |
| Cops6 COP9 signalosome complex subunit 6 | IPI00131873 (+1) | 36 kDa | 3 | 3 | 6 | 5 |
| Phactr1 Isoform 1 of Phosphatase and actin regulator 1 | IPI00719908 (+1) | 66 kDa | 3 | 3 | 6 | 5 |
| Pygb Glycogen phosphorylase, brain form | IPI00229796 | 97 kDa | 3 | 3 | 6 | 5 |
| Pspc1 Isoform 1 of Paraspeckle component 1 | IPI00321266 (+1) | 59 kDa | 3 | 3 | 6 | 5 |
| Prdx3 Thioredoxin-dependent peroxide reductase, mitochondrial precursor | IPI00116192 | 28 kDa | 3 | 2 | 6 | 5 |
| Endogl1 Nuclease EXOG, mitochondrial precursor | IPI00224069 | 41 kDa | 2 | 2 | 6 | 5 |
| Spag9 Isoform 2 of C-jun-amino-terminal kinase-interacting protein 4 | IPI00462746 (+2) | 145 kDa | 0 | 0 | 6 | 5 |
| Hexb Beta-hexosaminidase subunit beta precursor | IPI00115530 | 61 kDa | 0 | 0 | 6 | 5 |
| - 28 kDa protein | IPI00469509 | 28 kDa | 0 | 0 | 6 | 5 |
| Nt5dc3;BC030307 5'-nucleotidase domain-containing protein 3 | IPI00648312 | 63 kDa | 0 | 0 | 6 | 5 |
| Strap Serine-threonine kinase receptor-associated protein | IPI00130670 | 39 kDa | 0 | 0 | 6 | 5 |
| Kif2a Isoform 2 of Kinesin-like protein KIF2A | IPI00867847 | 75 kDa | 19 | 15 | 5 | 5 |
| 6330407J23Rik Uncharacterized protein C6orf174 homolog precursor | IPI00403618 | 103 kDa | 15 | 9 | 5 | 5 |
| Nipsnap1 Protein NipSnap homolog 1 | IPI00115824 | 33 kDa | 16 | 8 | 5 | 5 |
| Pcmt1 Isoform 1 of Protein-L-isoaspartate(D-aspartate) O-methyltransferase | IPI00329913 (+2) | 25 kDa | 7 | 5 | 5 | 5 |
| 2900073G15Rik myosin light chain, regulatory B-like | IPI00109044 (+1) | 20 kDa | 7 | 5 | 5 | 5 |
| Arf1 ADP-ribosylation factor 1 | IPI00221613 (+1) | 21 kDa | 7 | 5 | 5 | 5 |
| Suclg1 Succinyl-CoA ligase [GDP-forming] subunit alpha, mitochondrial precursor | IPI00406442 | 35 kDa | 6 | 5 | 5 | 5 |
| Gnaq cDNA, RIKEN full-length enriched library, clone:M5C1028J20 product:guanine nucleoti | IPI00228618 | 42 kDa | 4 | 4 | 5 | 5 |
| Dpp6 Dipeptidyl aminopeptidase-like protein 6 | IPI00130389 (+3) | 91 kDa | 3 | 3 | 5 | 5 |
| Eif3b Eif3b protein | IPI00229859 (+1) | 109 kDa | 3 | 3 | 5 | 5 |
| Srr Isoform 1 of Serine racemase | IPI00138217 (+1) | 36 kDa | 3 | 2 | 5 | 5 |
| Dlst Isoform 1 of Dihydrolipoyllysine-residue succinyltransferase component of 2- oxoglutar | IPI00134809 | 49 kDa | 2 | 2 | 5 | 5 |
| Ogt Isoform 1 of UDP-N-acetylglucosamine--peptide N-acetylglucosaminyltransferase 110 kI | IPI00420870 (+1) | 117 kDa | 2 | 2 | 5 | 5 |
| Got1 glutamate oxaloacetate transaminase 1, soluble | IPI00877205 | 46 kDa | 2 | 2 | 5 | 5 |
| Rars Arginyl-tRNA synthetase, cytoplasmic | IPI00315488 | 76 kDa | 0 | 0 | 5 | 5 |
| Ptpn23 Isoform 1 of Tyrosine-protein phosphatase non-receptor type 23 | IPI00464166 (+1) | 185 kDa | 0 | 0 | 5 | 5 |
| Pgam5 Isoform 2 of Phosphoglycerate mutase family member 5 precursor | IPI00226387 (+1) | 32 kDa | 0 | 0 | 5 | 5 |
| U2af1 Splicing factor U2AF 35 kDa subunit | IPI00318548 (+2) | 28 kDa | 0 | 0 | 5 | 5 |
| Srp72 signal recognition particle 72 | IPI00316133 (+1) | 75 kDa | 0 | 0 | 5 | 5 |
| Eml1 Isoform 2 of Echinoderm microtubule-associated protein-like 1 | IPI00354624 (+4) | 86 kDa | 0 | 0 | 5 | 5 |
| - 29 kDa protein | IPI00108454 (+2) | 29 kDa | 0 | 0 | 5 | 5 |
| Vps26b Isoform 1 of Vacuolar protein sorting-associated protein 26B | IPI00223759 (+1) | 39 kDa | 0 | 0 | 5 | 5 |
| Ddx19a ATP-dependent RNA helicase DDX19A | IPI00123651 (+3) | 54 kDa | 0 | 0 | 5 | 5 |
| Lgi3 Leucine-rich repeat LGI family member 3 precursor | IPI00170121 | 62 kDa | 0 | 0 | 5 | 5 |
| Ankfy1 Isoform 1 of Ankyrin repeat and FYVE domain-containing protein 1 | IPI00329843 (+1) | 129 kDa | 0 | 0 | 5 | 5 |

| | | | | | | |
|---|---|---|---|---|---|---|
| Ganab Isoform 2 of Neutral alpha-glucosidase AB precursor | IPI00115679 (+1) | 109 kDa | 0 | 0 | 5 | 5 |
| Hectd1 similar to HECT domain containing 1 | IPI00652600 (+1) | 289 kDa | 0 | 0 | 5 | 5 |
| Lancl1 LanC-like protein 1 | IPI00131535 (+1) | 45 kDa | 0 | 0 | 5 | 5 |
| Copg2 Isoform 1 of Coatomer subunit gamma-2 | IPI00316359 (+1) | 98 kDa | 0 | 0 | 5 | 5 |
| Anapc1 Anaphase-promoting complex subunit 1 | IPI00137228 | 216 kDa | 0 | 0 | 5 | 5 |
| A1bg alpha-1-B glycoprotein | IPI00129965 | 57 kDa | 0 | 0 | 5 | 5 |
| Vps52 Isoform 1 of Vacuolar protein sorting-associated protein 52 homolog | IPI00623259 | 82 kDa | 0 | 0 | 5 | 5 |
| Tuba4a Tubulin alpha-4A chain | IPI00117350 | 50 kDa | 305 | 5 | 248 | 4 |
| sp_TRYP_PIG | IPIsp_TRYP_PIG | 24 kDa | 148 | 4 | 115 | 4 |
| Ncam1 neural cell adhesion molecule 1 isoform 1 | IPI00831465 | 93 kDa | 94 | 2 | 113 | 4 |
| Acta1 Actin, alpha skeletal muscle | IPI00110827 (+2) | 42 kDa | 33 | 4 | 53 | 4 |
| Ank2 ankyrin 2, brain isoform 3 | IPI00404055 | 103 kDa | 17 | 8 | 19 | 4 |
| Actr1b Beta-centractin | IPI00153990 | 42 kDa | 9 | 3 | 18 | 4 |
| Nrxn3 neurexin III | IPI00228680 | 175 kDa | 0 | 0 | 12 | 4 |
| Ewsr1 Ewing sarcoma breakpoint region 1 | IPI00322492 (+3) | 69 kDa | 16 | 6 | 11 | 4 |
| Marcks Myristoylated alanine-rich C-kinase substrate | IPI00229534 | 30 kDa | 7 | 3 | 10 | 4 |
| sp_K1HB_HUMAN | IPIsp_K1HB_HUMAN | 46 kDa | 11 | 3 | 9 | 4 |
| Prkar2a CAMP-dependent protein kinase type II-alpha regulatory chain | IPI00169788 | 46 kDa | 0 | 0 | 9 | 4 |
| Thoc4 Isoform 1 of THO complex subunit 4 | IPI00114407 | 27 kDa | 21 | 7 | 8 | 4 |
| Dynll2 Dynein light chain 2, cytoplasmic | IPI00132734 | 10 kDa | 9 | 5 | 7 | 4 |
| Eno2 Gamma-enolase | IPI00331704 | 47 kDa | 5 | 4 | 7 | 4 |
| Mtap7d1 Isoform 1 of MAP7 domain-containing protein 1 | IPI00282957 (+1) | 93 kDa | 4 | 4 | 7 | 4 |
| Ruvbl1 RuvB-like 1 | IPI00133985 (+1) | 50 kDa | 4 | 3 | 7 | 4 |
| Psmc5 26S protease regulatory subunit 8 | IPI00135640 | 46 kDa | 3 | 3 | 7 | 4 |
| EG268795 hypothetical protein isoform 2 | IPI00265107 (+4) | 30 kDa | 0 | 0 | 7 | 4 |
| Hagh Hydroxyacylglutathione hydrolase | IPI00115866 | 34 kDa | 0 | 0 | 7 | 4 |
| Capza1 F-actin-capping protein subunit alpha-1 | IPI00330063 (+3) | 33 kDa | 0 | 0 | 7 | 4 |
| Gap43 Neuromodulin | IPI00128973 | 24 kDa | 10 | 8 | 6 | 4 |
| C1qbp complement component 1, q subcomponent binding protein | IPI00132799 | 31 kDa | 21 | 7 | 6 | 4 |
| 2010305A19Rik Hcp beta-lactamase-like protein C1orf163 homolog | IPI00121834 | 26 kDa | 7 | 6 | 6 | 4 |
| Pgam1 Phosphoglycerate mutase 1 | IPI00457898 | 29 kDa | 9 | 4 | 6 | 4 |
| Eif4g3 Isoform 4 of Eukaryotic translation initiation factor 4 gamma 3 | IPI00417153 (+4) | 175 kDa | 6 | 3 | 6 | 4 |
| Nono Isoform 1 of Non-POU domain-containing octamer-binding protein | IPI00320016 (+1) | 55 kDa | 5 | 2 | 6 | 4 |
| Pcbp2 Isoform 1 of Poly(rC)-binding protein 2 | IPI00127707 (+2) | 38 kDa | 4 | 2 | 6 | 4 |
| Snrpd1 Small nuclear ribonucleoprotein Sm D1 | IPI00322749 | 13 kDa | 2 | 2 | 6 | 4 |
| C2cd2l Transmembrane protein 24 | IPI00119785 | 76 kDa | 2 | 2 | 6 | 4 |
| Add3 Adducin 3 | IPI00387580 (+1) | 79 kDa | 0 | 0 | 6 | 4 |
| Mrps15 28S ribosomal protein S15, mitochondrial precursor | IPI00321858 | 29 kDa | 0 | 0 | 6 | 4 |
| Pcbd1 Pterin-4-alpha-carbinolamine dehydratase | IPI00223800 | 12 kDa | 0 | 0 | 6 | 4 |
| Pdxk Pyridoxal kinase | IPI00283511 | 35 kDa | 0 | 0 | 6 | 4 |
| Coro1a Coronin-1A | IPI00323600 (+1) | 51 kDa | 0 | 0 | 6 | 4 |

| | | | | | | |
|---|---|---|---|---|---|---|
| Ppfia3 Liprin-alpha-3 | IPI00399905 | 116 kDa | 28 | 21 | 5 | 4 |
| P140 131 kDa protein | IPI00875486 (+2) | 131 kDa | 20 | 18 | 5 | 4 |
| Mobkl3 Mps one binder kinase activator-like 3 | IPI00420832 | 26 kDa | 35 | 14 | 5 | 4 |
| Eea1 Early endosome antigen 1 | IPI00453776 | 161 kDa | 13 | 13 | 5 | 4 |
| Cltb Isoform 1 of Clathrin light chain B | IPI00228978 | 25 kDa | 32 | 11 | 5 | 4 |
| Myl6 Isoform Smooth muscle of Myosin light polypeptide 6 | IPI00354819 (+1) | 17 kDa | 12 | 9 | 5 | 4 |
| Lin7a Lin-7 homolog A | IPI00344142 | 26 kDa | 15 | 8 | 5 | 4 |
| Cops3 COP9 signalosome complex subunit 3 | IPI00131870 | 48 kDa | 8 | 8 | 5 | 4 |
| Atp6v1g2 Vacuolar proton pump subunit G 2 | IPI00123817 | 14 kDa | 7 | 7 | 5 | 4 |
| Prdx5 Isoform Mitochondrial of Peroxiredoxin-5, mitochondrial precursor | IPI00129517 (+1) | 22 kDa | 9 | 6 | 5 | 4 |
| Hgs HGF-regulated tyrosine kinase substrate | IPI00331568 (+1) | 86 kDa | 7 | 6 | 5 | 4 |
| Psmb2 Proteasome subunit beta type-2 | IPI00128945 | 23 kDa | 7 | 6 | 5 | 4 |
| Vapa Vesicle-associated membrane protein-associated protein A | IPI00125267 | 28 kDa | 13 | 5 | 5 | 4 |
| Gbas Protein NipSnap homolog 2 | IPI00115827 (+1) | 33 kDa | 12 | 5 | 5 | 4 |
| Prdx2 Peroxiredoxin-2 | IPI00117910 | 22 kDa | 5 | 4 | 5 | 4 |
| Cep170 Isoform 1 of Centrosomal protein of 170 kDa | IPI00667973 | 175 kDa | 4 | 4 | 5 | 4 |
| Mrps34 Mitochondrial 28S ribosomal protein S34 | IPI00120709 | 26 kDa | 4 | 3 | 5 | 4 |
| Pcsk1n ProSAAS precursor | IPI00135604 | 27 kDa | 4 | 3 | 5 | 4 |
| Cdc16 Cell division cycle protein 16 homolog | IPI00221621 | 71 kDa | 4 | 3 | 5 | 4 |
| Dpysl4 Isoform 1 of Dihydropyrimidinase-related protein 4 | IPI00313151 (+2) | 62 kDa | 3 | 3 | 5 | 4 |
| LOC100045866;Tceb1 Transcription elongation factor B polypeptide 1 | IPI00323130 | 12 kDa | 2 | 2 | 5 | 4 |
| 1500001M20Rik Isoform 1 of MMP37-like protein, mitochondrial precursor | IPI00108431 | 38 kDa | 2 | 2 | 5 | 4 |
| Rap1gds1 RAP1, GTP-GDP dissociation stimulator 1 isoform b | IPI00310972 (+1) | 61 kDa | 2 | 2 | 5 | 4 |
| Tanc2 Isoform 2 of Protein TANC2 | IPI00551112 (+1) | 221 kDa | 0 | 0 | 5 | 4 |
| EG626175 similar to ribosomal protein S26 | IPI00261455 (+3) | 13 kDa | 0 | 0 | 5 | 4 |
| Iars Isoleucyl-tRNA synthetase, cytoplasmic | IPI00225201 | 144 kDa | 0 | 0 | 5 | 4 |
| Pzp Alpha-2-macroglobulin precursor | IPI00624663 | 167 kDa | 0 | 0 | 5 | 4 |
| Pacs1 Phosphofurin acidic cluster sorting protein 1 | IPI00321922 | 105 kDa | 0 | 0 | 5 | 4 |
| Enpp6 Ectonucleotide pyrophosphatase/phosphodiesterase family member 6 precursor | IPI00308077 (+1) | 51 kDa | 0 | 0 | 5 | 4 |
| Nrbp2 Nuclear receptor-binding protein 2 | IPI00263549 | 30 kDa | 0 | 0 | 5 | 4 |
| Mrps9 28S ribosomal protein S9, mitochondrial precursor | IPI00336292 (+1) | 45 kDa | 0 | 0 | 5 | 4 |
| Centg1 Isoform 1 of Centaurin-gamma-1 | IPI00349020 (+1) | 125 kDa | 0 | 0 | 5 | 4 |
| Msi2 Isoform 1 of RNA-binding protein Musashi homolog 2 | IPI00120924 (+2) | 37 kDa | 0 | 0 | 5 | 4 |
| Gipc1 PDZ domain-containing protein GIPC1 | IPI00129107 | 36 kDa | 69 | 25 | 4 | 4 |
| Klhl22 Isoform 2 of Kelch-like protein 22 | IPI00224738 (+1) | 74 kDa | 42 | 25 | 4 | 4 |
| 2310035C23Rik Isoform 1 of LisH domain and HEAT repeat-containing protein KIAA1468 | IPI00228693 | 135 kDa | 8 | 8 | 4 | 4 |
| Pdcd10 Programmed cell death protein 10 | IPI00331188 | 25 kDa | 14 | 7 | 4 | 4 |
| Gstm1 Glutathione S-transferase Mu 1 | IPI00230212 (+1) | 26 kDa | 8 | 7 | 4 | 4 |
| Rufy3 Isoform 3 of Protein RUFY3 | IPI00453965 (+3) | 55 kDa | 9 | 6 | 4 | 4 |
| Psmb4 Proteasome subunit beta type-4 precursor | IPI00129512 | 29 kDa | 8 | 5 | 4 | 4 |
| Psma2 Proteasome subunit alpha type | IPI00420745 (+1) | 28 kDa | 6 | 5 | 4 | 4 |

| | | | | | | |
|---|---|---|---|---|---|---|
| LOC100047429;Atp5o ATP synthase subunit O, mitochondrial precursor | IPI00118986 (+1) | 23 kDa | 6 | 5 | 4 | 4 |
| Sdhb Succinate dehydrogenase [ubiquinone] iron-sulfur subunit, mitochondrial precursor | IPI00338536 | 32 kDa | 5 | 5 | 4 | 4 |
| Ubap2l Isoform 5 of Ubiquitin-associated protein 2-like | IPI00407835 (+1) | 117 kDa | 5 | 5 | 4 | 4 |
| Psmb7 Proteasome subunit beta type-7 precursor | IPI00136483 | 30 kDa | 7 | 4 | 4 | 4 |
| Mapt 76 kDa protein | IPI00113584 (+6) | 76 kDa | 6 | 4 | 4 | 4 |
| Cryab Alpha-crystallin B chain | IPI00138274 | 20 kDa | 6 | 4 | 4 | 4 |
| Ppp2r5e Serine/threonine-protein phosphatase 2A 56 kDa regulatory subunit epsilon isofor | IPI00224697 (+1) | 55 kDa | 5 | 4 | 4 | 4 |
| EG625298;Rps13;LOC100044992;ENSMUSG00000066362;LOC100048340 40S ribosomal pro | IPI00125901 | 17 kDa | 4 | 4 | 4 | 4 |
| Gdap1 Isoform 1 of Ganglioside-induced differentiation-associated protein 1 | IPI00134137 | 41 kDa | 4 | 4 | 4 | 4 |
| Dync1i2 Cytoplasmic dynein 1 intermediate chain 2 | IPI00131086 (+4) | 68 kDa | 4 | 4 | 4 | 4 |
| Mtap7 Isoform 2 of Ensconsin | IPI00380510 | 84 kDa | 4 | 3 | 4 | 4 |
| Cyc1 Isoform 1 of Cytochrome c1, heme protein, mitochondrial precursor | IPI00132728 (+1) | 35 kDa | 4 | 3 | 4 | 4 |
| Rogdi Isoform 1 of Protein rogdi homolog | IPI00399812 (+1) | 32 kDa | 3 | 3 | 4 | 4 |
| Phyhip Phytanoyl-CoA hydroxylase-interacting protein | IPI00169622 | 38 kDa | 3 | 3 | 4 | 4 |
| Snrpd2 Small nuclear ribonucleoprotein Sm D2 | IPI00119220 | 14 kDa | 3 | 2 | 4 | 4 |
| Rabgap1 Isoform 1 of Rab GTPase-activating protein 1 | IPI00378156 | 121 kDa | 2 | 2 | 4 | 4 |
| Abi2 Abl interactor 2 | IPI00380956 (+1) | 49 kDa | 2 | 2 | 4 | 4 |
| Ppm1e Protein phosphatase 1E | IPI00461286 | 83 kDa | 2 | 2 | 4 | 4 |
| Dync1li1 Cytoplasmic dynein 1 light intermediate chain 1 | IPI00153421 | 57 kDa | 2 | 2 | 4 | 4 |
| Sqstm1 Isoform 1 of Sequestosome-1 | IPI00133374 (+1) | 48 kDa | 2 | 2 | 4 | 4 |
| Snrpb Small nuclear ribonucleoprotein-associated protein B | IPI00114052 (+1) | 24 kDa | 2 | 2 | 4 | 4 |
| Pfkm 6-phosphofructokinase, muscle type | IPI00331541 | 85 kDa | 0 | 0 | 4 | 4 |
| Snrpa U1 small nuclear ribonucleoprotein A | IPI00122350 (+1) | 32 kDa | 0 | 0 | 4 | 4 |
| Snrpa1 U2 small nuclear ribonucleoprotein A' | IPI00170008 | 28 kDa | 0 | 0 | 4 | 4 |
| Rps19 40S ribosomal protein S19 | IPI00113241 (+7) | 16 kDa | 0 | 0 | 4 | 4 |
| Htra2 Serine protease HTRA2, mitochondrial precursor | IPI00275992 (+2) | 49 kDa | 0 | 0 | 4 | 4 |
| Sh3glb2 SH3-domain GRB2-like endophilin B2 | IPI00756786 | 45 kDa | 0 | 0 | 4 | 4 |
| Nptx1 Neuronal pentraxin-1 precursor | IPI00125992 | 47 kDa | 0 | 0 | 4 | 4 |
| Mrps31 28S ribosomal protein S31, mitochondrial precursor | IPI00124828 | 44 kDa | 0 | 0 | 4 | 4 |
| Tnks1bp1 182 kDa tankyrase 1-binding protein | IPI00459443 | 182 kDa | 0 | 0 | 4 | 4 |
| Smarcc2 Isoform 2 of SWI/SNF-related matrix-associated actin-dependent regulator of chro | IPI00381019 (+2) | 121 kDa | 0 | 0 | 4 | 4 |
| Rpl6 60S ribosomal protein L6 | IPI00313222 (+2) | 34 kDa | 0 | 0 | 4 | 4 |
| Asah1 Acid ceramidase precursor | IPI00125266 | 45 kDa | 0 | 0 | 4 | 4 |
| Rpl7 60S ribosomal protein L7 | IPI00311236 | 31 kDa | 0 | 0 | 4 | 4 |
| Aldh9a1 aldehyde dehydrogenase 9, subfamily A1 | IPI00124372 | 56 kDa | 0 | 0 | 4 | 4 |
| Dap3 Isoform 1 of Mitochondrial 28S ribosomal protein S29 | IPI00275050 (+1) | 45 kDa | 0 | 0 | 4 | 4 |
| Mrps35 28S ribosomal protein S35, mitochondrial precursor | IPI00222538 (+1) | 36 kDa | 0 | 0 | 4 | 4 |
| Kctd13 BTB/POZ domain-containing protein KCTD13 | IPI00228759 | 36 kDa | 0 | 0 | 4 | 4 |
| Gucy1b3 Guanylate cyclase soluble subunit beta-1 | IPI00118880 (+1) | 71 kDa | 0 | 0 | 4 | 4 |
| Kcnj10 ATP-sensitive inward rectifier potassium channel 10 | IPI00124792 (+1) | 42 kDa | 0 | 0 | 4 | 4 |
| Amz2 archaemetzincins-2 | IPI00625349 | 42 kDa | 0 | 0 | 4 | 4 |

| | | | | | | |
|---|---|---|---|---|---|---|
| 2010300C02Rik 2010300C02Rik protein | IPI00454053 (+1) | 126 kDa | 0 | 0 | 4 | 4 |
| Sccpdh Probable saccharopine dehydrogenase | IPI00153266 (+1) | 47 kDa | 0 | 0 | 4 | 4 |
| Plcb1 Plcb1 protein | IPI00468121 (+1) | 138 kDa | 0 | 0 | 4 | 4 |
| Fhl2 Four and a half LIM domains protein 2 | IPI00118205 | 32 kDa | 0 | 0 | 4 | 4 |
| Ivd Isovaleryl-CoA dehydrogenase, mitochondrial precursor | IPI00471246 | 46 kDa | 0 | 0 | 4 | 4 |
| Ampd2 Adenosine monophosphate deaminase 2 | IPI00623834 | 95 kDa | 0 | 0 | 4 | 4 |
| Cdc42bpb Serine/threonine-protein kinase MRCK beta | IPI00114613 | 195 kDa | 0 | 0 | 4 | 4 |
| Armc9 Isoform 1 of LisH domain-containing protein ARMC9 | IPI00378172 (+4) | 92 kDa | 0 | 0 | 4 | 4 |
| Elfn2 Leucine-rich repeat and fibronectin type-III domain-containing protein 6 precursor | IPI00471006 | 90 kDa | 0 | 0 | 4 | 4 |
| Araf A-Raf proto-oncogene serine/threonine-protein kinase | IPI00320608 (+3) | 68 kDa | 0 | 0 | 4 | 4 |
| Nos1 Isoform N-NOS-1 of Nitric oxide synthase, brain | IPI00129191 (+4) | 160 kDa | 0 | 0 | 4 | 4 |
| Mrps5 Mitochondrial 28S ribosomal protein S5 | IPI00118196 | 48 kDa | 0 | 0 | 4 | 4 |
| A230046K03Rik Uncharacterized protein KIAA1033 | IPI00831049 | 136 kDa | 0 | 0 | 4 | 4 |
| Cyfip1 Isoform 1 of Cytoplasmic FMR1-interacting protein 1 | IPI00330476 | 145 kDa | 19 | 2 | 26 | 3 |
| Atp2b2 Plasma membrane calcium-transporting ATPase 2 | IPI00127713 (+1) | 133 kDa | 0 | 0 | 25 | 3 |
| Camk2d Isoform 1 of Calcium/calmodulin-dependent protein kinase type II delta chain | IPI00112584 (+4) | 56 kDa | 0 | 0 | 21 | 3 |
| Hnrnph2 Heterogeneous nuclear ribonucleoprotein H2 | IPI00108143 | 49 kDa | 0 | 0 | 17 | 3 |
| Hba-a1 13 days embryo liver cDNA, RIKEN full-length enriched library, clone:2510040B16 pr | IPI00110658 (+1) | 15 kDa | 13 | 3 | 15 | 3 |
| Kif5b Kinesin-1 heavy chain | IPI00124954 (+1) | 110 kDa | 10 | 3 | 10 | 3 |
| Rtn3 Isoform 1 of Reticulon-3 | IPI00470981 (+1) | 104 kDa | 25 | 10 | 9 | 3 |
| Basp1 Brain acid soluble protein 1 | IPI00129519 | 22 kDa | 4 | 2 | 9 | 3 |
| Csnk2a2 Casein kinase II subunit alpha' | IPI00118795 | 41 kDa | 0 | 0 | 8 | 3 |
| Rab14 Ras-related protein Rab-14 | IPI00126042 | 24 kDa | 0 | 0 | 8 | 3 |
| Gpm6a Neuronal membrane glycoprotein M6-a | IPI00122974 | 31 kDa | 6 | 3 | 7 | 3 |
| Ldha L-lactate dehydrogenase A chain | IPI00319994 (+1) | 36 kDa | 0 | 0 | 7 | 3 |
| Prrt2 proline-rich transmembrane protein 2 | IPI00281761 | 36 kDa | 8 | 4 | 6 | 3 |
| Aplp2 CDE1-binding protein CDEBP | IPI00121338 (+1) | 87 kDa | 2 | 2 | 6 | 3 |
| Sf3a1 Splicing factor 3 subunit 1 | IPI00408796 | 89 kDa | 0 | 0 | 6 | 3 |
| Pnkd Isoform 4 of Probable hydrolase PNKD | IPI00121559 (+2) | 41 kDa | 0 | 0 | 6 | 3 |
| App Isoform APP770 of Amyloid beta A4 protein precursor (Fragment) | IPI00114389 (+2) | 87 kDa | 5 | 4 | 5 | 3 |
| Sec31a SEC31-like 1 | IPI00377592 (+1) | 144 kDa | 4 | 4 | 5 | 3 |
| Rap1gap Rap1 GTPase-activating protein | IPI00459745 (+3) | 81 kDa | 3 | 3 | 5 | 3 |
| Calm3;Calm2;Calm1 Calmodulin | IPI00761696 | 17 kDa | 4 | 2 | 5 | 3 |
| Rps17 40S ribosomal protein S17 | IPI00465880 | 16 kDa | 3 | 2 | 5 | 3 |
| 2410166I05Rik Isoform 1 of Protein FAM54B | IPI00109033 | 32 kDa | 3 | 2 | 5 | 3 |
| Pip5k1c Isoform 3 of Phosphatidylinositol-4-phosphate 5-kinase type-1 gamma | IPI00403328 (+2) | 70 kDa | 2 | 2 | 5 | 3 |
| Rps14 40S ribosomal protein S14 | IPI00322562 (+1) | 16 kDa | 2 | 2 | 5 | 3 |
| LOC672959;Rps12 40S ribosomal protein S12 | IPI00225634 (+1) | 15 kDa | 2 | 2 | 5 | 3 |
| Tsc22d1 TSC22-related inducible leucine zipper 1b | IPI00420803 (+1) | 110 kDa | 0 | 0 | 5 | 3 |
| Neo1 Isoform 1 of Neogenin precursor | IPI00129159 (+3) | 163 kDa | 0 | 0 | 5 | 3 |
| Rpl18a 60S ribosomal protein L18a | IPI00162790 (+1) | 21 kDa | 0 | 0 | 5 | 3 |

| | | | | | | |
|---|---|---|---|---|---|---|
| Pafah1b1 Isoform 1 of Platelet-activating factor acetylhydrolase IB subunit alpha | IPI00309207 (+1) | 47 kDa | 0 | 0 | 5 | 3 |
| Rasgrf2 Ras-specific guanine nucleotide-releasing factor 2 | IPI00108372 | 136 kDa | 0 | 0 | 5 | 3 |
| Mical3 Adult male testis cDNA, RIKEN full-length enriched library, clone:4921518D21 produc | IPI00229906 (+3) | 99 kDa | 0 | 0 | 5 | 3 |
| Purg Isoform 1 of Purine-rich element-binding protein gamma | IPI00153880 | 40 kDa | 0 | 0 | 5 | 3 |
| Strn4 Isoform 1 of Striatin-4 | IPI00119524 (+1) | 82 kDa | 30 | 15 | 4 | 3 |
| 1110031B06Rik Isoform 1 of t-SNARE coiled-coil homology domain-containing protein C1orf | IPI00269408 | 47 kDa | 7 | 7 | 4 | 3 |
| Psmc4 26S protease regulatory subunit 6B | IPI00108895 (+1) | 47 kDa | 9 | 6 | 4 | 3 |
| Gripap1 GRIP1-associated protein 1 | IPI00267855 (+2) | 93 kDa | 7 | 6 | 4 | 3 |
| Trim28 Isoform 1 of Transcription intermediary factor 1-beta | IPI00312128 | 89 kDa | 7 | 5 | 4 | 3 |
| Rps20 40S ribosomal protein S20 | IPI00323819 | 13 kDa | 6 | 4 | 4 | 3 |
| Lmnb2 Isoform B2 of Lamin-B2 | IPI00126191 | 67 kDa | 6 | 4 | 4 | 3 |
| Rps5 20 kDa protein | IPI00857345 | 20 kDa | 5 | 4 | 4 | 3 |
| Napg Gamma-soluble NSF attachment protein | IPI00109870 (+1) | 35 kDa | 5 | 4 | 4 | 3 |
| 2700060E02Rik UPF0568 protein C14orf166 homolog | IPI00132456 | 28 kDa | 5 | 4 | 4 | 3 |
| Psmb3 Proteasome subunit beta type-3 | IPI00314467 | 23 kDa | 4 | 3 | 4 | 3 |
| Hist2h2ac;Hist2h2ab Histone H2A type 2-C | IPI00272033 (+2) | 14 kDa | 4 | 3 | 4 | 3 |
| Snrpd3 Small nuclear ribonucleoprotein Sm D3 | IPI00119224 | 14 kDa | 3 | 3 | 4 | 3 |
| Atp5f1 ATP synthase subunit b, mitochondrial precursor | IPI00341282 | 29 kDa | 3 | 3 | 4 | 3 |
| Letm1 Leucine zipper-EF-hand-containing transmembrane protein 1, mitochondrial precurso | IPI00131177 | 83 kDa | 3 | 3 | 4 | 3 |
| Gm1337 Novel ankyrin repeat domain containing protein | IPI00379202 | 41 kDa | 3 | 2 | 4 | 3 |
| C1qb Complement C1q subcomponent subunit B precursor | IPI00311147 (+1) | 27 kDa | 0 | 0 | 4 | 3 |
| Larp1 la related protein | IPI00344088 | 149 kDa | 0 | 0 | 4 | 3 |
| Eif3a Eukaryotic translation initiation factor 3 subunit A | IPI00129276 | 162 kDa | 0 | 0 | 4 | 3 |
| EG245405 Novel protein silmilar to ribosomal protein L32 | IPI00128267 (+1) | 16 kDa | 0 | 0 | 4 | 3 |
| Rpl15 60S ribosomal protein L15 | IPI00273803 (+4) | 24 kDa | 0 | 0 | 4 | 3 |
| Ppp2r5c MKIAA0044 protein | IPI00408059 | 67 kDa | 0 | 0 | 4 | 3 |
| Homer3 Isoform 1 of Homer protein homolog 3 | IPI00262616 (+1) | 40 kDa | 0 | 0 | 4 | 3 |
| Spock1 Testican-1 precursor | IPI00123549 (+1) | 50 kDa | 0 | 0 | 4 | 3 |
| Traf3 Tnf receptor-associated factor 3 isoform b | IPI00788392 (+1) | 62 kDa | 0 | 0 | 4 | 3 |
| 4932438A13Rik Isoform 1 of Uncharacterized protein KIAA1109 | IPI00755374 (+4) | 555 kDa | 0 | 0 | 4 | 3 |
| D030016E14Rik hypothetical protein LOC320714 | IPI00409255 | 128 kDa | 0 | 0 | 4 | 3 |
| Wtap Isoform 1 of Pre-mRNA-splicing regulator WTAP | IPI00410967 | 44 kDa | 0 | 0 | 4 | 3 |
| Spg3a Atlastin-1 | IPI00221754 (+1) | 63 kDa | 0 | 0 | 4 | 3 |
| Larp5 Isoform 2 of La-related protein 5 | IPI00469235 (+1) | 73 kDa | 0 | 0 | 4 | 3 |
| Copg 8 days embryo whole body cDNA, RIKEN full-length enriched library, clone:5730412D0 | IPI00223437 | 98 kDa | 0 | 0 | 4 | 3 |
| ORF34 Isoform 1 of Constitutive coactivator of PPAR-gamma-like protein 2 | IPI00416122 (+1) | 120 kDa | 16 | 14 | 3 | 3 |
| Myo6 MKIAA0389 protein | IPI00462752 (+2) | 148 kDa | 24 | 8 | 3 | 3 |
| Glg1 Golgi apparatus protein 1 precursor | IPI00122399 | 134 kDa | 7 | 7 | 3 | 3 |
| Ndufa4 NADH dehydrogenase [ubiquinone] 1 alpha subcomplex subunit 4 | IPI00125929 | 9 kDa | 9 | 6 | 3 | 3 |
| 2610301G19Rik Protein KIAA1967 homolog | IPI00123624 | 103 kDa | 7 | 6 | 3 | 3 |
| Clasp2 CLIP-associating protein CLASP2 c | IPI00407863 (+1) | 166 kDa | 7 | 6 | 3 | 3 |

| | | | | | | |
|---|---|---|---|---|---|---|
| Cpsf6 Cleavage and polyadenylation specificity factor subunit 6 | IPI00421085 | 59 kDa | 7 | 6 | 3 | 3 |
| Caprin1 cytoplasmic activation/proliferation-associated protein 1 isoform c | IPI00121515 (+2) | 77 kDa | 6 | 5 | 3 | 3 |
| Pcdh1 Protocadherin 1 | IPI00154057 | 112 kDa | 5 | 4 | 3 | 3 |
| Eif3c Eukaryotic translation initiation factor 3 subunit C | IPI00321647 | 106 kDa | 5 | 4 | 3 | 3 |
| Smc3 Structural maintenance of chromosomes protein 3 | IPI00132122 | 142 kDa | 5 | 4 | 3 | 3 |
| Alg2 Alpha-1,3-mannosyltransferase ALG2 | IPI00121575 | 47 kDa | 4 | 4 | 3 | 3 |
| Prkcsh Isoform 1 of Glucosidase 2 subunit beta precursor | IPI00115680 (+1) | 59 kDa | 4 | 4 | 3 | 3 |
| Arpc4 Actin-related protein 2/3 complex subunit 4 | IPI00138691 | 20 kDa | 4 | 4 | 3 | 3 |
| Trappc3 Trafficking protein particle complex subunit 3 | IPI00115056 | 20 kDa | 5 | 3 | 3 | 3 |
| Cbr1 Carbonyl reductase [NADPH] 1 | IPI00314191 | 31 kDa | 4 | 3 | 3 | 3 |
| Ttc7b similar to tetratricopeptide repeat domain 7B isoform 1 | IPI00754876 (+1) | 100 kDa | 4 | 3 | 3 | 3 |
| Mrps7 28S ribosomal protein S7, mitochondrial precursor | IPI00330688 | 28 kDa | 3 | 3 | 3 | 3 |
| Myef2 Isoform 2 of Myelin expression factor 2 | IPI00154084 (+3) | 62 kDa | 3 | 3 | 3 | 3 |
| Uchl5 Isoform 1 of Ubiquitin carboxyl-terminal hydrolase isozyme L5 | IPI00124938 (+1) | 38 kDa | 3 | 3 | 3 | 3 |
| Syne1 synaptic nuclear envelope 1 isoform 3 | IPI00403993 | 1010 kDa | 3 | 3 | 3 | 3 |
| Nlgn3 Neuroligin-3 precursor | IPI00227168 (+4) | 91 kDa | 3 | 2 | 3 | 3 |
| EG243302 similar to ribosomal protein S25 | IPI00115992 (+5) | 14 kDa | 2 | 2 | 3 | 3 |
| Diras2 GTP-binding protein Di-Ras2 | IPI00126551 | 22 kDa | 2 | 2 | 3 | 3 |
| Snw1 SNW domain-containing protein 1 | IPI00317298 | 61 kDa | 2 | 2 | 3 | 3 |
| Gad1 Glutamate decarboxylase 1 | IPI00318496 | 67 kDa | 2 | 2 | 3 | 3 |
| Hectd3 Isoform 1 of Probable E3 ubiquitin-protein ligase HECTD3 | IPI00223661 | 97 kDa | 2 | 2 | 3 | 3 |
| Pycrl Pyrroline-5-carboxylate reductase 3 | IPI00153234 (+1) | 29 kDa | 2 | 2 | 3 | 3 |
| Exoc5 exocyst complex component 5 | IPI00749893 (+1) | 82 kDa | 2 | 2 | 3 | 3 |
| Map1lc3a Microtubule-associated proteins 1A/1B light chain 3A precursor | IPI00137594 | 14 kDa | 2 | 2 | 3 | 3 |
| Fth1 Ferritin heavy chain | IPI00230145 | 21 kDa | 2 | 2 | 3 | 3 |
| Ccdc92 Coiled-coil domain-containing protein 92 | IPI00123541 | 35 kDa | 2 | 2 | 3 | 3 |
| D10Wsu52e UPF0027 protein C22orf28 homolog | IPI00116850 | 55 kDa | 2 | 2 | 3 | 3 |
| Lgi2 Isoform 2 of Leucine-rich repeat LGI family member 2 precursor | IPI00337001 | 62 kDa | 0 | 0 | 3 | 3 |
| Rpl12 Rpl12 protein (Fragment) | IPI00463634 (+1) | 23 kDa | 0 | 0 | 3 | 3 |
| Pfn2 Isoform 1 of Profilin-2 | IPI00228645 (+1) | 15 kDa | 0 | 0 | 3 | 3 |
| Brcc3 BRCA1/BRCA2-containing complex subunit 3 | IPI00137103 | 33 kDa | 0 | 0 | 3 | 3 |
| Psmd14 26S proteasome non-ATPase regulatory subunit 14 | IPI00113262 | 35 kDa | 0 | 0 | 3 | 3 |
| Ipo7 Importin-7 | IPI00331444 | 119 kDa | 0 | 0 | 3 | 3 |
| Lactb;LOC677144 Serine beta-lactamase-like protein LACTB, mitochondrial precursor | IPI00109293 | 61 kDa | 0 | 0 | 3 | 3 |
| Wdr37 WD repeat-containing protein 37 | IPI00387421 | 55 kDa | 0 | 0 | 3 | 3 |
| Mpp2 Isoform 1 of MAGUK p55 subfamily member 2 | IPI00125147 (+1) | 62 kDa | 0 | 0 | 3 | 3 |
| Dsp desmoplakin | IPI00553419 (+2) | 333 kDa | 0 | 0 | 3 | 3 |
| Exoc4 Exocyst complex component 4 | IPI00129959 | 111 kDa | 0 | 0 | 3 | 3 |
| Rpl19;LOC100046041 60S ribosomal protein L19 | IPI00122426 (+3) | 23 kDa | 0 | 0 | 3 | 3 |
| Rps7 40S ribosomal protein S7 | IPI00136984 (+2) | 22 kDa | 0 | 0 | 3 | 3 |
| Vat1 Synaptic vesicle membrane protein VAT-1 homolog | IPI00126072 | 43 kDa | 0 | 0 | 3 | 3 |

| | | | | | | |
|---|---|---|---|---|---|---|
| Atp5j2 ATP synthase subunit f, mitochondrial | IPI00271986 | 10 kDa | 0 | 0 | 3 | 3 |
| OTTMUSG00000014710;Rpl29;EG666642 60S ribosomal protein L29 | IPI00222548 (+4) | 18 kDa | 0 | 0 | 3 | 3 |
| Exoc1 Exocyst complex component 1 | IPI00153813 (+2) | 103 kDa | 0 | 0 | 3 | 3 |
| Hnrpll Isoform 1 of Heterogeneous nuclear ribonucleoprotein L-like | IPI00121760 | 64 kDa | 0 | 0 | 3 | 3 |
| Polr2b DNA-directed RNA polymerase II subunit RPB2 | IPI00320034 | 134 kDa | 0 | 0 | 3 | 3 |
| Ap1g1 14 days pregnant adult female placenta cDNA, RIKEN full-length enriched library, clor | IPI00224151 (+1) | 92 kDa | 0 | 0 | 3 | 3 |
| Usf1 Upstream stimulatory factor 1 | IPI00117062 | 34 kDa | 0 | 0 | 3 | 3 |
| Rpl8 60S ribosomal protein L8 | IPI00137787 | 28 kDa | 0 | 0 | 3 | 3 |
| Cdk5 Cell division protein kinase 5 | IPI00309262 | 33 kDa | 0 | 0 | 3 | 3 |
| Cse1l Exportin-2 | IPI00112414 | 110 kDa | 0 | 0 | 3 | 3 |
| Fxr2 Fragile X mental retardation, autosomal homolog 2 | IPI00126389 (+1) | 74 kDa | 0 | 0 | 3 | 3 |
| Ddx1 ATP-dependent RNA helicase DDX1 | IPI00127172 | 83 kDa | 0 | 0 | 3 | 3 |
| Actr3 Actin-related protein 3 | IPI00115627 | 47 kDa | 0 | 0 | 3 | 3 |
| Bre Isoform 2 of Protein BRE | IPI00128466 (+2) | 44 kDa | 0 | 0 | 3 | 3 |
| Trappc9 Isoform 2 of NIK- and IKBKB-binding protein | IPI00454019 (+3) | 127 kDa | 0 | 0 | 3 | 3 |
| Gpd1 Glycerol-3-phosphate dehydrogenase [NAD+], cytoplasmic | IPI00230185 | 38 kDa | 0 | 0 | 3 | 3 |
| Rpl11 60S ribosomal protein L11 | IPI00331461 (+2) | 20 kDa | 0 | 0 | 3 | 3 |
| Mrpl22 39S ribosomal protein L22, mitochondrial precursor | IPI00225318 | 24 kDa | 0 | 0 | 3 | 3 |
| Ptcd3 CRL-1722 L5178Y-R cDNA, RIKEN full-length enriched library, clone:I730029A14 produ | IPI00338458 | 78 kDa | 0 | 0 | 3 | 3 |
| Gdi1 Rab GDP dissociation inhibitor alpha | IPI00323179 | 51 kDa | 0 | 0 | 3 | 3 |
| Gdap1l1 Ganglioside-induced differentiation-associated protein 1-like 1 | IPI00123756 (+1) | 42 kDa | 0 | 0 | 3 | 3 |
| Zc3h14 Isoform 2 of Zinc finger CCCH domain-containing protein 14 | IPI00380641 (+4) | 65 kDa | 0 | 0 | 3 | 3 |
| Scn2a1 sodium channel, voltage-gated, type II, alpha 1 | IPI00761641 | 228 kDa | 0 | 0 | 3 | 3 |
| Nars Asparaginyl-tRNA synthetase, cytoplasmic | IPI00223415 | 64 kDa | 0 | 0 | 3 | 3 |
| Sart3 Isoform 1 of Squamous cell carcinoma antigen recognized by T-cells 3 | IPI00458854 | 110 kDa | 0 | 0 | 3 | 3 |
| Mycbp2 Isoform 2 of Probable E3 ubiquitin-protein ligase MYCBP2 | IPI00380409 (+1) | 517 kDa | 0 | 0 | 3 | 3 |
| Supt6h Transcription elongation factor SPT6 | IPI00454050 | 199 kDa | 0 | 0 | 3 | 3 |
| Acadl acyl-Coenzyme A dehydrogenase, long-chain | IPI00894588 | 48 kDa | 0 | 0 | 3 | 3 |
| Wdr57 WD repeat-containing protein 57 | IPI00461621 | 39 kDa | 0 | 0 | 3 | 3 |
| Mars Methionyl-tRNA synthetase, cytoplasmic | IPI00461469 | 101 kDa | 0 | 0 | 3 | 3 |
| Srp68 Signal recognition particle 68 kDa protein | IPI00222977 (+2) | 71 kDa | 0 | 0 | 3 | 3 |
| Nol5a Nucleolar protein 5A | IPI00318048 | 64 kDa | 0 | 0 | 3 | 3 |
| Rpusd4 RNA pseudouridylate synthase domain-containing protein 4 | IPI00109744 | 42 kDa | 0 | 0 | 3 | 3 |
| Ccdc22 Coiled-coil domain-containing protein 22 | IPI00331244 | 71 kDa | 0 | 0 | 3 | 3 |
| Larp7 Isoform 1 of La-related protein 7 | IPI00340860 | 65 kDa | 0 | 0 | 3 | 3 |
| Wdr47 WD repeat-containing protein 47 | IPI00330193 (+1) | 102 kDa | 0 | 0 | 3 | 3 |
| EG436081 hypothetical protein | IPI00463297 (+3) | 12 kDa | 0 | 0 | 3 | 3 |
| Crocc Isoform 1 of Rootletin | IPI00229917 (+3) | 227 kDa | 0 | 0 | 3 | 3 |
| Rps15 40S ribosomal protein S15 | IPI00319231 (+3) | 17 kDa | 0 | 0 | 3 | 3 |
| Matn4 Isoform Long of Matrilin-4 precursor | IPI00130665 (+1) | 69 kDa | 0 | 0 | 3 | 3 |
| Gapvd1 Isoform 2 of GTPase-activating protein and VPS9 domain-containing protein 1 | IPI00406596 (+4) | 160 kDa | 0 | 0 | 3 | 3 |

| | | | | | | |
|---|---|---|---|---|---|---|
| Adam11 a disintegrin and metallopeptidase domain 11 isoform 2 | IPI00408232 (+1) | 84 kDa | 0 | 0 | 3 | 3 |
| Rhot1 Isoform 3 of Mitochondrial Rho GTPase 1 | IPI00123186 (+3) | 77 kDa | 0 | 0 | 3 | 3 |
| Nomo1 Nodal modulator 1 | IPI00222429 | 133 kDa | 0 | 0 | 3 | 3 |
| Tuba1a Tubulin alpha-1A chain | IPI00110753 | 50 kDa | 324 | 2 | 274 | 2 |
| Spire1 Spir-1 protein isoform 1 | IPI00380020 | 74 kDa | 0 | 0 | 256 | 2 |
| AU042671 similar to chromosome 12 open reading frame 51 | IPI00808443 | 456 kDa | 0 | 0 | 186 | 2 |
| Dnm1 Isoform 1 of Dynamin-1 | IPI00272878 (+2) | 98 kDa | 0 | 0 | 95 | 2 |
| Stxbp1 Isoform 2 of Syntaxin-binding protein 1 | IPI00415403 | 69 kDa | 0 | 0 | 40 | 2 |
| Actbl2 Beta-actin-like protein 2 | IPI00221528 | 42 kDa | 0 | 0 | 31 | 2 |
| Eef1a2 Elongation factor 1-alpha 2 | IPI00119667 | 50 kDa | 0 | 0 | 30 | 2 |
| Mtap4 Isoform 4 of Microtubule-associated protein 4 | IPI00406741 | 95 kDa | 14 | 2 | 24 | 2 |
| sp_K1C15_SHEEP | IPIsp_K1C15_SHEEP | 49 kDa | 20 | 2 | 14 | 2 |
| Prkacb Isoform 1 of cAMP-dependent protein kinase catalytic subunit beta | IPI00263822 (+3) | 41 kDa | 6 | 2 | 14 | 2 |
| Krt76 Adult female vagina cDNA, RIKEN full-length enriched library, clone:9930024P18 prod | IPI00346834 | 63 kDa | 0 | 0 | 13 | 2 |
| Actn4 Alpha-actinin-4 | IPI00118899 | 105 kDa | 20 | 5 | 12 | 2 |
| Spnb4 Beta4-spectrin | IPI00270149 (+1) | 288 kDa | 25 | 4 | 12 | 2 |
| Krt42 Keratin, type I cytoskeletal 42 | IPI00468696 | 50 kDa | 0 | 0 | 12 | 2 |
| Tnik Isoform 1 of Traf2 and NCK-interacting protein kinase | IPI00187510 (+6) | 150 kDa | 0 | 0 | 10 | 2 |
| Rab8a Ras-related protein Rab-8A | IPI00331128 (+1) | 24 kDa | 0 | 0 | 10 | 2 |
| Gnai2 Guanine nucleotide-binding protein G(i), alpha-2 subunit | IPI00228617 (+1) | 40 kDa | 0 | 0 | 9 | 2 |
| Luc7l Isoform 1 of Putative RNA-binding protein Luc7-like 1 | IPI00410804 (+3) | 44 kDa | 0 | 0 | 8 | 2 |
| sp_PPIA_HUMAN | IPIsp_PPIA_HUMAN | 18 kDa | 7 | 2 | 7 | 2 |
| Rab1b Ras-related protein Rab-1B | IPI00133706 | 22 kDa | 0 | 0 | 7 | 2 |
| Asna1 Arsenical pump-driving ATPase | IPI00624501 | 39 kDa | 3 | 2 | 6 | 2 |
| Ppp3cb Isoform 2 of Serine/threonine-protein phosphatase 2B catalytic subunit beta isoforr | IPI00112312 (+2) | 59 kDa | 0 | 0 | 6 | 2 |
| Ppfia3 115 kDa protein | IPI00857748 | 115 kDa | 29 | 4 | 5 | 2 |
| Clu Clusterin precursor | IPI00320420 (+1) | 52 kDa | 9 | 3 | 5 | 2 |
| Rab6b Ras-related protein Rab-6B | IPI00378145 | 23 kDa | 6 | 3 | 5 | 2 |
| Bcas1 Bcas1 protein | IPI00756676 | 62 kDa | 3 | 3 | 5 | 2 |
| Rab12 RAB12, member RAS oncogene family | IPI00169699 | 32 kDa | 0 | 0 | 5 | 2 |
| Ap2s1 AP-2 complex subunit sigma-1 | IPI00118022 | 17 kDa | 7 | 5 | 4 | 2 |
| Cisd1 CDGSH iron sulfur domain-containing protein 1 | IPI00128346 | 12 kDa | 8 | 3 | 4 | 2 |
| Cadm2 Isoform 2 of Cell adhesion molecule 2 precursor | IPI00453537 (+3) | 44 kDa | 2 | 2 | 4 | 2 |
| Hist1h2bl;Hist1h2bf;Hist1h2bn;Hist1h2bj;LOC100046213 Histone H2B type 1-F/J/L | IPI00114642 (+20) | 14 kDa | 2 | 2 | 4 | 2 |
| Cad carbamoyl-phosphate synthetase 2, aspartate transcarbamylase, and dihydroorotase | IPI00380280 (+1) | 243 kDa | 0 | 0 | 4 | 2 |
| Ndrg2 Isoform 1 of Protein NDRG2 | IPI00136134 (+1) | 41 kDa | 0 | 0 | 4 | 2 |
| Podxl2 Isoform 3 of Podocalyxin-like protein 2 precursor | IPI00229420 (+1) | 58 kDa | 0 | 0 | 4 | 2 |
| Dynll1 Dynein light chain 1, cytoplasmic | IPI00121623 | 10 kDa | 0 | 0 | 4 | 2 |
| Jup Junction plakoglobin | IPI00229475 | 82 kDa | 0 | 0 | 4 | 2 |
| Dpp10 Inactive dipeptidyl peptidase 10 | IPI00396687 (+1) | 91 kDa | 0 | 0 | 4 | 2 |
| ENSMUSG00000042718;Fem1a Protein fem-1 homolog A-A | IPI00311229 (+1) | 72 kDa | 0 | 0 | 4 | 2 |

| | | | | | | |
|---|---|---|---|---|---|---|
| Tsc22d2 TSC22 domain family 2 | IPI00762599 | 78 kDa | 0 | 0 | 4 | 2 |
| Srgap3 Isoform 1 of SLIT-ROBO Rho GTPase-activating protein 3 | IPI00330039 (+3) | 124 kDa | 10 | 9 | 3 | 2 |
| Mapre3 Microtubule-associated protein RP/EB family member 3 | IPI00343557 | 32 kDa | 7 | 4 | 3 | 2 |
| Snrpf small nuclear ribonucleoprotein polypeptide F | IPI00117371 (+2) | 20 kDa | 6 | 3 | 3 | 2 |
| Rala Ras-related protein Ral-A precursor | IPI00124282 | 24 kDa | 4 | 3 | 3 | 2 |
| Oxr1 Isoform 2 of Oxidation resistance protein 1 | IPI00117929 (+1) | 86 kDa | 3 | 3 | 3 | 2 |
| Pgrmc1 Membrane-associated progesterone receptor component 1 | IPI00319973 | 22 kDa | 3 | 3 | 3 | 2 |
| 1700054N08Rik Uncharacterized protein C1orf96 homolog | IPI00187478 | 28 kDa | 3 | 3 | 3 | 2 |
| AU040829 Isoform 1 of Rho GTPase-activating protein RICH2 | IPI00128632 (+2) | 89 kDa | 3 | 3 | 3 | 2 |
| Syngr1 Isoform 1A of Synaptogyrin-1 | IPI00115763 (+1) | 26 kDa | 8 | 2 | 3 | 2 |
| Psmc3 26S protease regulatory subunit 6A | IPI00133206 (+2) | 49 kDa | 4 | 2 | 3 | 2 |
| Gng3 Guanine nucleotide-binding protein G(I)/G(S)/G(O) subunit gamma-3 precursor | IPI00129268 | 8 kDa | 3 | 2 | 3 | 2 |
| Nap1l4 Bone marrow stroma cell CRL-2028 SR-4987 cDNA, RIKEN full-length enriched library | IPI00133977 (+1) | 47 kDa | 3 | 2 | 3 | 2 |
| Lars Leucyl-tRNA synthetase, cytoplasmic | IPI00453819 | 134 kDa | 3 | 2 | 3 | 2 |
| Uchl1 Ubiquitin carboxyl-terminal hydrolase isozyme L1 | IPI00313962 (+1) | 25 kDa | 2 | 2 | 3 | 2 |
| Ranbp1 Ran-specific GTPase-activating protein (Fragment) | IPI00321978 | 24 kDa | 2 | 2 | 3 | 2 |
| Gfer growth factor, erv1 (S. cerevisiae)-like | IPI00410978 | 23 kDa | 2 | 2 | 3 | 2 |
| mt-Atp8 ATP synthase protein 8 | IPI00116896 | 8 kDa | 2 | 2 | 3 | 2 |
| Atp5l Adult male small intestine cDNA, RIKEN full-length enriched library, clone:2010009O0 | IPI00133342 (+1) | 14 kDa | 2 | 2 | 3 | 2 |
| Arl8a ADP-ribosylation factor-like protein 8A | IPI00124610 (+2) | 21 kDa | 0 | 0 | 3 | 2 |
| Pfkp Isoform 1 of 6-phosphofructokinase type C | IPI00124444 (+1) | 85 kDa | 0 | 0 | 3 | 2 |
| Rpl13a 60S ribosomal protein L13a | IPI00223217 (+2) | 23 kDa | 0 | 0 | 3 | 2 |
| Vps29 Isoform 1 of Vacuolar protein sorting-associated protein 29 | IPI00136936 (+3) | 20 kDa | 0 | 0 | 3 | 2 |
| Dctn4 Isoform 2 of Dynactin subunit 4 | IPI00317289 (+1) | 52 kDa | 0 | 0 | 3 | 2 |
| Gpi1 Glucose-6-phosphate isomerase | IPI00228633 | 63 kDa | 0 | 0 | 3 | 2 |
| Rps27;ENSMUSG00000050621;LOC100047537 40S ribosomal protein S27 | IPI00173160 (+1) | 9 kDa | 0 | 0 | 3 | 2 |
| Mtap9 Isoform 2 of Microtubule-associated protein 9 | IPI00625151 | 70 kDa | 0 | 0 | 3 | 2 |
| Sv2b Synaptic vesicle glycoprotein 2B | IPI00221456 | 77 kDa | 0 | 0 | 3 | 2 |
| Eif4a2 Isoform 1 of Eukaryotic initiation factor 4A-II | IPI00400432 (+2) | 46 kDa | 0 | 0 | 3 | 2 |
| ENSMUSG00000066443;LOC100046290;OTTMUSG00000017565;LOC100044606;OTTMUSG | IPI00315548 (+9) | 19 kDa | 0 | 0 | 3 | 2 |
| Rab11fip2 NOD-derived CD11c +ve dendritic cells cDNA, RIKEN full-length enriched library, c | IPI00281670 | 60 kDa | 0 | 0 | 3 | 2 |
| Flnb Filamin-B | IPI00663627 | 278 kDa | 0 | 0 | 3 | 2 |
| Mapk8ip3 Isoform 1a of C-jun-amino-terminal kinase-interacting protein 3 | IPI00230497 (+7) | 144 kDa | 0 | 0 | 3 | 2 |
| Rpl14 60S ribosomal protein L14 | IPI00133185 (+1) | 24 kDa | 0 | 0 | 3 | 2 |
| Acot11 Acyl-coenzyme A thioesterase 11 | IPI00128588 (+1) | 67 kDa | 0 | 0 | 3 | 2 |
| Rdh14 Retinol dehydrogenase 14 | IPI00112377 | 36 kDa | 0 | 0 | 3 | 2 |
| Coro1c Coronin-1C | IPI00124820 | 53 kDa | 0 | 0 | 3 | 2 |
| LOC100044107 similar to Guanine nucleotide binding protein, alpha z subunit | IPI00850311 (+1) | 42 kDa | 0 | 0 | 3 | 2 |
| Fbxo3 F-box only protein 3 | IPI00120202 (+2) | 55 kDa | 0 | 0 | 3 | 2 |
| H2afy Isoform 1 of Core histone macro-H2A.1 | IPI00137852 (+1) | 39 kDa | 0 | 0 | 3 | 2 |
| Snrp70 Isoform 1 of U1 small nuclear ribonucleoprotein 70 kDa | IPI00625105 | 52 kDa | 0 | 0 | 3 | 2 |

| | | | | | | |
|---|---|---|---|---|---|---|
| Plxdc1 Isoform 1 of Plexin domain-containing protein 1 precursor | IPI00222883 | 56 kDa | 0 | 0 | 3 | 2 |
| Zfp828 Zinc finger protein 828 | IPI00453800 (+1) | 88 kDa | 0 | 0 | 3 | 2 |
| Dnaja4 DnaJ homolog subfamily A member 4 | IPI00125454 | 45 kDa | 0 | 0 | 3 | 2 |
| Nlgn2 Neuroligin-2 precursor | IPI00468605 | 91 kDa | 0 | 0 | 3 | 2 |
| Llgl1 116 kDa protein | IPI00380354 (+1) | 116 kDa | 0 | 0 | 3 | 2 |
| Cul3 Cullin-3 | IPI00467383 | 89 kDa | 42 | 31 | 2 | 2 |
| Cuedc1 CUE domain-containing protein 1 | IPI00321773 (+1) | 43 kDa | 28 | 12 | 2 | 2 |
| Ak5 adenylate kinase 5 | IPI00116072 | 63 kDa | 11 | 8 | 2 | 2 |
| Clmn Isoform 4 of Calmin | IPI00331290 (+3) | 113 kDa | 7 | 7 | 2 | 2 |
| Tpr Nuclear pore complex-associated intranuclear coiled-coil protein TPR | IPI00880644 | 267 kDa | 6 | 6 | 2 | 2 |
| Cdh2 Cadherin-2 precursor | IPI00323134 (+2) | 100 kDa | 5 | 5 | 2 | 2 |
| Rnf126 RING finger protein 126 | IPI00130263 (+1) | 34 kDa | 8 | 4 | 2 | 2 |
| Rac1 Mammary gland RCB-0527 Jyg-MC(B) cDNA, RIKEN full-length enriched library, clone:G | IPI00127408 (+1) | 23 kDa | 4 | 4 | 2 | 2 |
| Chchd3 Coiled-coil-helix-coiled-coil-helix domain-containing protein 3, mitochondrial precur | IPI00133562 | 26 kDa | 4 | 4 | 2 | 2 |
| Ndufs3 NADH dehydrogenase [ubiquinone] iron-sulfur protein 3, mitochondrial precursor | IPI00121309 | 30 kDa | 4 | 4 | 2 | 2 |
| Dnajb11 DnaJ homolog subfamily B member 11 precursor | IPI00320241 | 41 kDa | 4 | 4 | 2 | 2 |
| Eps15l1 Isoform 1 of Epidermal growth factor receptor substrate 15-like 1 | IPI00420185 (+4) | 99 kDa | 4 | 4 | 2 | 2 |
| Dlg1;LOC100047603 Isoform 3 of Disks large homolog 1 | IPI00125861 (+3) | 103 kDa | 4 | 4 | 2 | 2 |
| Slk Isoform 1 of STE20-like serine/threonine-protein kinase | IPI00331076 (+1) | 141 kDa | 4 | 4 | 2 | 2 |
| Nme2 Nucleoside diphosphate kinase B | IPI00127417 (+4) | 17 kDa | 5 | 3 | 2 | 2 |
| Psmd8 26S proteasome non-ATPase regulatory subunit 8 | IPI00403509 (+1) | 30 kDa | 4 | 3 | 2 | 2 |
| Cops7a Isoform 1 of COP9 signalosome complex subunit 7a | IPI00123465 (+2) | 30 kDa | 4 | 3 | 2 | 2 |
| Dnajb2 Isoform 2 of DnaJ homolog subfamily B member 10 | IPI00136216 (+2) | 29 kDa | 4 | 3 | 2 | 2 |
| Wasf3 Wiskott-Aldrich syndrome protein family member 3 | IPI00128341 | 55 kDa | 4 | 3 | 2 | 2 |
| Srgap2 124 kDa protein | IPI00652316 (+1) | 124 kDa | 3 | 3 | 2 | 2 |
| Rab5c Ras-related protein Rab-5C | IPI00224518 (+1) | 23 kDa | 3 | 3 | 2 | 2 |
| Skp1a S-phase kinase-associated protein 1 | IPI00331163 | 19 kDa | 3 | 3 | 2 | 2 |
| Atp6v1g1 Vacuolar proton pump subunit G 1 | IPI00133163 | 14 kDa | 3 | 3 | 2 | 2 |
| Gng2 Guanine nucleotide-binding protein G(I)/G(S)/G(O) subunit gamma-2 precursor | IPI00230194 | 8 kDa | 3 | 3 | 2 | 2 |
| Sod2 Superoxide dismutase [Mn], mitochondrial precursor | IPI00109109 | 25 kDa | 3 | 3 | 2 | 2 |
| Ccdc128 Isoform 1 of Coiled-coil domain-containing protein 128 | IPI00308333 (+1) | 88 kDa | 3 | 3 | 2 | 2 |
| Gpd1l Isoform 1 of Glycerol-3-phosphate dehydrogenase 1-like protein | IPI00336807 (+1) | 38 kDa | 3 | 3 | 2 | 2 |
| Prdx6 Peroxiredoxin-6 | IPI00555059 (+2) | 25 kDa | 3 | 3 | 2 | 2 |
| Fbxo21 Isoform 1 of F-box only protein 21 | IPI00123296 (+1) | 72 kDa | 3 | 3 | 2 | 2 |
| Thy1 Thy-1 membrane glycoprotein precursor | IPI00109727 | 18 kDa | 5 | 2 | 2 | 2 |
| Ppfia2 Liprin-alpha-2 | IPI00224775 (+1) | 143 kDa | 5 | 2 | 2 | 2 |
| Dnaja3 Isoform 3 of DnaJ homolog subfamily A member 3, mitochondrial precursor | IPI00120414 (+2) | 47 kDa | 4 | 2 | 2 | 2 |
| Tppp Tubulin polymerization-promoting protein | IPI00119067 | 24 kDa | 2 | 2 | 2 | 2 |
| Echs1 Enoyl-CoA hydratase, mitochondrial precursor | IPI00454049 | 31 kDa | 2 | 2 | 2 | 2 |
| Ggt7 Gamma-glutamyltransferase 7 precursor | IPI00115429 (+1) | 70 kDa | 2 | 2 | 2 | 2 |
| Ubqln2 Ubiquilin-2 | IPI00387416 (+1) | 67 kDa | 2 | 2 | 2 | 2 |

| | | | | | | |
|---|---|---|---|---|---|---|
| Mrpl12 39S ribosomal protein L12, mitochondrial precursor | IPI00118963 | 22 kDa | 2 | 2 | 2 | 2 |
| Uqcc Isoform 1 of Ubiquinol-cytochrome c reductase complex chaperone CBP3 homolog | IPI00109603 (+3) | 34 kDa | 2 | 2 | 2 | 2 |
| Lonp1 Lon protease homolog, mitochondrial precursor | IPI00761408 | 106 kDa | 2 | 2 | 2 | 2 |
| 1500005I02Rik BTB/POZ and BACK domain-containing protein LOC388419 homolog precurs | IPI00119130 | 53 kDa | 2 | 2 | 2 | 2 |
| Txndc10 Protein disulfide-isomerase TXNDC10 precursor | IPI00453798 | 52 kDa | 2 | 2 | 2 | 2 |
| OTTMUSG00000022083;Rps23;LOC100046668 11 days embryo whole body cDNA, RIKEN ful | IPI00131357 | 16 kDa | 2 | 2 | 2 | 2 |
| Rps15a 40S ribosomal protein S15a | IPI00230660 (+6) | 15 kDa | 2 | 2 | 2 | 2 |
| Eif4e2 Eukaryotic translation initiation factor 4E type 2 | IPI00222983 (+8) | 28 kDa | 0 | 0 | 2 | 2 |
| Mrps36 Mitochondrial 28S ribosomal protein S36 | IPI00315808 (+1) | 11 kDa | 0 | 0 | 2 | 2 |
| Ddhd1 Phosphatidic acid-preferring phospholipase A1 variant 1 | IPI00377614 (+2) | 95 kDa | 0 | 0 | 2 | 2 |
| Farp1 FERMRhoGEF (Arhgef) and pleckstrin domain protein 1 | IPI00356904 (+1) | 119 kDa | 0 | 0 | 2 | 2 |
| EG668829 similar to ribosomal protein L24 | IPI00134202 (+3) | 12 kDa | 0 | 0 | 2 | 2 |
| Rpl17 60S ribosomal protein L17 | IPI00453768 (+8) | 21 kDa | 0 | 0 | 2 | 2 |
| Hnrnpab Heterogeneous nuclear ribonucleoprotein A/B | IPI00117288 (+2) | 31 kDa | 0 | 0 | 2 | 2 |
| Cab39 Calcium-binding protein 39 | IPI00318778 (+1) | 40 kDa | 0 | 0 | 2 | 2 |
| Usp46 Ubiquitin carboxyl-terminal hydrolase 46 | IPI00228384 (+1) | 42 kDa | 0 | 0 | 2 | 2 |
| Acaa2 3-ketoacyl-CoA thiolase, mitochondrial | IPI00226430 (+1) | 42 kDa | 0 | 0 | 2 | 2 |
| Kpna4 Importin subunit alpha-4 | IPI00129792 | 58 kDa | 0 | 0 | 2 | 2 |
| Bsg Isoform 2 of Basigin precursor | IPI00113869 (+2) | 30 kDa | 0 | 0 | 2 | 2 |
| Dnpep aspartyl aminopeptidase isoform a | IPI00331394 (+1) | 52 kDa | 0 | 0 | 2 | 2 |
| U2af2 Splicing factor U2AF 65 kDa subunit | IPI00113746 | 54 kDa | 0 | 0 | 2 | 2 |
| Pddc1 Parkinson disease 7 domain-containing protein 1 precursor | IPI00224733 | 28 kDa | 0 | 0 | 2 | 2 |
| Cdc23 Cell division cycle protein 23 homolog | IPI00221793 | 69 kDa | 0 | 0 | 2 | 2 |
| Mfn2 Isoform 1 of Mitofusin-2 | IPI00312244 | 86 kDa | 0 | 0 | 2 | 2 |
| Hist1h1c Histone H1.2 | IPI00223713 (+2) | 21 kDa | 0 | 0 | 2 | 2 |
| Odz4 Isoform 3 of Teneurin-4 | IPI00157497 (+3) | 315 kDa | 0 | 0 | 2 | 2 |
| Usp5 Ubiquitin carboxyl-terminal hydrolase 5 | IPI00113214 (+1) | 96 kDa | 0 | 0 | 2 | 2 |
| Ncl Nucleolin | IPI00317794 | 77 kDa | 0 | 0 | 2 | 2 |
| Rab3gap2 similar to Rab3 GTPase-activating protein non-catalytic subunit (Rab3 GTPase-act | IPI00404477 | 155 kDa | 0 | 0 | 2 | 2 |
| Elmo2 Isoform 1 of Engulfment and cell motility protein 2 | IPI00402900 (+2) | 84 kDa | 0 | 0 | 2 | 2 |
| LOC100048600;Erc1 Isoform 1 of ELKS/RAB6-interacting/CAST family member 1 | IPI00117731 (+2) | 128 kDa | 0 | 0 | 2 | 2 |
| Reps1 RalBP1 associated Eps domain containing 1 isoform 1 | IPI00119795 (+1) | 87 kDa | 0 | 0 | 2 | 2 |
| Pik3r4 Phosphoinositide 3-kinase regulatory subunit 4 | IPI00406045 | 153 kDa | 0 | 0 | 2 | 2 |
| Calb1 Calbindin | IPI00331066 | 30 kDa | 0 | 0 | 2 | 2 |
| Gbl Target of rapamycin complex subunit LST8 | IPI00458055 | 36 kDa | 0 | 0 | 2 | 2 |
| Ppih;LOC629952;OTTMUSG00000002349 Isoform 1 of Peptidyl-prolyl cis-trans isomerase H | IPI00112065 (+4) | 20 kDa | 0 | 0 | 2 | 2 |
| Tbc1d10b TBC1 domain family, member 10b | IPI00469012 | 87 kDa | 0 | 0 | 2 | 2 |
| Kcna1 Potassium voltage-gated channel subfamily A member 1 | IPI00133719 | 56 kDa | 0 | 0 | 2 | 2 |
| Eif4b Eukaryotic translation initiation factor 4B | IPI00221581 | 69 kDa | 0 | 0 | 2 | 2 |
| Rbbp7 Histone-binding protein RBBP7 | IPI00122698 (+2) | 48 kDa | 0 | 0 | 2 | 2 |
| 2410002F23Rik Activated spleen cDNA, RIKEN full-length enriched library, clone:F830206M( | IPI00109705 | 32 kDa | 0 | 0 | 2 | 2 |

| | | | | | | |
|---|---|---|---|---|---|---|
| Mrps2 Mitochondrial 28S ribosomal protein S2 | IPI00126011 | 32 kDa | 0 | 0 | 2 | 2 |
| Rab2a Ras-related protein Rab-2A | IPI00137227 | 24 kDa | 0 | 0 | 2 | 2 |
| Cdkn2aip CDKN2A-interacting protein | IPI00263028 | 60 kDa | 0 | 0 | 2 | 2 |
| Centg3 Isoform 1 of Centaurin-gamma-3 | IPI00128319 (+1) | 98 kDa | 0 | 0 | 2 | 2 |
| Wwox Isoform 1 of WW domain-containing oxidoreductase | IPI00331266 | 47 kDa | 0 | 0 | 2 | 2 |
| Eif2s1 Eukaryotic translation initiation factor 2 subunit 1 | IPI00474446 | 36 kDa | 0 | 0 | 2 | 2 |
| Strbp Isoform 1 of Spermatid perinuclear RNA-binding protein | IPI00128254 (+1) | 74 kDa | 0 | 0 | 2 | 2 |
| Slc25a11 Mitochondrial 2-oxoglutarate/malate carrier protein | IPI00230754 (+1) | 34 kDa | 0 | 0 | 2 | 2 |
| Wdr68 WD repeat-containing protein 68 | IPI00262605 (+1) | 39 kDa | 0 | 0 | 2 | 2 |
| Ppp2r2a Serine/threonine-protein phosphatase 2A 55 kDa regulatory subunit B alpha isofor | IPI00667471 | 52 kDa | 0 | 0 | 2 | 2 |
| Fhl1 Isoform 1 of Four and a half LIM domains protein 1 | IPI00309997 (+5) | 32 kDa | 0 | 0 | 2 | 2 |
| Por NADPH--cytochrome P450 reductase | IPI00621548 (+1) | 77 kDa | 0 | 0 | 2 | 2 |
| Grpel1 GrpE protein homolog 1, mitochondrial precursor | IPI00117083 | 24 kDa | 0 | 0 | 2 | 2 |
| Prpsap1 similar to likely ortholog of H. sapiens phosphoribosyl pyrophosphate synthetase-as | IPI00315538 (+1) | 60 kDa | 0 | 0 | 2 | 2 |
| Rps29;LOC637599;OTTMUSG00000016246 40S ribosomal protein S29 | IPI00222553 | 7 kDa | 0 | 0 | 2 | 2 |
| Sez6l Sez6l protein (Fragment) | IPI00605000 (+3) | 54 kDa | 0 | 0 | 2 | 2 |
| Ppp1r12a Ppp1r12a protein | IPI00671847 (+1) | 115 kDa | 0 | 0 | 2 | 2 |
| Ugp2 Isoform 1 of UTP--glucose-1-phosphate uridylyltransferase | IPI00131204 (+1) | 57 kDa | 0 | 0 | 2 | 2 |
| Hapln2 Hyaluronan and proteoglycan link protein 2 precursor | IPI00113410 | 38 kDa | 0 | 0 | 2 | 2 |
| Tars Threonyl-tRNA synthetase, cytoplasmic | IPI00468688 | 83 kDa | 0 | 0 | 2 | 2 |
| Car2 Carbonic anhydrase 2 | IPI00121534 | 29 kDa | 0 | 0 | 2 | 2 |
| Arcn1 Coatomer subunit delta | IPI00313558 (+1) | 57 kDa | 0 | 0 | 2 | 2 |
| Acadvl Very long-chain specific acyl-CoA dehydrogenase, mitochondrial precursor | IPI00119203 | 71 kDa | 0 | 0 | 2 | 2 |
| Gpx4 Isoform Mitochondrial of Phospholipid hydroperoxide glutathione peroxidase, mitocho | IPI00117281 (+1) | 22 kDa | 0 | 0 | 2 | 2 |
| Rps24 Isoform 2 of 40S ribosomal protein S24 | IPI00402981 (+3) | 15 kDa | 0 | 0 | 2 | 2 |
| Alcam CD166 antigen precursor | IPI00121378 | 65 kDa | 0 | 0 | 2 | 2 |
| OTTMUSG00000016037 similar to 60S ribosomal protein L9 | IPI00117610 (+7) | 22 kDa | 0 | 0 | 2 | 2 |
| Rps21 40S ribosomal protein S21 | IPI00132950 | 9 kDa | 0 | 0 | 2 | 2 |
| Sf1 Isoform CW17 of Splicing factor 1 | IPI00116284 (+4) | 70 kDa | 0 | 0 | 2 | 2 |
| 9030624J02Rik Isoform 2 of UPF0505 protein C16orf62 homolog | IPI00135151 (+2) | 48 kDa | 0 | 0 | 2 | 2 |
| Prpf4 U4/U6 small nuclear ribonucleoprotein Prp4 | IPI00458908 | 58 kDa | 0 | 0 | 2 | 2 |
| Mccc2 Methylcrotonoyl-CoA carboxylase beta chain, mitochondrial precursor | IPI00553717 | 61 kDa | 0 | 0 | 2 | 2 |
| 2500003M10Rik Isoform 2 of Uncharacterized protein C1orf77 homolog | IPI00120174 (+7) | 24 kDa | 0 | 0 | 2 | 2 |
| H2-Ke6 Isoform Short of Estradiol 17-beta-dehydrogenase 8 | IPI00115598 (+1) | 27 kDa | 0 | 0 | 2 | 2 |
| Cadm3 Cell adhesion molecule 3 precursor | IPI00118020 (+1) | 43 kDa | 0 | 0 | 2 | 2 |
| - 68 kDa protein | IPI00229697 (+2) | 68 kDa | 0 | 0 | 2 | 2 |
| Fn3k Fructosamine-3-kinase | IPI00111519 | 35 kDa | 0 | 0 | 2 | 2 |
| Eif3e Eukaryotic translation initiation factor 3 subunit E | IPI00132250 | 52 kDa | 0 | 0 | 2 | 2 |
| Hdac2 Histone deacetylase 2 | IPI00137668 (+2) | 55 kDa | 0 | 0 | 2 | 2 |
| Pdk3 [Pyruvate dehydrogenase [lipoamide]] kinase isozyme 3, mitochondrial precursor | IPI00123004 | 48 kDa | 0 | 0 | 2 | 2 |
| Sdc3 Syndecan-3 precursor | IPI00135452 | 46 kDa | 0 | 0 | 2 | 2 |

| | | | | | | |
|---|---|---|---|---|---|---|
| Rpl23 60S ribosomal protein L23 | IPI00139780 | 15 kDa | 0 | 0 | 2 | 2 |
| Phyhipl Isoform 1 of Phytanoyl-CoA hydroxylase-interacting protein-like | IPI00224093 | 42 kDa | 0 | 0 | 2 | 2 |
| N28178;LOC100044356 Isoform 2 of Protein KIAA1045 | IPI00377681 (+2) | 41 kDa | 0 | 0 | 2 | 2 |
| Gramd1a Isoform 1 of GRAM domain-containing protein 1A | IPI00124534 (+1) | 81 kDa | 0 | 0 | 2 | 2 |
| Rpl28;OTTMUSG00000021838;LOC100047349 60S ribosomal protein L28 | IPI00222547 (+2) | 16 kDa | 0 | 0 | 2 | 2 |
| D1Ertd53e Protein FAM126B | IPI00226426 (+1) | 59 kDa | 0 | 0 | 2 | 2 |
| 3300001P08Rik Isoform 2 of Cisplatin resistance-associated overexpressed protein | IPI00122418 (+2) | 58 kDa | 0 | 0 | 2 | 2 |
| Sdpr Serum deprivation-response protein | IPI00135660 | 47 kDa | 0 | 0 | 2 | 2 |
| Rab21 Ras-related protein Rab-21 | IPI00337980 | 24 kDa | 0 | 0 | 2 | 2 |
| Rpl23a;ENSMUSG00000063556 60S ribosomal protein L23a | IPI00461456 (+7) | 18 kDa | 0 | 0 | 2 | 2 |
| Impact Protein IMPACT | IPI00319956 | 36 kDa | 0 | 0 | 2 | 2 |
| Elp2 Isoform 1 of Elongator complex protein 2 | IPI00461189 (+1) | 93 kDa | 0 | 0 | 2 | 2 |
| Capn3 Isoform Long of Calpain-3 | IPI00136966 (+3) | 94 kDa | 0 | 0 | 2 | 2 |
| Agl amylo-1,6-glucosidase, 4-alpha-glucanotransferase | IPI00662244 | 174 kDa | 0 | 0 | 2 | 2 |
| Akr1a4 Alcohol dehydrogenase | IPI00466128 (+2) | 37 kDa | 0 | 0 | 2 | 2 |
| Zc3h11a Zinc finger CCCH domain-containing protein 11A | IPI00421162 | 86 kDa | 0 | 0 | 2 | 2 |
| Elp3 Isoform 2 of Elongator complex protein 3 | IPI00224682 (+1) | 64 kDa | 0 | 0 | 2 | 2 |
| Tmem1 Transport protein particle subunit TMEM1 | IPI00408734 (+1) | 142 kDa | 0 | 0 | 2 | 2 |
| Dhrs13 Isoform 1 of Dehydrogenase/reductase SDR family member 13 precursor | IPI00223154 | 41 kDa | 0 | 0 | 2 | 2 |
| Hapln3 Hyaluronan and proteoglycan link protein 3 precursor | IPI00330609 (+1) | 41 kDa | 0 | 0 | 2 | 2 |
| Zfml Isoform 4 of Zinc finger protein 638 | IPI00121264 (+1) | 214 kDa | 0 | 0 | 2 | 2 |
| Arfgef1 ADP-ribosylation factor guanine nucleotide-exchange factor 1 | IPI00626782 (+1) | 209 kDa | 0 | 0 | 2 | 2 |
| Lamc1 Laminin subunit gamma-1 precursor | IPI00400016 | 177 kDa | 0 | 0 | 2 | 2 |
| Lsm14a LSM14 protein homolog A | IPI00172202 | 51 kDa | 0 | 0 | 2 | 2 |
| Xrn2 Isoform 1 of 5'-3' exoribonuclease 2 | IPI00120046 (+1) | 109 kDa | 0 | 0 | 2 | 2 |
| Nfs1 nitrogen fixation gene, yeast homolog 1 | IPI00311072 (+1) | 51 kDa | 0 | 0 | 2 | 2 |
| 2310030N02Rik UPF0551 protein C8orf38 homolog | IPI00341302 | 38 kDa | 0 | 0 | 2 | 2 |
| Itsn1 Isoform 1 of Intersectin-1 | IPI00129356 (+1) | 194 kDa | 0 | 0 | 2 | 2 |
| Serbp1 Isoform 1 of Plasminogen activator inhibitor 1 RNA-binding protein | IPI00471475 (+3) | 45 kDa | 0 | 0 | 2 | 2 |
| Eif2c2 Eukaryotic translation initiation factor 2C 2 | IPI00229988 (+1) | 97 kDa | 0 | 0 | 2 | 2 |
| 4921505C17Rik Isoform 1 of Rapamycin-insensitive companion of mTOR | IPI00399440 (+1) | 192 kDa | 0 | 0 | 2 | 2 |
| Mark1 Serine/threonine-protein kinase MARK1 | IPI00128363 | 89 kDa | 0 | 0 | 2 | 2 |
| Daam2 Isoform 2 of Disheveled-associated activator of morphogenesis 2 | IPI00338570 | 124 kDa | 94 | 41 | 0 | 0 |
| Appl2 DCC-interacting protein 13-beta | IPI00170083 | 74 kDa | 87 | 35 | 0 | 0 |
| Lmtk2 Serine/threonine-protein kinase LMTK2 precursor | IPI00135746 (+1) | 160 kDa | 87 | 29 | 0 | 0 |
| Pex5 Isoform 1 of Peroxisomal targeting signal 1 receptor | IPI00788355 | 71 kDa | 46 | 27 | 0 | 0 |
| Plcd3 1-phosphatidylinositol-4,5-bisphosphate phosphodiesterase delta-3 | IPI00331040 | 89 kDa | 23 | 17 | 0 | 0 |
| Cttnbp2 cortactin binding protein 2 | IPI00672924 (+1) | 179 kDa | 27 | 15 | 0 | 0 |
| Bat3 Large proline-rich protein BAT3 | IPI00130381 (+1) | 121 kDa | 25 | 14 | 0 | 0 |
| Clpx ATP-dependent Clp protease ATP-binding subunit clpX-like, mitochondrial precursor | IPI00119808 (+1) | 69 kDa | 16 | 14 | 0 | 0 |
| Ubr5 ubiquitin protein ligase E3 component n-recognin 5 isoform 1 | IPI00831567 (+2) | 309 kDa | 15 | 14 | 0 | 0 |

| | | | | | | |
|---|---|---|---|---|---|---|
| Zkscan3 10 days neonate skin cDNA, RIKEN full-length enriched library, clone:4732468M21 | IPI00228736 | 63 kDa | 20 | 13 | 0 | 0 |
| Dab2 Isoform p93 of Disabled homolog 2 | IPI00222828 (+1) | 80 kDa | 18 | 13 | 0 | 0 |
| Rcn2 2 days neonate thymus thymic cells cDNA, RIKEN full-length enriched library, clone:E4 | IPI00474959 (+1) | 41 kDa | 21 | 12 | 0 | 0 |
| Nwd1 Isoform 1 of NACHT and WD repeat domain-containing protein 1 | IPI00869414 | 173 kDa | 13 | 12 | 0 | 0 |
| Mtap7d2 Isoform 2 of MAP7 domain-containing protein 2 | IPI00459177 (+1) | 83 kDa | 13 | 11 | 0 | 0 |
| Git1 ARF GTPase-activating protein GIT1 | IPI00470095 (+1) | 85 kDa | 10 | 10 | 0 | 0 |
| Arrb2 Isoform 1 of Beta-arrestin-2 | IPI00130214 (+1) | 46 kDa | 33 | 9 | 0 | 0 |
| Rcn1 Reticulocalbin-1 precursor | IPI00137831 | 38 kDa | 15 | 9 | 0 | 0 |
| 6330569M22Rik Isoform 1 of Protein FAM40A | IPI00223670 (+1) | 96 kDa | 9 | 9 | 0 | 0 |
| Zfp191 Zinc finger protein 24 | IPI00127067 | 42 kDa | 24 | 8 | 0 | 0 |
| Vgf VGF nerve growth factor inducible | IPI00378764 | 68 kDa | 13 | 8 | 0 | 0 |
| Rab11fip3 RAB11 family interacting protein 3 (class II) isoform 1 | IPI00461984 (+1) | 124 kDa | 12 | 8 | 0 | 0 |
| Syngap1 similar to SynGAP-a | IPI00663736 | 142 kDa | 9 | 8 | 0 | 0 |
| Parc p53-associated parkin-like cytoplasmic protein | IPI00624086 | 281 kDa | 8 | 8 | 0 | 0 |
| Appl1 DCC-interacting protein 13-alpha | IPI00170084 | 79 kDa | 13 | 7 | 0 | 0 |
| Hook3 Hook homolog 3 | IPI00353927 | 83 kDa | 10 | 7 | 0 | 0 |
| Fgfr1op2 Isoform 2 of FGFR1 oncogene partner 2 homolog | IPI00110657 (+3) | 25 kDa | 8 | 7 | 0 | 0 |
| 5730469M10Rik Uncharacterized protein C10orf58 homolog precursor | IPI00187272 (+1) | 24 kDa | 7 | 7 | 0 | 0 |
| Ubl4 Ubiquitin-like protein 4A | IPI00471341 | 18 kDa | 7 | 7 | 0 | 0 |
| Fbln5 Fibulin-5 precursor | IPI00323035 | 50 kDa | 10 | 6 | 0 | 0 |
| Atp5h ATP synthase subunit d, mitochondrial | IPI00230507 (+2) | 19 kDa | 10 | 6 | 0 | 0 |
| D330017J20Rik Isoform 1 of Protein FAM40B | IPI00465770 (+1) | 96 kDa | 9 | 6 | 0 | 0 |
| Lrrc7 leucine rich repeat containing 7 | IPI00875695 (+1) | 173 kDa | 6 | 6 | 0 | 0 |
| Gak Isoform 1 of Cyclin G-associated kinase | IPI00420480 (+1) | 144 kDa | 7 | 5 | 0 | 0 |
| Syngr3 Synaptogyrin-3 | IPI00331579 | 25 kDa | 6 | 5 | 0 | 0 |
| Arhgef12 Rho guanine nucleotide exchange factor 12 | IPI00754880 | 172 kDa | 6 | 5 | 0 | 0 |
| Dlg4 Isoform 2 of Disks large homolog 4 | IPI00122094 (+2) | 85 kDa | 6 | 5 | 0 | 0 |
| Dctn3 Dynactin subunit 3 | IPI00331339 | 21 kDa | 6 | 5 | 0 | 0 |
| Ahnak AHNAK nucleoprotein isoform 1 | IPI00553798 | 604 kDa | 5 | 5 | 0 | 0 |
| Aftph Isoform 2 of Aftiphilin | IPI00314905 (+1) | 98 kDa | 5 | 5 | 0 | 0 |
| Akap8 A-kinase anchor protein 8 | IPI00321739 | 76 kDa | 5 | 5 | 0 | 0 |
| 2310022B05Rik Uncharacterized protein C1orf198 homolog | IPI00225267 | 35 kDa | 8 | 4 | 0 | 0 |
| Pde4b phosphodiesterase 4B, cAMP specific | IPI00276327 (+2) | 82 kDa | 8 | 4 | 0 | 0 |
| Arhgef7 Isoform C of Rho guanine nucleotide exchange factor 7 | IPI00230704 (+6) | 89 kDa | 7 | 4 | 0 | 0 |
| Ppfia4 similar to mKIAA0897 protein isoform 1 | IPI00470177 | 130 kDa | 7 | 4 | 0 | 0 |
| Calu Calumenin precursor | IPI00135186 | 37 kDa | 6 | 4 | 0 | 0 |
| 1110014J01Rik Gastric cancer antigen Zg14 homolog | IPI00133594 | 32 kDa | 6 | 4 | 0 | 0 |
| Clpp Putative ATP-dependent Clp protease proteolytic subunit, mitochondrial precursor | IPI00133270 | 30 kDa | 5 | 4 | 0 | 0 |
| Pja2 Isoform 2 of E3 ubiquitin-protein ligase Praja2 | IPI00127272 (+1) | 71 kDa | 5 | 4 | 0 | 0 |
| Ndufaf1 NADH dehydrogenase (ubiquinone) 1 alpha subcomplex, assembly factor 1 | IPI00226687 | 38 kDa | 4 | 4 | 0 | 0 |
| Ptk2b Protein tyrosine kinase 2 beta | IPI00133132 (+2) | 116 kDa | 4 | 4 | 0 | 0 |

| | | | | | | |
|---|---|---|---|---|---|---|
| Stam Mammary gland RCB-0526 Jyg-MC(A) cDNA, RIKEN full-length enriched library, clone:( | IPI00137731 (+1) | 60 kDa | 4 | 4 | 0 | 0 |
| Pebp1 Phosphatidylethanolamine-binding protein 1 | IPI00137730 (+1) | 21 kDa | 4 | 4 | 0 | 0 |
| Tomm70a Mitochondrial precursor proteins import receptor | IPI00377728 (+1) | 68 kDa | 4 | 4 | 0 | 0 |
| Enah Isoform 4 of Protein enabled homolog | IPI00284246 (+2) | 84 kDa | 4 | 4 | 0 | 0 |
| Zfp326 Isoform 1 of Zinc finger protein 326 | IPI00314507 | 65 kDa | 4 | 4 | 0 | 0 |
| Mib1 E3 ubiquitin-protein ligase MIB1 | IPI00330112 (+1) | 110 kDa | 4 | 4 | 0 | 0 |
| Ctnnd1 Isoform 3 of Catenin delta-1 | IPI00316623 (+12) | 103 kDa | 4 | 4 | 0 | 0 |
| Sart1 U4/U6.U5 tri-snRNP-associated protein 1 | IPI00323674 | 91 kDa | 4 | 4 | 0 | 0 |
| Pcsk2 Neuroendocrine convertase 2 precursor | IPI00126390 | 71 kDa | 4 | 4 | 0 | 0 |
| Centa1 centaurin, alpha 1 | IPI00378768 | 43 kDa | 4 | 4 | 0 | 0 |
| Ddef1 Isoform 1 of 130 kDa phosphatidylinositol 4,5-biphosphate-dependent ARF1 GTPase- | IPI00134544 (+3) | 127 kDa | 4 | 4 | 0 | 0 |
| Tubb2b Tubulin beta-2B chain | IPI00109061 | 50 kDa | 749 | 3 | 0 | 0 |
| Spnb1 spectrin beta 1 | IPI00131376 | 268 kDa | 53 | 3 | 0 | 0 |
| Ahcyl1 Putative adenosylhomocysteinase 2 | IPI00162781 | 59 kDa | 47 | 3 | 0 | 0 |
| sp_ALBU_BOVIN | IPIsp_ALBU_BOVIN | 69 kDa | 7 | 3 | 0 | 0 |
| Cul4a Cullin-4A | IPI00321407 | 88 kDa | 5 | 3 | 0 | 0 |
| Eef1b2 Elongation factor 1-beta | IPI00320208 | 25 kDa | 5 | 3 | 0 | 0 |
| Emd Emerin | IPI00114401 (+1) | 29 kDa | 5 | 3 | 0 | 0 |
| Atad3a Isoform 1 of ATPase family AAA domain-containing protein 3 | IPI00126913 | 67 kDa | 5 | 3 | 0 | 0 |
| Cops2 Isoform 1 of COP9 signalosome complex subunit 2 | IPI00120513 (+1) | 52 kDa | 4 | 3 | 0 | 0 |
| Mog Myelin-oligodendrocyte glycoprotein precursor | IPI00126405 (+1) | 28 kDa | 4 | 3 | 0 | 0 |
| ENSMUSG00000047016;LOC100048389 similar to G protein pathway suppressor 1 | IPI00467796 (+1) | 55 kDa | 4 | 3 | 0 | 0 |
| Dkk3 Dickkopf-related protein 3 precursor | IPI00131904 | 38 kDa | 4 | 3 | 0 | 0 |
| ENSMUSG00000045455;Timm8a1;LOC100045062 Mitochondrial import inner membrane tr | IPI00125776 | 11 kDa | 4 | 3 | 0 | 0 |
| Cops8 COP9 signalosome complex subunit 8 | IPI00187407 | 23 kDa | 4 | 3 | 0 | 0 |
| Stx1a Syntaxin-1A | IPI00131618 (+1) | 33 kDa | 4 | 3 | 0 | 0 |
| Pde6d Retinal rod rhodopsin-sensitive cGMP 3',5'-cyclic phosphodiesterase subunit delta | IPI00115190 | 17 kDa | 4 | 3 | 0 | 0 |
| Atp5k ATP synthase subunit e, mitochondrial | IPI00111770 | 8 kDa | 3 | 3 | 0 | 0 |
| Vsnl1 Visinin-like protein 1 | IPI00230418 | 22 kDa | 3 | 3 | 0 | 0 |
| Dpm1 Dolichol-phosphate mannosyltransferase | IPI00115668 | 29 kDa | 3 | 3 | 0 | 0 |
| Rhoa Transforming protein RhoA precursor | IPI00315100 (+1) | 22 kDa | 3 | 3 | 0 | 0 |
| Epha4 Isoform Long of Ephrin type-A receptor 4 precursor | IPI00129198 (+2) | 110 kDa | 3 | 3 | 0 | 0 |
| Ntrk3 Isoform 1 of NT-3 growth factor receptor precursor | IPI00380220 | 93 kDa | 3 | 3 | 0 | 0 |
| Bag4 BAG family molecular chaperone regulator 4 | IPI00109089 | 49 kDa | 3 | 3 | 0 | 0 |
| Ccdc136 similar to KIAA1793 protein isoform 5 | IPI00124212 (+2) | 137 kDa | 3 | 3 | 0 | 0 |
| Atp6v1f Vacuolar proton pump subunit F | IPI00315999 | 13 kDa | 3 | 3 | 0 | 0 |
| Tiprl TIP41-like protein | IPI00221741 | 31 kDa | 3 | 3 | 0 | 0 |
| Sash1 SAM and SH3 domain-containing protein 1 | IPI00338954 | 136 kDa | 3 | 3 | 0 | 0 |
| Ltbp1 latent transforming growth factor beta binding protein 1 isoform LTBP-1S | IPI00352982 (+3) | 153 kDa | 3 | 3 | 0 | 0 |
| Cdh11 Cadherin-11 precursor | IPI00138190 | 88 kDa | 3 | 3 | 0 | 0 |
| Pja1 Isoform 1 of E3 ubiquitin-protein ligase Praja1 | IPI00309237 | 64 kDa | 3 | 3 | 0 | 0 |

| | | | | | | |
|---|---|---|---|---|---|---|
| Itm2b Integral membrane protein 2B | IPI00130758 | 30 kDa | 3 | 3 | 0 | 0 |
| Sec61b Protein transport protein Sec61 subunit beta | IPI00133030 | 10 kDa | 3 | 3 | 0 | 0 |
| Ctsb Cathepsin B precursor | IPI00113517 | 37 kDa | 3 | 3 | 0 | 0 |
| Usp15 Isoform 1 of Ubiquitin carboxyl-terminal hydrolase 15 | IPI00154012 (+2) | 112 kDa | 3 | 3 | 0 | 0 |
| Maged1 Melanoma-associated antigen D1 | IPI00136178 | 86 kDa | 3 | 3 | 0 | 0 |
| Mbp Isoform 6 of Myelin basic protein | IPI00223379 | 17 kDa | 255 | 2 | 0 | 0 |
| Tuba8 Tubulin alpha-8 chain | IPI00311175 | 50 kDa | 194 | 2 | 0 | 0 |
| Tom1l2 Isoform 2 of TOM1-like protein 2 | IPI00648853 | 53 kDa | 141 | 2 | 0 | 0 |
| Krt6a Keratin, type II cytoskeletal 6A | IPI00131368 (+1) | 59 kDa | 25 | 2 | 0 | 0 |
| Ppp1cc Isoform Gamma-1 of Serine/threonine-protein phosphatase PP1-gamma catalytic su | IPI00123862 (+1) | 37 kDa | 18 | 2 | 0 | 0 |
| Lin7c Lin-7 homolog C | IPI00136498 | 22 kDa | 11 | 2 | 0 | 0 |
| Atp2b4 Plasma membrane Ca++ transporting ATPase 4 splice variant a | IPI00463589 (+2) | 129 kDa | 11 | 2 | 0 | 0 |
| Actn2 actinin alpha 2 | IPI00387557 | 104 kDa | 10 | 2 | 0 | 0 |
| Picalm Isoform 1 of Phosphatidylinositol-binding clathrin assembly protein | IPI00264501 (+5) | 72 kDa | 9 | 2 | 0 | 0 |
| Rtn1 reticulon 1 isoform RTN1-C | IPI00459442 | 24 kDa | 8 | 2 | 0 | 0 |
| Afg3l2 AFG3-like protein 2 | IPI00170357 | 90 kDa | 7 | 2 | 0 | 0 |
| Lrch3 Leucine-rich repeat and calponin homology domain-containing protein 3 precursor | IPI00343346 (+1) | 86 kDa | 6 | 2 | 0 | 0 |
| Lin7b Lin-7 homolog B | IPI00136496 | 23 kDa | 6 | 2 | 0 | 0 |
| Cend1 Cell cycle exit and neuronal differentiation protein 1 | IPI00122826 | 15 kDa | 5 | 2 | 0 | 0 |
| Hnrnpf Isoform 1 of Heterogeneous nuclear ribonucleoprotein F | IPI00226073 (+1) | 46 kDa | 5 | 2 | 0 | 0 |
| Rbx1 RING-box protein 1 | IPI00124752 (+2) | 12 kDa | 5 | 2 | 0 | 0 |
| Arf4 ADP-ribosylation factor 4 | IPI00331663 (+1) | 20 kDa | 5 | 2 | 0 | 0 |
| 6330503C03Rik Isoform 2 of Protein FAM131B | IPI00135561 (+1) | 38 kDa | 4 | 2 | 0 | 0 |
| Gstm5 Glutathione S-transferase Mu 5 | IPI00114380 | 27 kDa | 4 | 2 | 0 | 0 |
| Cul4b cullin 4B | IPI00224689 (+1) | 111 kDa | 4 | 2 | 0 | 0 |
| Dnajc5 Dnajc5 protein | IPI00132206 (+3) | 23 kDa | 3 | 2 | 0 | 0 |
| Lasp1 LIM and SH3 domain protein 1 | IPI00125091 (+2) | 30 kDa | 3 | 2 | 0 | 0 |
| Coq5 Ubiquinone biosynthesis methyltransferase COQ5, mitochondrial precursor | IPI00379695 | 37 kDa | 3 | 2 | 0 | 0 |
| Eif4e Eukaryotic translation initiation factor 4E | IPI00119057 (+1) | 25 kDa | 3 | 2 | 0 | 0 |
| Mlf2 Myeloid leukemia factor 2 | IPI00116372 (+1) | 28 kDa | 3 | 2 | 0 | 0 |
| Snph Syntaphilin | IPI00227035 (+2) | 57 kDa | 3 | 2 | 0 | 0 |
| Ptpra Isoform 1 of Receptor-type tyrosine-protein phosphatase alpha precursor | IPI00108685 (+2) | 94 kDa | 3 | 2 | 0 | 0 |
| Pdia3 Protein disulfide-isomerase A3 precursor | IPI00230108 | 57 kDa | 3 | 2 | 0 | 0 |
| Apba1 Adult male brain UNDEFINED_CELL_LINE cDNA, RIKEN full-length enriched library, clc | IPI00223019 | 93 kDa | 3 | 2 | 0 | 0 |
| Nedd8 NEDD8 precursor | IPI00127021 (+1) | 9 kDa | 3 | 2 | 0 | 0 |
| Cacnb4 Isoform 1 of Voltage-dependent L-type calcium channel subunit beta-4 | IPI00131720 (+2) | 55 kDa | 3 | 2 | 0 | 0 |
| Pcbp1 Poly(rC)-binding protein 1 | IPI00128904 | 37 kDa | 3 | 2 | 0 | 0 |
| Gprin3 G protein-regulated inducer of neurite outgrowth 3 | IPI00226416 | 80 kDa | 3 | 2 | 0 | 0 |
| Cacnb1 Isoform 1 of Voltage-dependent L-type calcium channel subunit beta-1 | IPI00321850 (+1) | 65 kDa | 3 | 2 | 0 | 0 |
| Psmd4 Isoform Rpn10A of 26S proteasome non-ATPase regulatory subunit 4 | IPI00128564 (+1) | 41 kDa | 2 | 2 | 0 | 0 |
| Gabarapl2 Gamma-aminobutyric acid receptor-associated protein-like 2 | IPI00309200 (+1) | 14 kDa | 2 | 2 | 0 | 0 |

| | | | | | | |
|---|---|---|---|---|---|---|
| Exoc8 Exocyst complex component 8 | IPI00377299 | 81 kDa | 2 | 2 | 0 | 0 |
| Rpl10a 60S ribosomal protein L10a | IPI00127085 (+1) | 25 kDa | 2 | 2 | 0 | 0 |
| Vamp2 Vesicle-associated membrane protein 2 | IPI00229703 (+1) | 13 kDa | 2 | 2 | 0 | 0 |
| Cirbp Cold-inducible RNA-binding protein | IPI00121073 | 19 kDa | 2 | 2 | 0 | 0 |
| Plekhb1 Isoform 2 of Pleckstrin homology domain-containing family B member 1 | IPI00135982 (+3) | 23 kDa | 2 | 2 | 0 | 0 |
| Psmb6 Proteasome subunit beta type-6 precursor | IPI00119239 | 25 kDa | 2 | 2 | 0 | 0 |
| 2310061I04Rik hypothetical protein LOC69662 | IPI00665637 | 36 kDa | 2 | 2 | 0 | 0 |
| Sar1a Osteoclast-like cell cDNA, RIKEN full-length enriched library, clone:I420011G14 produ | IPI00115644 | 22 kDa | 2 | 2 | 0 | 0 |
| Pde1b Calcium/calmodulin-dependent 3',5'-cyclic nucleotide phosphodiesterase 1B | IPI00118900 | 61 kDa | 2 | 2 | 0 | 0 |
| Arpc3 Actin-related protein 2/3 complex subunit 3 | IPI00124829 | 21 kDa | 2 | 2 | 0 | 0 |
| Lrrfip1 10 days neonate cortex cDNA, RIKEN full-length enriched library, clone:A830097E06 | IPI00403280 | 37 kDa | 2 | 2 | 0 | 0 |
| Exoc3 Exocyst complex component 3 | IPI00454030 | 86 kDa | 2 | 2 | 0 | 0 |
| Gucy1a2;LOC100044212 guanylate cyclase 1, soluble, alpha 2 | IPI00658782 | 82 kDa | 2 | 2 | 0 | 0 |
| Ahi1 Isoform 1 of Jouberin | IPI00170079 (+1) | 120 kDa | 2 | 2 | 0 | 0 |
| Tnpo1 transportin 1 isoform 1 | IPI00221523 (+2) | 102 kDa | 2 | 2 | 0 | 0 |
| Cdc42 Isoform 2 of Cell division control protein 42 homolog precursor | IPI00113849 | 21 kDa | 2 | 2 | 0 | 0 |
| Eps15 Epidermal growth factor receptor substrate 15 | IPI00117454 (+1) | 98 kDa | 2 | 2 | 0 | 0 |
| Reep5 receptor accessory protein 5 | IPI00315463 | 21 kDa | 2 | 2 | 0 | 0 |
| Hip1r Huntingtin-interacting protein 1-related protein | IPI00311726 (+1) | 119 kDa | 2 | 2 | 0 | 0 |
| Ssbp1 Single-stranded DNA-binding protein, mitochondrial precursor | IPI00111877 (+1) | 17 kDa | 2 | 2 | 0 | 0 |
| Park7 Protein DJ-1 | IPI00117264 (+3) | 20 kDa | 2 | 2 | 0 | 0 |
| Ndn Necdin | IPI00108492 | 37 kDa | 2 | 2 | 0 | 0 |
| Calcoco1 Calcium-binding and coiled-coil domain-containing protein 1 | IPI00461826 | 77 kDa | 2 | 2 | 0 | 0 |
| Cope Coatomer subunit epsilon | IPI00130840 | 35 kDa | 2 | 2 | 0 | 0 |
| Gaa Lysosomal alpha-glucosidase precursor | IPI00111960 | 106 kDa | 2 | 2 | 0 | 0 |
| Saps1 SAPS domain family member 1 | IPI00380742 | 95 kDa | 2 | 2 | 0 | 0 |
| Apoe Apolipoprotein E precursor | IPI00323571 | 36 kDa | 2 | 2 | 0 | 0 |
| P4hb 17 days embryo kidney cDNA, RIKEN full-length enriched library, clone:I920160L24 prc | IPI00122815 (+1) | 57 kDa | 2 | 2 | 0 | 0 |
| Pitpna Phosphatidylinositol transfer protein alpha isoform | IPI00230003 | 32 kDa | 2 | 2 | 0 | 0 |
| 2310003C23Rik Protein C20orf11 homolog | IPI00110487 | 27 kDa | 2 | 2 | 0 | 0 |
| Arpc5l Actin-related protein 2/3 complex subunit 5-like protein | IPI00111117 | 17 kDa | 2 | 2 | 0 | 0 |
| Gm672 Uncharacterized protein KIAA0427 homolog | IPI00400197 (+1) | 70 kDa | 2 | 2 | 0 | 0 |
| Tmem33 Transmembrane protein 33 | IPI00133234 (+2) | 28 kDa | 2 | 2 | 0 | 0 |
| Mpst Adult male cecum cDNA, RIKEN full-length enriched library, clone:9130405K21 produc | IPI00604945 (+1) | 33 kDa | 2 | 2 | 0 | 0 |
| Tcf25 Isoform 1 of Transcription factor 25 | IPI00224699 (+5) | 77 kDa | 2 | 2 | 0 | 0 |
| Sort1 Isoform 1 of Sortilin precursor | IPI00420955 (+1) | 91 kDa | 2 | 2 | 0 | 0 |
| Timm10;Timm13 Mitochondrial import inner membrane translocase subunit Tim13 | IPI00134484 | 10 kDa | 2 | 2 | 0 | 0 |
| Vcpip1 Isoform 1 of Deubiquitinating protein VCIP135 | IPI00377609 | 135 kDa | 2 | 2 | 0 | 0 |
| BC033915 hypothetical protein LOC70661 | IPI00453673 (+2) | 151 kDa | 2 | 2 | 0 | 0 |
| Gtf2i Isoform 1 of General transcription factor II-I | IPI00113686 (+6) | 112 kDa | 2 | 2 | 0 | 0 |
| 5730470L24Rik Suppressor of IKK-epsilon protein A | IPI00461173 | 24 kDa | 2 | 2 | 0 | 0 |

| | | | | | | |
|---|---|---|---|---|---|---|
| Hpca Neuron-specific calcium-binding protein hippocalcin | IPI00230310 (+3) | 22 kDa | 2 | 2 | 0 | 0 |
| Ktn1 Isoform 1 of Kinectin | IPI00122559 (+15) | 153 kDa | 2 | 2 | 0 | 0 |
| Zfp364 Zinc finger protein 364 | IPI00177287 | 34 kDa | 2 | 2 | 0 | 0 |
| Nr1i2 Isoform 1 of Nuclear receptor subfamily 1 group I member 2 | IPI00119793 | 50 kDa | 2 | 2 | 0 | 0 |
| Dda1 hypothetical protein LOC66498 | IPI00381086 (+1) | 16 kDa | 2 | 2 | 0 | 0 |
| Sf3b1 Splicing factor 3B subunit 1 | IPI00623284 | 146 kDa | 2 | 2 | 0 | 0 |
| Map6d1 MAP6 domain-containing protein 1 | IPI00816921 | 20 kDa | 2 | 2 | 0 | 0 |
| Mrpl47 39S ribosomal protein L47, mitochondrial precursor | IPI00322422 | 30 kDa | 2 | 2 | 0 | 0 |
| Dnajb12 0 day neonate cerebellum cDNA, RIKEN full-length enriched library, clone:C230062 | IPI00331598 | 42 kDa | 2 | 2 | 0 | 0 |
| Tsc1 Hamartin | IPI00109178 (+2) | 129 kDa | 2 | 2 | 0 | 0 |
| Chgb Secretogranin-1 precursor | IPI00130307 (+1) | 78 kDa | 2 | 2 | 0 | 0 |
| Pum1 Isoform 1 of Pumilio homolog 1 | IPI00330262 (+5) | 127 kDa | 2 | 2 | 0 | 0 |
| Ppp6c Serine/threonine-protein phosphatase 6 | IPI00132965 | 35 kDa | 2 | 2 | 0 | 0 |
| Qk Isoform 7 of Protein quaking | IPI00130483 (+6) | 38 kDa | 2 | 2 | 0 | 0 |
| Jakmip2 98 kDa protein | IPI00474995 (+1) | 98 kDa | 2 | 2 | 0 | 0 |
| 2810405K02Rik Uncharacterized protein C1orf93 homolog | IPI00119094 | 22 kDa | 2 | 2 | 0 | 0 |
| Magi2 Isoform 4 of Membrane-associated guanylate kinase, WW and PDZ domain-containin | IPI00378020 (+3) | 121 kDa | 2 | 2 | 0 | 0 |

| | **MyoVI-GTD** | **MyoVa-MGT** |
|---|---|---|
| Total Proteins | 848 | 1094 |
| Total Spectral Count | 22274 | 23596 |
| Total Unique Peptides | 6976 | 9640 |
| Mean Spectral Count | 26,3 | 21,6 |
| Median Spectral Count | 6 | 7 |
| Mean Unique Peptides | 8,2 | 8,8 |
| Median Unique Peptides | 4 | 5 |
| No. of reverse hits | 0 | 0 |
| % FDR | 0 | 0 |
